# Guide RNA structure design enables combinatorial CRISPRa programs for biosynthetic profiling

**DOI:** 10.1101/2023.11.17.567465

**Authors:** Jason Fontana, David Sparkman-Yager, Ian Faulkner, Ryan Cardiff, Cholpisit Kiattisewee, Aria Walls, Tommy G. Primo, Patrick C. Kinnunen, Hector Garcia Martin, Jesse G. Zalatan, James M. Carothers

## Abstract

Engineering bacterial metabolism to efficiently produce chemicals and materials from multi-step pathways requires optimizing multi-gene expression programs to achieve enzyme balance. CRISPR-Cas transcriptional control systems are emerging as important metabolic engineering tools for programming multi-gene expression regulation. However, poor predictability of guide RNA folding can disrupt enzyme balance through unreliable expression control. We devised a set of computational parameters that can describe guide RNA folding, and we expect them to be broadly applicable across CRISPR-Cas9 systems. Here, we correlate efficacy of modified guide RNAs (scRNAs) for CRISPR activation (CRISPRa) in *E. coli* with a kinetic parameter describing folding rate into the active structure. This parameter also enables forward design of new scRNAs, with no observed failures in our screen. We use CRISPRa target sequences from this set to design a system of three synthetic promoters that can orthogonally activate and tune expression of chosen outputs over a >35-fold dynamic range. Independent activation tuning allows experimental exploration of a three-dimensional expression design space *via* a 64-member combinatorial triple-scRNA library. We apply these CRISPRa programs to two biosynthetic pathways, demonstrating production of valuable pteridine and human milk oligosaccharide products in *E. coli*. Profiling these design spaces indicated expression combinations producing up to 2.3-fold higher titer than that produced by maximal expression. Mapping production can also identify bottlenecks as targets for pathway redesign, improving titer of the oligosaccharide lacto-*N*-tetraose by 6-fold. Aided by computational scRNA efficacy prediction, the combinatorial CRISPRa strategy enables effective optimization of multi-step metabolic pathways. More broadly, the guide RNA design rules uncovered here may enable the routine design of effective multi-guide programs for a wide range of model- and data-driven applications of CRISPR gene regulation in bacterial hosts.

## INTRODUCTION

Synthetic biology and metabolic engineering have great potential for enabling chemical bioproduction from sustainable feedstocks as part of a circular bioeconomy^1–3^. Efficient microbial conversion of simple substrates into valuable chemicals and materials often requires precise expression control across multiple genes to optimize enzyme levels and stoichiometry. Despite recent advances in gene expression technologies, it remains challenging to engineer and optimize multi-step metabolic pathways^4–6^. CRISPR-Cas transcriptional control systems have emerged as promising routes for programming the precise expression of multiple genes, which could accelerate the development of engineered organisms for a wide variety of applications^7–10^. We recently developed an approach for the construction of multi-gene CRISPR transcriptional control programs in bacteria, with activation (CRISPRa) or repression (CRISPRi) functions specified through the regulated expression of multiple guide RNAs (gRNAs)^11,12^. Recent demonstrations of dynamic multi-layer CRISPRa/i gene regulatory network designs in *E. coli*^13,14^ and CRISPR-based metabolic pathway engineering in the soil microbe *Pseudomonas putida*^15–17^ highlight the versatility of these systems for programmable multi-gene control. However, gaps in knowledge and technique continue to prevent the routine design of CRISPRa/i programs capable of quantitatively tuning activated expression from multiple bacterial genes at the same time^9,18^.

Quantitatively tunable multi-gene expression programs are particularly useful for microbial metabolic engineering applications^19^. It is important to identify gene expression programs that minimize enzyme imbalances in multi-gene heterologous pathways and tune endogenous networks to redirect metabolic flux towards the desired output^4,6,20^. Balanced enzyme expression helps minimize bottlenecks, prevent excess metabolic burden, and avoid accumulation of toxic intermediates. Identifying these programs is challenging, in part because we lack tools to systematically explore large, multi-dimensional spaces of gene expression programs. Addressing this challenge with CRISPRa/i systems requires reliable and tunable regulation of gene expression, in turn requiring predictive gRNA design tools for bacterial hosts. Significant progress has been made in gRNA design using folding energetics predictions, cell-based screens, and machine learning, although these methods have been applied primarily for gene editing applications in mammalian cells^21^. General design strategies for tunable CRISPRi with modified gRNAs have been reported for both mammalian and bacterial systems^19,22^. However, many bacterial CRISPRa systems use gRNAs with additional structured elements^11,12,23^, and it is unknown whether design rules for effective gRNA function are generalizable across applications and organisms.

Here, we identified structural properties that enable routine guide RNA design for tunable multi-gene bacterial CRISPRa programs. Our CRISPRa system uses modified single guide RNAs (sgRNAs) that are extended with hairpin sequences, termed scaffold RNAs (scRNAs), to recruit the transcriptional activator SoxS upstream of a promoter^11,12^. This recruitment results in activation of a weak minimal promoter to high expression levels. To identify design variables affecting CRISPRa, we investigated a set of thermodynamic and kinetic guide RNA folding parameters. We found that the largest impact emerged from the size of the energy barrier separating the most stable scRNA structure from the active scRNA structure: this single kinetic parameter accurately predicts about 80% of the variation in CRISPR-activated expression. By comparison, we discovered that commonly-used computational tools for gRNA design cannot consistently identify scRNAs for effective bacterial CRISPRa. We expect that our computational approach could be generalized to identify effective gRNAs for a broad range of CRISPR applications, because the parameters are intrinsic to the RNA sequence.

Starting from highly effective and orthogonal scRNAs, we generated predictable variations in gene activation by truncating scRNA spacer sequences. Using these design strategies, we engineered multi-guide programs that simultaneously direct tunable variations in CRISPRa from multiple promoters independently. We applied a combinatorial set of these CRISPRa programs to drive the design of engineered metabolic pathways producing valuable biopterins and oligosaccharide molecules in *E. coli*. Screening productive variants from these multi-gene programs is a simple method of engineering efficient microbial bioproduction. This approach to biosynthetic profiling enables quantitative tuning of various pathways, and therefore is a versatile approach for a broad range of bioproduction applications. Furthermore, the capacity to reliably implement tunable, multiplexed gene expression will improve the ability to precisely implement perturbations computationally predicted^24,25^ to optimize production strains.

## RESULTS

### scRNA target site sequences have variable effects on gene activation

To build multi-gene CRISPRa programs for metabolic engineering, we need promoters that can be selectively targeted for activation through the expression of a matched, or cognate, scRNA (Figure 1). The rules for effective CRISPRa from bacterial promoters are known to be complex^12^. In particular, the 20 bp scRNA target site must be precisely positioned relative to the transcription start site for effective gene activation. We previously identified a highly-effective promoter (J3) with an appropriately-positioned target site^12^. By altering only the target site sequence of the J3 promoter, we expected to generate orthogonal promoters that retain high levels of gene expression.

**Figure 1.**
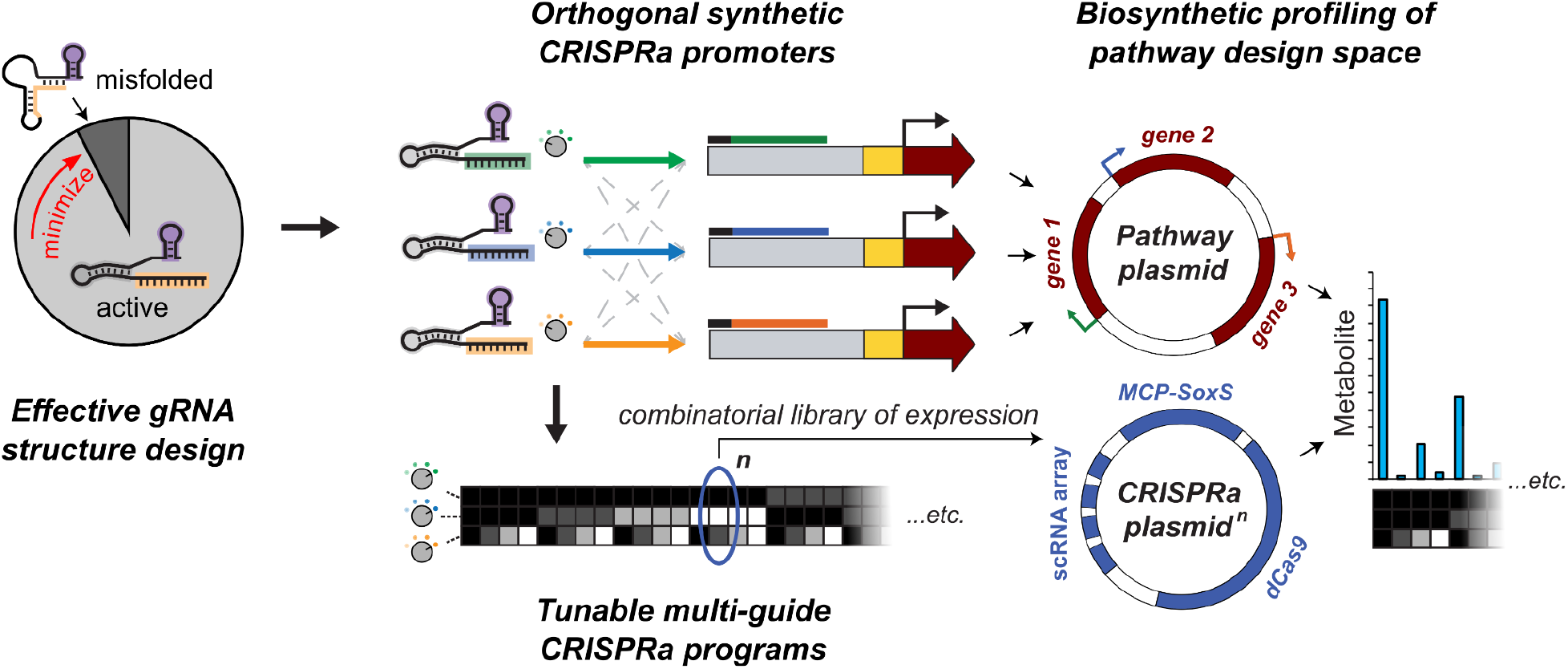
Structure-based guide RNA design and synthetic promoters enable design space mapping with tunable CRISPRa. Computational analysis of scRNA sequence identified a kinetic parameter describing the rate of conversion between the most stable structure and the active structure for CRISPRa, and scRNAs screened using this parameter predictably activated bacterial expression from a set of synthetic promoters. Tuning the activation of these promoters by truncating their scRNA spacer sequences—and again computationally verifying their efficacy—allows combinations of activation level at each promoter. The promoters can be paired with chosen output ORFs, including metabolic pathways. This method of controlling pathway gene expression allows for profiling of pathway design spaces for metabolic engineering using a combinatorial library of CRISPR-activated expression levels.

We modified the J3 target site sequence to generate 14 additional synthetic promoters with fully randomized target sites, each paired with its cognate scRNA (Figure 2a). Targeting the CRISPRa complex in this way to each of the 15 promoters activated expression of a downstream fluorescent reporter gene (Figure 2b). All of the promoter variants showed measurable activation compared to the off-target scRNA control, but there was significant variability over a 3-fold range in expression levels (Figure 2b). Consistent with previous findings^12^, these results suggest that the target site sequence identity can have unexpectedly large effects on gene activation.

**Figure 2.**
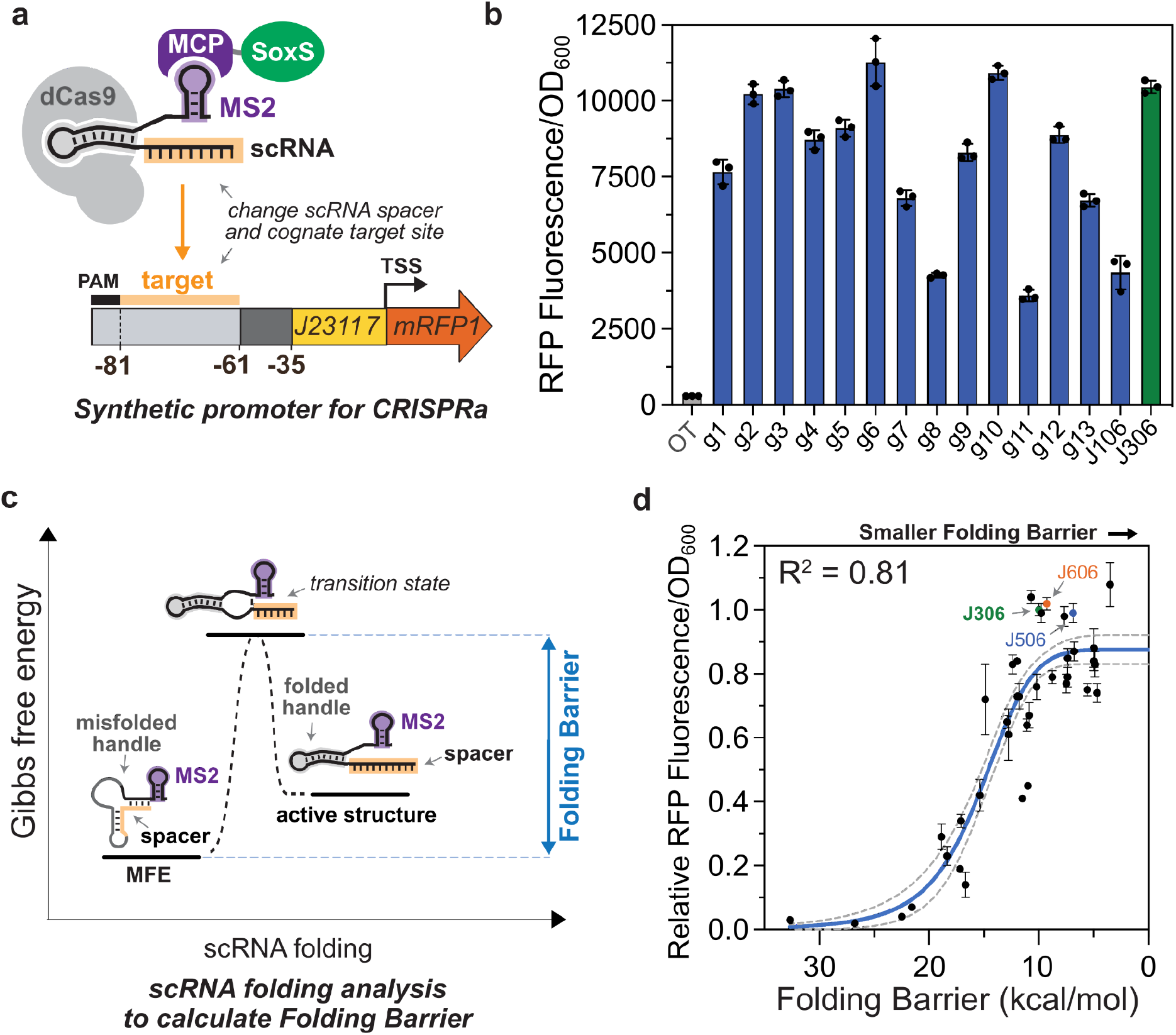
CRISPRa is sensitive to scRNA target sequence. **a** Experimental system for testing the role of scRNA target site sequence on CRISPRa activity. Orthogonal 20 bp target sequences (Supplementary Table 3) were selected at random from the human genome. These sequences replaced the J306 target sequence in the previously described J3 promoter^12^, and the cognate scRNAs contained the complementary spacer sequences. No other part of the promoter was altered. **b** CRISPR-activated RFP expression from each promoter variant. In the presence of the cognate scRNA, sequence-dependent expression variation was measured across the set. Blue bars (g1-J106) represent the Fluorescence/OD_600_ of strains harboring each synthetic promoter and the cognate scRNA. The green bar represents the Fluorescence/OD_600_ of the J3 promoter with its cognate J306 scRNA. The grey bar (OT) represents the baseline expression of the J3 promoter, obtained by expressing an off-target scRNA (J206). **c** Folding Barrier (FB) was identified as a critical parameter determining CRISPR-activated expression. Additional kinetic and thermodynamic parameters are described in Supplementary Figure 2 and Supplementary Methods. Folding Barrier can be calculated as the height of the energy barrier separating the minimum free energy (MFE) secondary structure of a scRNA from the secondary structure of that scRNA with the dCas9-binding handle, MS2 hairpin and spacer folded into the active conformations. **d** Folding Barrier predicts the CRISPR-activated expression of synthetic promoters based on the sequence of their cognate scRNA. In addition to the 15 promoters from panel **b**, 24 new synthetic promoters were designed to test expression dependence on FB. These target site sequences are paired with cognate scRNAs with FBs ranging from 4.7 kcal/mol to 32.7 kcal/mol (Supplementary Table 3). The *y*-axis values represent Fluorescence/OD_600_ of strains harboring each promoter variant and expressing the cognate scRNA, relative to the Fluorescence/OD_600_ of the J3 promoter and the J306 scRNA (green). Blue and red dots respectively indicate the values of the strains expressing the J506 and J606 scRNAs targeting their cognate promoters (Figure 3). The blue line represents a Hill function fit to the data, and the grey dotted lines represent the 95% confidence interval for the fit. R^2^ represents the coefficient of determination for the fit. Values in panels **b** and **d** represent the average ± standard deviation calculated from *n* = 3 biologically independent samples. Source data for **b** and **d** are provided as a Source Data file.

### The kinetic folding barrier predicts scRNA activity for CRISPRa

Variable activation from the orthogonal synthetic promoters could occur if the corresponding 20 base scRNA spacer sequences have different effects on folding. Changes to the spacer sequence could lead to scRNA misfolding that disrupts binding to dCas9, recruitment of the SoxS activator, or binding to DNA. We reasoned that the kinetic and thermodynamic properties associated with the conversion of a misfolded scRNA into the correctly-folded structure could be important determinants of CRISPRa activity. Scaffold RNAs could be more effective in a kinetic sense if they readily transition to the correctly-folded state, or could be more effective in a thermodynamic sense if they are more likely to occupy the correctly-folded state.

To test these possibilities, we developed two coarse-grained parameters that describe the energetics of scRNA folding: Folding Barrier to capture kinetic properties and Folding Energy to capture thermodynamic properties (Figure 2c, Supplementary Figure 2a). We defined the Folding Energy as the free energy difference between the most stable scRNA structure (Minimum Free Energy, or MFE) and the correctly-folded, CRISPR-active structure. The Folding Energy is large when the correctly-folded structure is less stable than the MFE, and approaches zero as the correctly-folded structure increases in stability. The Folding Barrier is the height of the activation energy barrier separating the MFE structure from the correctly-folded structure. When the MFE structure can easily overcome this barrier and rearrange into the correctly-folded structure, the Folding Barrier is low. The correctly-folded structure was defined as the conformation in which the spacer is unstructured and the Cas9-binding handle adopts the fold observed in the crystal structure of the Cas9-sgRNA-DNA complex^26^. Energetic parameters were calculated using custom algorithms that apply programs in the ViennaRNA folding package^27^ (see Methods).

To probe the relationships between our calculated parameters and CRISPRa function, we experimentally tested a set of 39 scRNA-promoter pairs. This set includes the original J3 sequence, the 14 randomly selected targets described above, and 24 additional scRNAs designed to have Folding Barriers ranging from 5 to 35 kcal/mol (Supplementary Methods). High levels of CRISPR-activated expression correlated with smaller Folding Energies (r_s_ = 0.7) and lower Folding Barriers (r_s_ = 0.8) (Figure 2d, Supplementary Figure 3a). Consistently, the MFE structures of the highest-activation scRNAs in our set closely resembled the active scRNA conformations, whereas the least effective scRNAs misfolded extensively (Supplementary Figure 6). Interestingly, we found that Folding Barrier alone may be sufficient for identifying highly effective scRNAs. The most effective scRNA in our 39-member set had the smallest Folding Barrier. In contrast, three of the worst-performing scRNAs, which generated 95% less gene activation than the J306 scRNA, had the largest Folding Barriers in the set. We also considered other thermodynamic and kinetic parameters for use in predicting scRNA folding, but found that Folding Barrier was the most effective predictor of CRISPRa function, with Folding Energy and Net Binding Energy providing limited additional predictive power for low-FB scRNAs. (Supplementary Figures 2 & 3, Supplementary Methods).

Our data suggest that Folding Barrier analysis could be used to drive the design of scRNAs with a lower chance of weak activity. Out of the 24 rationally-designed scRNAs, the 15 scRNAs with the lowest Folding Barrier all yielded effective CRISPRa (at least 50% of J306 output, or about 18-fold activation), and their CRISPR-activated expression levels showed less variability than those of the 15 randomly-designed scRNAs (Coefficient of variation = 12% vs. 31% for the random set) (Supplementary Figure 4). We observed in our promoter set that high-performing scRNAs tended to have Folding Barriers ≤10 kcal/mol, and all defective scRNAs (<50% of J306 activation) were >10 kcal/mol. Therefore, a Folding Barrier threshold of <10 kcal/mol could provide a useful computational screening metric for rapid development of novel scRNAs.

To further evaluate this new kinetic parameter as a screening tool to design highly effective scRNAs, we compared Folding Barrier with pre-existing models currently in wide use for gRNA design. A common approach to analyze gRNAs involves calculating the free energy of binding a correctly-folded gRNA to its target DNA^28,29^ (termed Binding Energy in Supplementary Figure 2a). In this approach, gRNAs with more negative Binding Energies have unstructured spacer sequences that should favor the DNA-bound state, and should therefore be more active. In our study, however, the scRNAs with the lowest Binding Energy included a significant fraction of defective scRNAs (33%), suggesting that Binding Energy is not sufficient to account for CRISPRa functionality (Supplementary Figure 3, Supplementary Figure 4a). In contrast, the Folding Barrier metric correctly predicts these failures within the low-Binding-Energy set: defective scRNAs had relatively high Folding Barriers averaging 17.6 kcal/mol. Effective (≥50% of J306) scRNAs in this set had an average Folding Barrier of 9.3 kcal/mol, further supporting the use of a Folding Barrier threshold to screen functional scRNAs.

Several machine learning models have also been developed to predict gRNA activity^21,30–35^. These models were trained with supervised learning to extract gRNA design rules from large gene editing datasets and are widely used to aid the selection of gRNA target sites. Among the models we tested, none yielded predictions strongly correlated with observed CRISPR-activated expression from the scRNAs in our set. For example, the widely used Azimuth, Doench ‘16^21^, and Moreno-Mateos^30^ tools had correlation coefficients (r_s_) of 0.22, 0.02, and 0.09, respectively, and incorrectly selected several defective guides as the best (Supplementary Figures 3 and 4). The top 15 scRNAs predicted by these tools contained both defective scRNAs (with consistently higher Folding Barriers, e.g. 21.6 kcal/mol average using Azimuth) and effective ones (7.3 kcal/mol average using Azimuth). Differences between gRNA-directed editing and scRNA-directed activation may account for the poor performance of these models in this application. A machine learning model trained on scRNAs used in bacteria could potentially be effective, but generating large enough bacterial CRISPRa datasets for such a model to account for the stringent target site requirements^12^ might be impractical. Given the predictive success and ease of calculation of the Folding Barrier, we proceeded with this kinetic parameter as a strategy to rapidly design highly effective scRNAs for bacterial CRISPRa.

### Tunable CRISPRa expression from orthogonal synthetic promoters

By forward engineering scRNAs through computational folding design, our tools provide an avenue for developing synthetic promoters driving high levels of CRISPR-activated expression. To be useful for programming combinatorial variations in multi-gene expression, as in a metabolic engineering application, two additional capabilities are needed. First, the synthetic promoters must exhibit orthogonality with no cross-activation from other non-cognate scRNAs expressed in the cell. Second, a strategy is needed to tune expression levels from each of the promoters by independently modulating CRISPRa activity at each site. In this section, we show that promoter orthogonality is readily obtainable and that 5’ spacer sequence truncations enable quantitative and independent tuning of CRISPRa levels.

To construct three sequence-orthogonal synthetic promoters, we selected three high-performing scRNAs from the set identified through folding design. Because most randomly selected 20 base sequences will be orthogonal, we did not apply any explicit filters for orthogonality to select these sequences. The sequences included two new scRNAs, termed J506 and J606, and the previously-described J306 scRNA with its cognate J3 promoter. All three scRNAs have low Folding Barriers (≤10 kcal/mol), consistent with the threshold criterion for effective scRNA selection. To construct cognate synthetic promoters for J506 and J606, termed J5 and J6, we inserted each target site at the optimal position 81 bases upstream of the transcription start site (Figure 3a). To minimize repeating sequence elements between the promoters, we inserted distinct sequences in the intervening 26 bases between the target site and the minimal promoter, using sequences previously screened to permit high CRISPRa activity in this context^12^. We also randomized about 120 bases upstream of the target site PAM in J5 and J6, without introducing additional dCas9 PAMs. From the new J5 and J6 promoters, we observe high levels of CRISPR-activated expression, similar to the expression level from the J3 promoter (Figure 3b). To confirm orthogonality of J3/J5/J6, we measured the response of each promoter paired with either non-cognate scRNA and observed no activation (Figure 3B).

**Figure 3.**
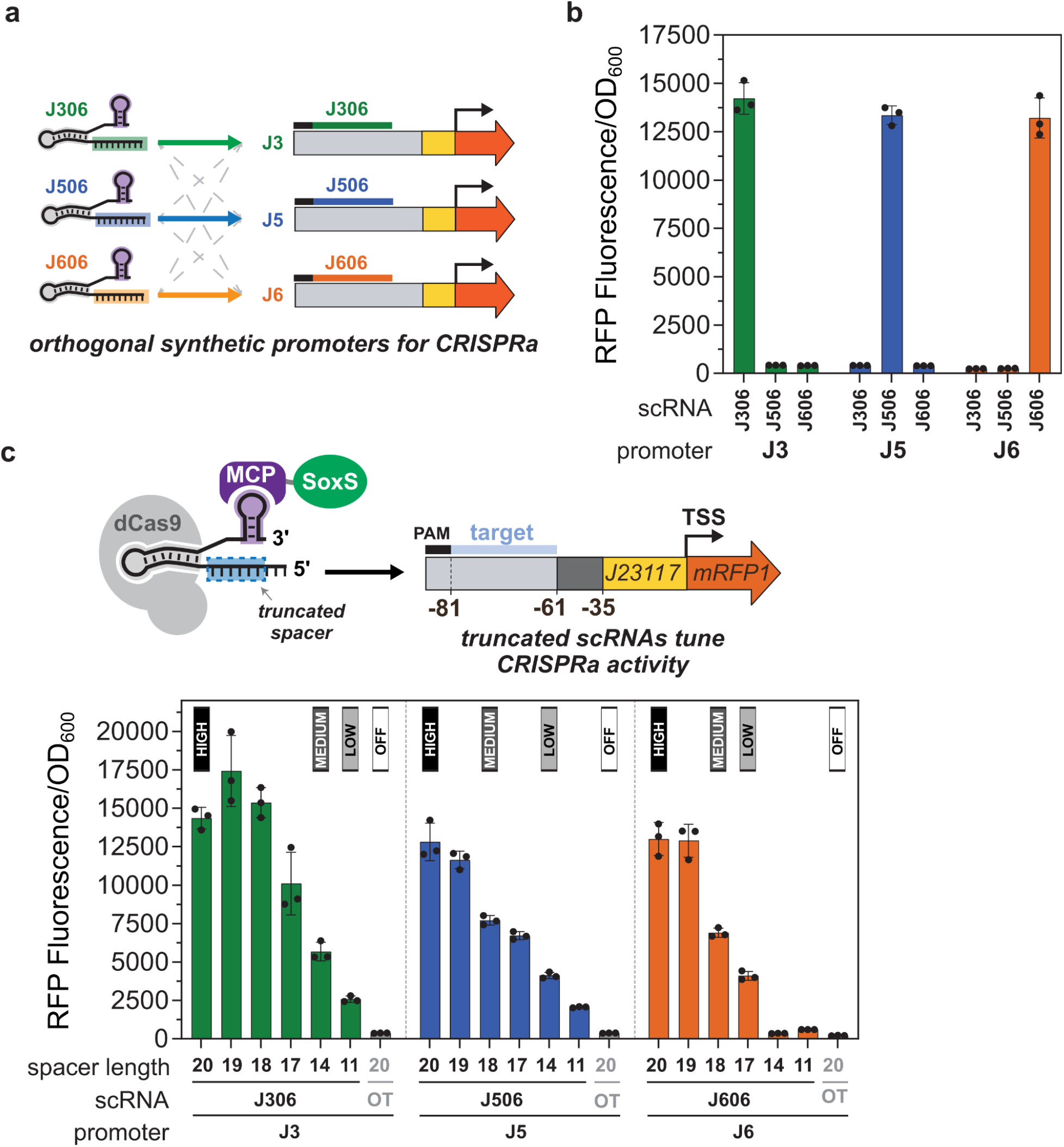
CRISPR activation of orthogonal synthetic promoters can be tuned using truncated scRNAs. **a** Orthogonal CRISPR activation was achieved for the J3, J5, and J6 synthetic promoters by the sequence orthogonality of their cognate scRNAs (J306, J506, J606, respectively). While J3 was previously described^12^, J5 and J6 were selected from our set of 38 synthetic promoters (Figure 2d) because they generated similar CRISPR-activated expression levels as J3. **b** Synthetic promoters for CRISPRa can be selectively activated by expressing their cognate scRNAs. Bars represent the Fluorescence/OD_600_ of strains harboring the J3, J5, or J6 promoters and expressing the cognate or non-cognate scRNAs. **c** CRISPR-activated expression from the J3, J5, and J6 promoters can be tuned with truncated scRNAs by removing nucleotides from the 5’ end of the spacer. Bars represent the Fluorescence/OD_600_ of strains harboring J3, J5, or J6 and expressing the cognate scRNAs truncated to 19, 18, 17, 14, and 11 bases. Grey bars represent the baseline expression of the promoters, obtained from strains expressing an off-target scRNA (J206). Labels above bars indicate the spacer length chosen to encode high, medium, low and off expression levels in the combinatorial scRNA library (Figure 4). Values in panels **b** and **c** represent the average ± standard deviation calculated from *n* = 3 biologically independent samples. Source data for **b** and **d** are provided as a Source Data file. The full sequences of the J3, J5, and J6 promoters are described in Supplementary Methods.

To generate independently tunable expression from our orthogonal CRISPRa promoters, we considered multiple strategies. Several approaches have been described, generally either by modulating gRNA expression level or by direct modification of gRNA sequence. For example, CRISPRi or CRISPRa activity can be tuned using different strengths of constitutive promoters to drive gRNA expression^23,36^. Alternatively, introducing mismatches in the gRNA spacer sequence can modulate CRISPRi gene repression^37–40^, and truncating the gRNA target sequence from the 5’ end has also been shown to reduce CRISPRi activity^37^. Here, we reasoned that truncation-based tuning would yield a more predictable response than spacer mismatches, and would allow us to keep the same constitutive promoter strength expressing each scRNA. This approach simplifies cloning and decreases the risk of dCas9 binding competition effects^41,42^.

We screened J3-, J5-, and J6-targeted scRNAs truncated 1–9 bases from the 5’ end to identify guides that encode discrete intermediate levels of CRISPR-activated gene expression. Across all three promoters, scRNA spacer truncation gradually reduced CRISPR-activated expression (Figure 3c), and from those functions we selected high, medium, and low activation levels. The folding parameters predict similarly high efficacy for all truncations (Folding Barrier ≤10 kcal/mol), while the Net Binding Energy generally becomes less favorable with truncation (Supplementary Table 6). This effect is consistent with the smaller number of RNA bases available to pair with the DNA target, and loosely correlates with output activation (Supplementary Figure 7). Specifically, the full-length J306 scRNA with a 20 base spacer generated 38-fold activation, and truncated scRNAs with 17, 14, or 11 base spacers tuned CRISPRa to 27-fold, 15-fold, and 7-fold activation, respectively. For the J506 and J606 scRNA truncations, the expected monotonically decreasing trends were observed, although the precise truncations to achieve similar activation levels were not the same (Figure 3c). In particular, the J606 scRNA was more sensitive to truncation than J306 and J506. For instance, the 14-base J606 truncation activated gene expression by only 2-fold, while the 14-base J306 and J506 scRNAs activated their promoters by 15-fold and 11-fold respectively. Consistent with previous work investigating DNA-level sequence context effects on CRISPRa^43^, sequences adjacent to the spacer targets in the J3/J5/J6 promoters might affect truncation response. Even if the energetic parameters here do not quantitatively explain the sensitivity of each promoter’s truncation response (Supplementary Figure 7), they generally reflect the rank order of the tuned outputs (*R_s_* = 0.83 for J306, *R_s_* = 1 for J506, *R_s_* = 0.94 for J606).

Interestingly, the J306 scRNA with a 19 base spacer generated higher activation than the 20 base spacer (46-fold vs. 38-fold) even though the Net Binding Energy for the 20 base spacer (−32.3) was similar to that of the 19 base spacer (−31.4). Taken together, the energetic parameters do not indicate impaired folding of the 20 base spacer or any other indication that the 19 base spacer should perform better for CRISPRa. It is possible that spacer truncations could affect transcription of the scRNA itself or could introduce scRNA folding characteristics not captured by our screening parameters. For practical applications, however, we can empirically choose the appropriate scRNA spacer length from within the truncation datasets to obtain tunable high, medium, or low activation from each of the three promoters.

### Combinatorial CRISPRa library enables tuning of multi-gene expression programs

Encoding expression levels directly in multi-scRNA programs creates a straightforward way to implement combinatorial variations in the expression of multi-gene systems. Genes of interest can be cloned under the control of a set of synthetic CRISPRa promoters and tuned by simply changing the identity of the scRNAs transcribed in the cell. For example, driving the expression of three genes with the J3, J5, and J6 promoters and expressing a combination of a J306 scRNA with an 11 base spacer, J506 with a 20 base spacer and J606 with an 18 base spacer would result in low, high, and medium expression of the corresponding genes. By extending such a strategy to encompass all possible combinations of truncated J306, J506 and J606 scRNAs, we can rapidly explore large combinatorial spaces of gene expression under the control of CRISPRa promoters (Figure 4a).

**Figure 4.**
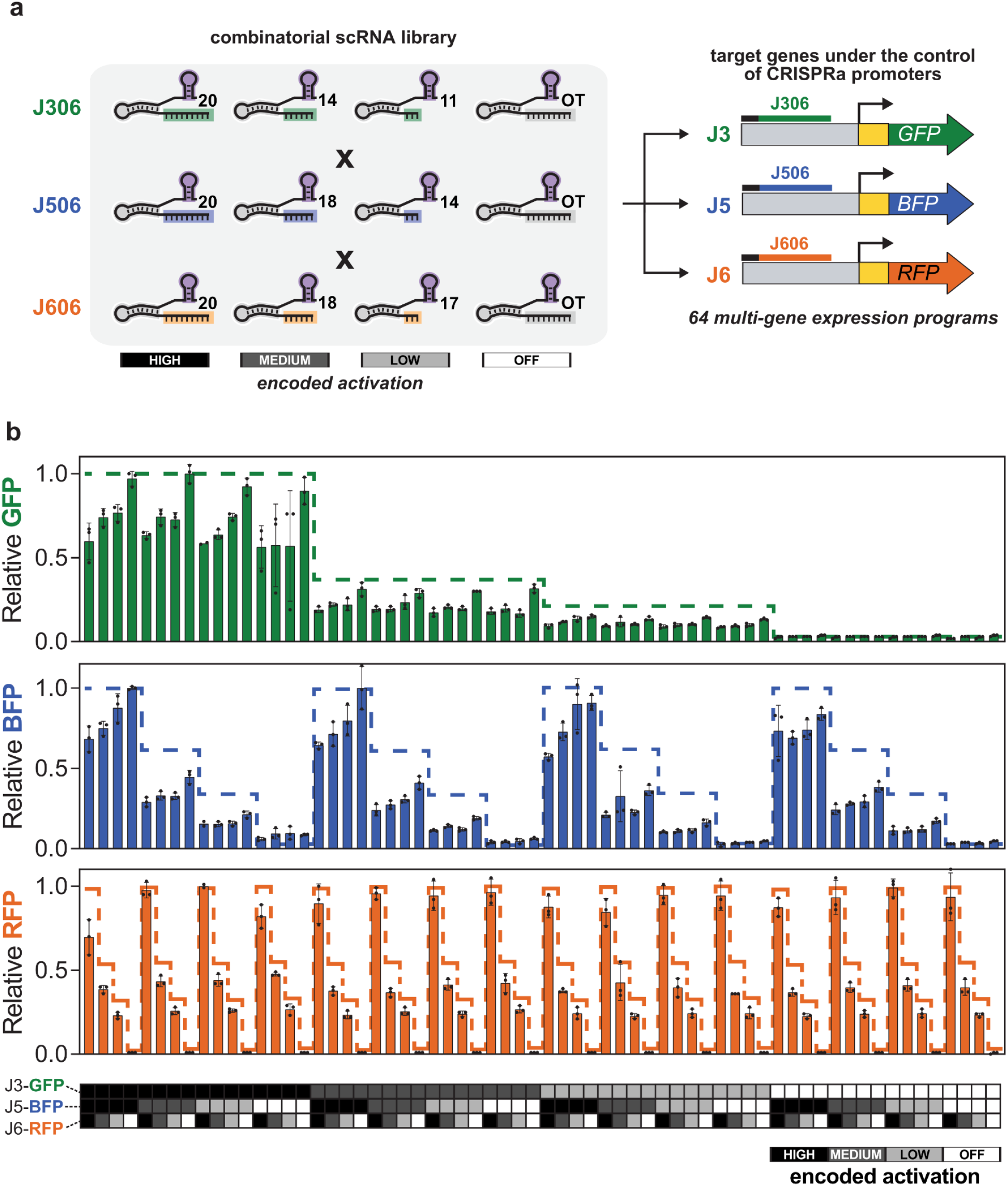
Multi-gene expression can be rapidly tuned using combinatorial CRISPRa programs. **a** Combinatorial library encoding all combinations of four CRISPR-activated expression levels across three genes. The library expresses three scRNAs (variants of J306, J506, and J606). Each scRNA is present in the library in three truncation variants to generate high, medium and low levels of expression of their target promoters (J3, J5, and J6, respectively). In addition to the three truncation variants, the library contains strains with an off-target scRNA in place of each of the J306, J506, and J606 scRNAs to encode a condition in which the target promoter remains unactivated. The lengths of the J306 scRNA variants are 20, 14, and 11 bases. The J506 scRNA variants are 20, 18, and 14 bases. The J606 scRNA variants are 20, 18, and 17 bases. **b** Use of the combinatorial scRNA library to specify the expression of multiple genes independently. Each member of the combinatorial scRNA library was delivered to a strain harboring a plasmid expressing J3-*gfp*, J5-*bfp*, and J6-*rfp* reporters, generating 64 strains expressing different combinations of the three fluorescent proteins. Bars represent the flow cytometry median of GFP, BFP, and RFP from each strain, normalized to the maximum level across the experiment. The heatmap table below the plot indicates the encoded promoter expression for each strain, as described on the bottom right. Dashed lines represent the Relative Fluorescence/OD_600_ of strains harboring only one of the three fluorescent reporters and only the cognate scRNA (see Supplementary Table 2 for plasmids and Supplementary Figure 10 for variation in single-channel expression), again normalized to the maximum value. Bars in panel **b** represent the average ± standard deviation calculated from *n* = 3 biologically independent samples. Source data are provided as a Source Data file. The sequence of each scRNA in the combinatorial library can be found in Supplementary Table 3. The sequence of the reporter plasmid expressing J3-*gfp*, J5-*bfp*, and J6-*rfp* is described in the Supplementary Methods.

We demonstrate the immediate utility of this design strategy by creating a set of genetic tools for combinatorial gene expression profiling. We constructed a library of multi-scRNA “program” plasmids that encode the expression levels from the set of synthetic CRISPRa promoters, which control a set of desired genes on an “output” plasmid. Three-gene combinatorial expression profiling is then enabled by simply combining an output plasmid with each member of the program library (Figure 1), allowing the same scRNA library to be used for arbitrary outputs. We constructed a full library of scRNA plasmid variants to encode all possible combinations of high, medium, low (Figure 3c) and basal expression of three target genes. Basal expression from each of the targeted promoters was minimal and resulted from an off-target scRNA. Together, the library is composed of 64 plasmids (4^3^) that can be combined with any construct containing genes driven by the J3, J5, and J6 synthetic promoters, resulting in strains encoding 64 different combinations of multi-gene expression.

As an initial validation of our strategy, we tested the combinatorial multi-scRNA library using fluorescent reporter expression. We delivered each of the 64 constructs from the library to an *E. coli* strain containing GFP, BFP, and RFP reporters under the control of the J3, J5, and J6 promoters, respectively. The resulting strains displayed every combination of high, medium, low, and basal expression for the three reporters. Across this set, the strains displayed variations in relative expression levels consistent with the multi-scRNA programs they contained (Figure 4b). However, we also observed that tuning one gene could affect expression of the other genes. First, we found that total expression was reduced by 30-40% when high activation was simultaneously encoded for all three reporters, suggesting that high heterologous gene expression is limited by host expression capacity. Although these effects will vary with different target genes and ribosome binding site strengths, they indicate that maximal expression of multiple genes in a pathway can have unintended consequences that may result in suboptimal behavior. Second, we observed that high expression specifically of RFP had a deleterious effect on GFP and BFP levels (Supplementary Figure 8). It is well-established that expression burden, metabolic burden, or toxicity can have effects on gene expression levels that are difficult to predict^44,45^. Our findings underscore the importance of systematically exploring the combinatorial design spaces of multi-gene expression programs to optimize engineered systems. Using this strategy, we applied our CRISPRa tools to build combinatorial expression programs to optimize flux through two engineered metabolic pathways.

### Biosynthetic profiling of an engineered tetrahydrobiopterin pathway with combinatorial CRISPRa programs

To determine if combinatorial optimization would be effective for metabolic engineering, we applied our CRISPRa promoters and library approach to regulate tetrahydrobiopterin (BH4) biosynthesis. BH4 is a central cofactor in aromatic amino acid metabolism and a treatment for life-threatening metabolic disorders, including a form of phenylketonuria^46^. It can be produced from a three-enzyme pathway^47–49^ using the *E. coli gtpch* and *M. alpina ptps* and *sr* genes, as described previously^15^. Production can be monitored with a fluorimetric assay^47–49^, providing a convenient model system for combinatorial screening. We placed codon-optimized *gtpch*, *ptps,* and *sr* genes in a BH4 pathway plasmid with enzyme expression controlled by the J3, J5, and J6 synthetic promoters, respectively (Figure 5a and b). Co-transforming the BH4 pathway plasmid into *E. coli* with each member of our combinatorial multi-scRNA library resulted in 64 new strains, each encoding a different combination of high, medium, low, and basal expression of the BH4 pathway enzymes. We monitored biosynthetic flux through this pathway by measuring the absorbance at 340 nm, which reports on the spontaneous BH4 oxidation products dihydrobiopterin (BH2) and biopterin^15^.

**Figure 5.**
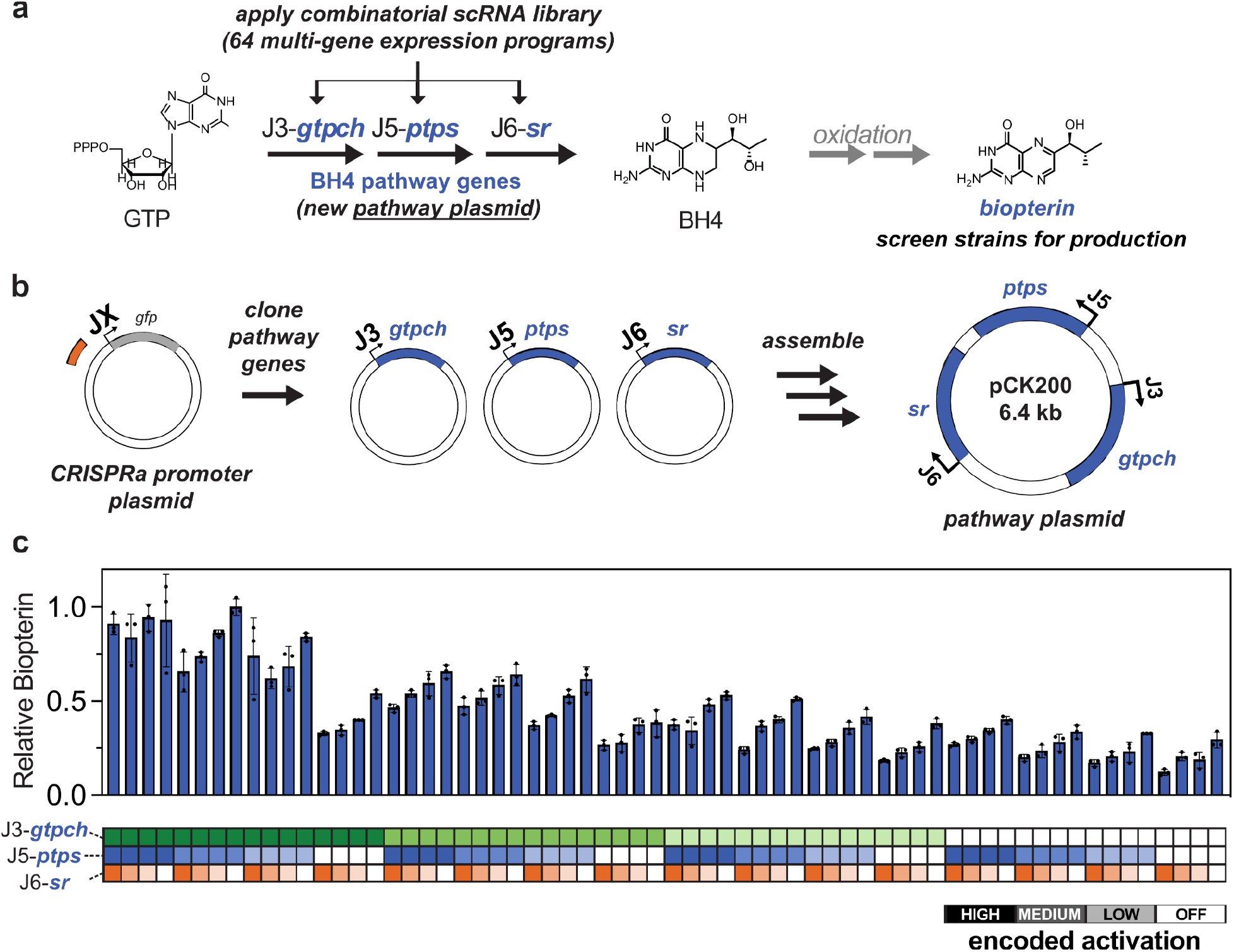
Combinatorial CRISPRa programs can be applied to tune biosynthetic pathways. **a** Tetrahydrobiopterin (BH4) production was tuned by delivering the combinatorial scRNA library to an *E. coli* strain harboring a BH4 pathway plasmid. BH4 is synthesized from GTP by expressing the *gtpch* gene from *E. coli* and the *ptps* and *sr* genes from *M. alpina*. BH4 then undergoes two oxidative decomposition steps yielding dihydrobiopterin (BH2, not shown) and biopterin. The BH4 pathway plasmid was constructed by placing the *gtpch*, *ptps* and *sr* genes under control of the J3, J5, and J6 promoters, respectively. **b** Tuning gene expression in new biosynthetic pathways only requires constructing a new pathway plasmid. The new plasmid is then cotransformed with the same scRNA library from Figure 4. **c** Combinatorial tuning of BH4 pathway reveals that *gtpch* activity is limiting and that the *sr* gene is expressed in excess. Bars represent the relative biopterin of each strain in the combinatorial library harboring the BH4 pathway plasmid. Relative biopterin is measured through OD_600_-normalized fluorescence (excitation: 340 nm, emission: 440 nm), which corresponds to BH4 and its oxidized derivative BH2. Values are relative to the strain with the highest Fluorescence/OD_600_ within the set. The *x*-axis heatmap is color coded to indicate the encoded promoter expression for each strain, as described on the bottom right. Values in panel **c** represent the average ± standard deviation calculated from *n* = 3 biologically independent samples. Source data are provided as a Source Data file. The sequence of the pathway plasmid containing J3-*gtpch*, J5-*ptps* and J6-*sr* is described in the Supplementary Methods.

We observed the highest BH4 production in strains with high expression of the first enzyme in the pathway, GTPCH, indicating that *gtpch* expression is a sensitive control point in this system (Figure 5c). Reducing J3-*gtpch* activation from high to low decreased production by an average of 54%. Changes in expression of the second enzyme, PTPS, had relatively little impact on production across the whole set of combinatorial programs (J5-*ptps* high to low reduced production by an average of 20%), except for conditions in which its expression was basal (high to basal reduced production by an average of 46%). Interestingly, basal expression of the SR enzyme was not only sufficient for BH4 production, but increasing its expression led to reduction in product titers. For example, increasing J6-*sr* activation from off-target to high reduced production by an average of 36%. This reduction was widespread and consistent, with 13 out of 16 J6-high strains producing significantly less BH4 than their off-target counterparts. Previous kinetic characterization of SR renders this result unsurprising^50^, because even basal SR expression provides a vast excess of activity relative to the flux delivered by the upstream pathway. Additional SR beyond the basal level presumably only contributes additional expression burden without increasing overall pathway flux. Taken together, these results identify effective enzyme levels for BH4 biosynthesis and highlight that maximal expression of all enzymes is not optimal.

### Applying biosynthetic profiling for efficient production of a human milk oligosaccharide

We next applied our CRISPRa system to perform combinatorial expression analysis of a multi-gene pathway for producing the valuable oligosaccharide lacto-*N*-tetraose (LNT)^51,52^. Human milk oligosaccharides (HMOs) are major components of human milk^53^ with substantial effects on infant immune development^54^, microbiome establishment^55,56^, anti-inflammation^57,58^, and more^59^. Microbial production may provide routes to obtain scalable quantities of HMOs for research, nutrition, and therapeutic applications that are otherwise difficult to obtain using traditional chemical synthesis^60,61^. LNT is a highly abundant HMO, a valuable formula additive, and a core structure of several other structurally-diverse HMOs^60,62^.

A three gene pathway consisting of the LacY lactose permease and two heterologous enzymes, LgtA^63^ and WbgO^64^, can produce LNT in *E. coli*^51,52^ (Figure 6a). Starting from a lactose feedstock supplied in the media, *E. coli* LacY imports the lactose into the cell, where LgtA, a β-1,3-*N*-acetylglucosaminyltransferase from *Neisseria meningitidis*, produces the intermediate metabolite lacto-*N*-triose II (LNT II) using the hexose sugar from endogenous UDP-*N*-acetylglucosamine. WbgO, a β-1,3-galactosyltransferase from *E. coli* O55:H7, then produces LNT using LNT II and endogenous UDP-galactose. Knocking out endogenous β-galactosidase activity (*lacZ*) is also necessary to prevent cleavage of the lactose feedstock into its constituent monosaccharides glucose and galactose, which would divert flux away from LNT biosynthesis and toward glycolysis^51,60,65–68^.

**Figure 6.**
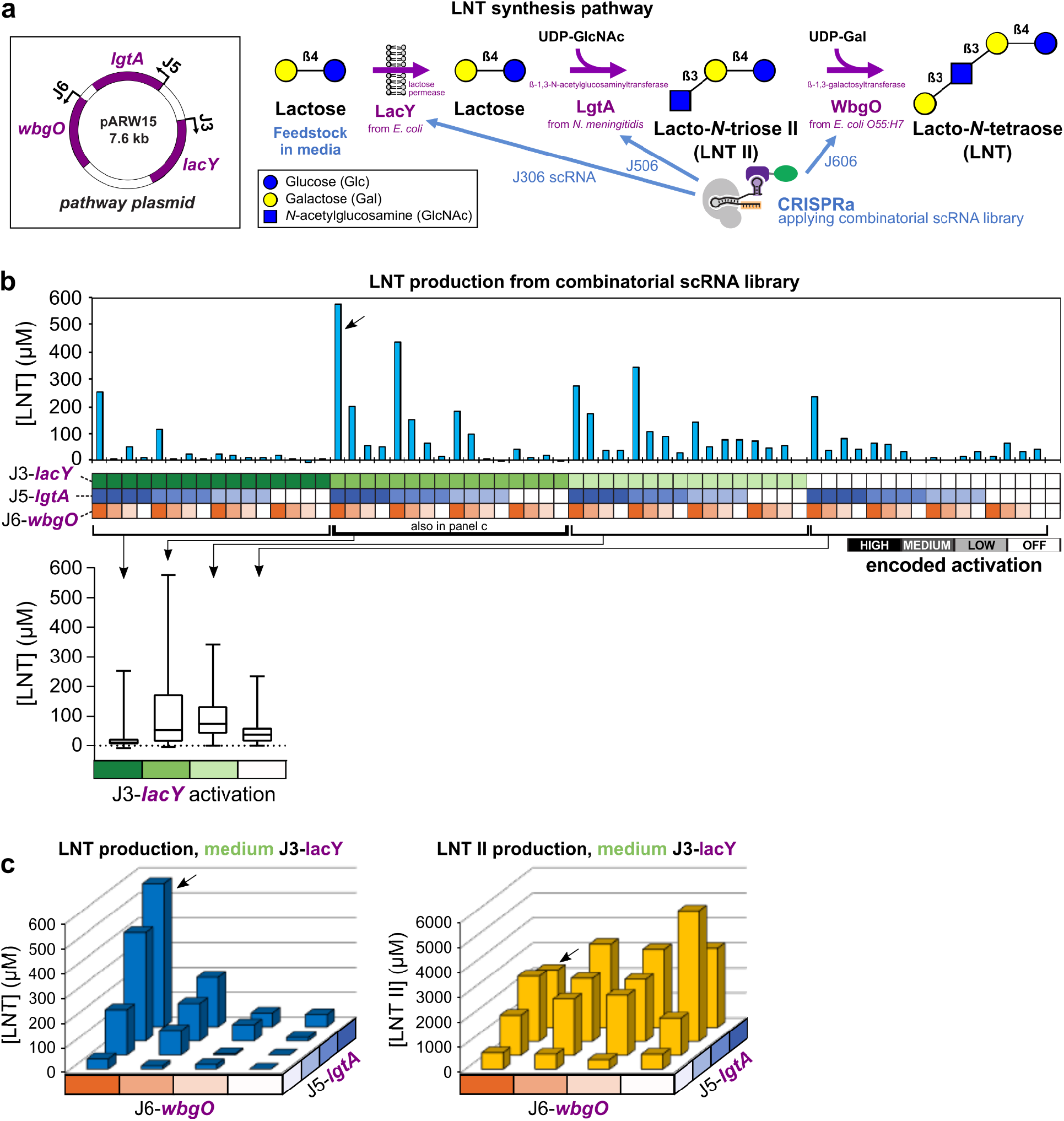
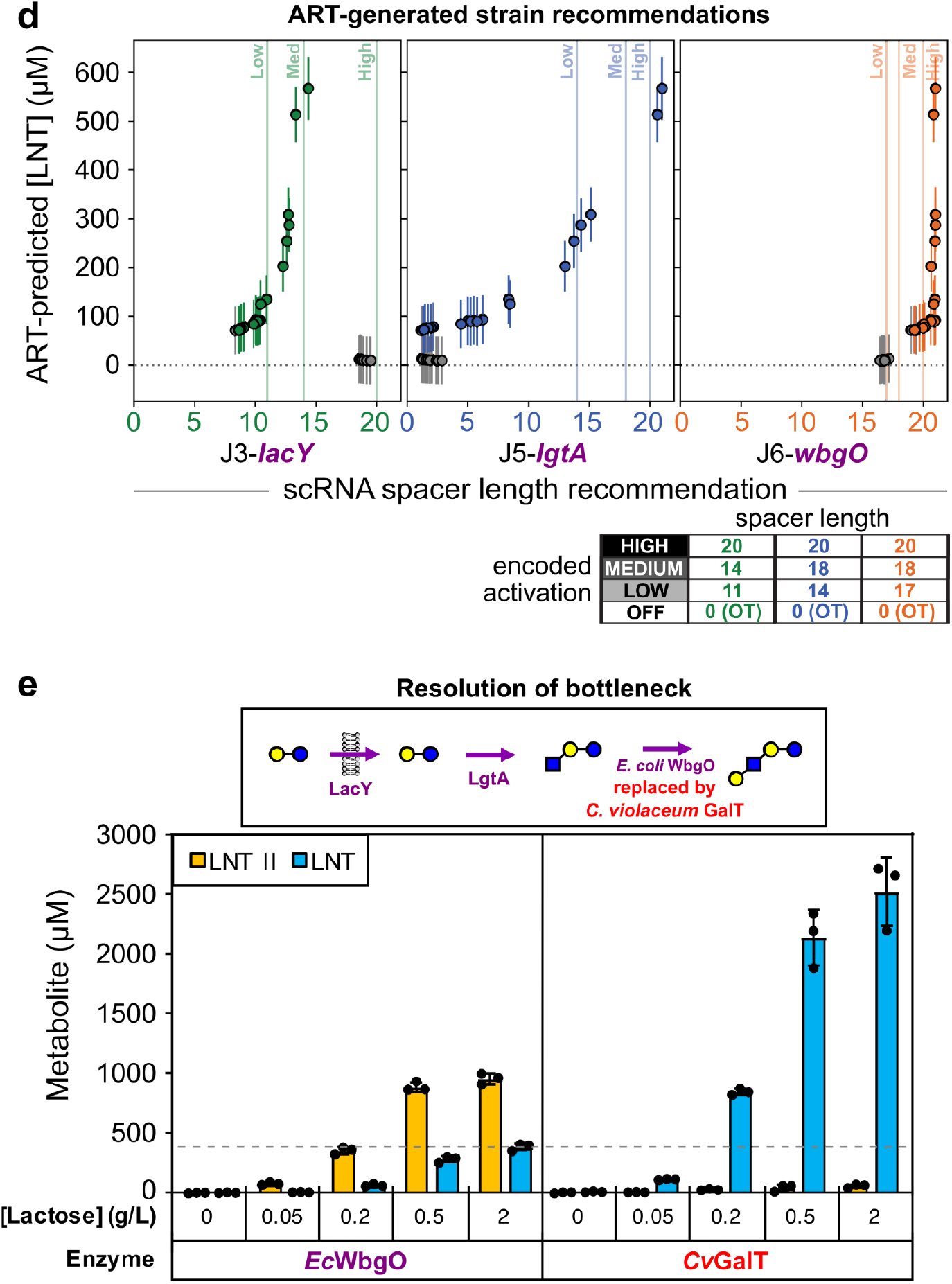
Combinatorial CRISPRa library applied to an HMO biosynthesis pathway identifies high-producing strains and pathway bottlenecks. **a** The LNT pathway consists of *lacY* overexpression from the J3 promoter, *lgtA* from the J5 promoter, and *wbgO* from the J6 promoter, activated by the J306, J506, and J606 scRNAs, respectively. The pathway enzymes import and elongate the lactose feedstock into a triose (LNT II) and then a tetraose (LNT). The pathway is expressed in an *E. coli* host with a *lacZ* knockout (JM109). The substrates UDP-GlcNAc and UDP-Gal are derived from endogenous metabolism. **b** HPLC analysis of supernatant from cultured library members indicates LNT production levels from the scRNA library. The highest producing strain (#17, black arrow) was used in the galactosyltransferase comparison in **d.** The *x*-axis heatmap is color coded to indicate the encoded promoter expression for each strain, as described on the bottom right. The 65th strain (right) is a no-pathway control culture carrying an empty vector. For comparison, LNT II levels are shown in Figure S6. **c** Dependence of LNT (left panel) and LNT II (right panel) production on *lgtA* and *wbgO* activation highlights sensitivity to wbgO activation and accumulation of LNT II. Only medium-*lacY* strains are shown here, due to their rich variance across the subset (box plot in **b**). The highest producing strain (#17) is again highlighted with a black arrow. **d** Computational strain recommendations from the Automated Recommendation Tool (ART) and their predicted LNT titers. Directed to try to maximize LNT production, ART generated 32 strain recommendations, defined by their scRNA truncation levels (spacer lengths measured in nucleotides, though non-integer values are allowed here), which determine degree of CRISPR activation (lower right). The top 20 strains, ordered by predicted LNT titer, are highlighted in color on each subgraph, while the bottom 12 are shown in grey. The same 32 strains are shown on each subgraph. Spacer lengths defined as high, medium, and low expression in the experimental scRNA library are indicated as vertical lines for each channel. Recommendation bounds were constrained between 0 and 21 nucleotides. ART-recommended strains tend to favor medium LacY expression, a wide range of LgtA levels, and only high WbgO expression. Error bars in **d** indicate the 95% credible interval of the predictive posterior distribution. See Supplementary Figure 13 for correlation of predicted versus observed LNT production. See Supplementary Figure 14 for how recommendations are combined within each strain. **e** A more active β-1,3-galactosyltransferase enzyme from *C. violaceum*^72^ resolves accumulation of LNT II. The lower-activity WbgO enzyme, even at high CRISPR activation, results in significant accumulation of LNT II, scaling with initial feedstock concentration (left panel). The additional activity of *Cv*GalT relieves the intermediate accumulation and results in higher LNT titers (right panel). The horizontal line indicates LNT titer achieved with WbgO and 2 g/L initial lactose, showing that *Cv*GalT can achieve similar titer using only 0.05–0.2 g/L initial lactose. Bar values in **e** represent the average ± standard deviation calculated from *n* = 3 biologically independent samples. Source data for **b**, **c**, **d**, and **e** are provided as a Source Data file. The sequence of the pathway plasmid containing J3-*lacY*, J5-*lgtA* and J6-*wbgO* or J6-*CvGalT* is described in the Supplementary Methods.

To establish CRISPRa control of LNT production, we generated a new output plasmid in which expression of the codon-optimized *lacY*, *lgtA* and *wbgO* genes are independently controlled by the J3, J5, and J6 synthetic promoters, respectively (Figure 6a). We delivered this LNT pathway plasmid, together with our existing multi-scRNA library, to the *lacZ* knockout *E. coli* strain JM109. Using HPLC to quantify accumulation in the culture supernatant of LNT and intermediate metabolite LNT II, we found a wide range of extracellular titers across the library, from zero to nearly 600 μM LNT (Figure 6b). A majority of the strains produced low or no LNT in supernatant, including some of the highest-expressing variants. For example, the strain with maximal expression (high-*lacY*, high-*lgtA*, high-*wbgO*) produced only 252 μM LNT (178 mg/L), while a strain with reduced *lacY* activation (medium-*lacY*, high-*lgtA*, high-*wbgO*) produced 576 μM LNT (408 mg/L). In general, we found that LNT production was compromised in the strains where *lacY* expression was highest, with only two out of 16 high-*lacY* strains producing >50 μM LNT (Figure 6b, left). This finding is consistent with toxic proton transport resulting from LacY activity^69,70^, and exemplifies an underlying mechanism of non-monotonic genotype-phenotype relationship. When *lacY* is reduced to medium levels, there is a large spread in LNT production, with eight out of 16 strains producing >50 μM LNT (Figure 6b). The J3-*lacY* local maximum highlights the importance of exploring a wide combinatorial space of enzyme expression, and the high variation of medium-*lacY* LNT production indicates the need for additional optimization of the other enzymes.

To understand the relative importance of LgtA and WbgO, we focused on the subset of medium *lacY* strains. In the medium-*lacY* sublibrary (Figure 6c), LNT production appeared to be more sensitive to variation of J6-*wbgO* expression than to variation of J5-*lgtA* expression. High LNT production (>400 uM) required high *wbgO* expression, indicating a steep expression-production relationship. For *lgtA*, high production was possible at high or medium expression, indicating a more gradual expression-production relationship. Reducing *wbgO* expression from high to low decreased titer from 576 μM to 56 μM (90.3% reduction compared to the maximum), but reducing *lgtA* expression from high to low only decreased titer to 182 μM (68.4% reduction) (Figure 6c). In most of these expression combinations, we also observed significant extracellular accumulation of the LNT II intermediate, the substrate for WbgO to convert into LNT. This accumulation was only avoided when *lgtA* was not activated (basal expression). When LNT II did accumulate, its titer did not depend strongly on low, medium, or high *lgtA* activation (Figure 6c). High LNT II titers were much more widespread across the library than high LNT titers (35 strains with LNT II titer above 25% maximal, compared to 10 strains for LNT) (Supplementary Figure 11). Taken together, these results suggest that limited β-1,3-galactosyltransferase activity of WbgO is a metabolic bottleneck in this pathway, confirming previous observations^52^. Our use of a combinatorial library to profile a multi-enzyme design space allowed us to easily characterize bottlenecks by probing for sensitive control points in the pathway.

A machine-learning analysis further validated the *wbgO* bottleneck. We used scRNA truncation levels from the library strains as inputs to the Automated Recommendation Tool (ART)^71^ to predict LNT production as a response variable, achieving high prediction accuracy (R^2^ = 0.7, Supplementary Figure 13) after training with the experimental LNT production data from the library. ART then used the predictions and uncertainties to make recommendations of the most productive enzyme expression levels. The most highly-recommended strains consistently prioritized maximal *wbgO* expression to achieve high LNT production. ART did not provide similarly stringent recommendations for *lacY* and *lgtA* (Figure 6d and Supplementary Figure 14), allowing substantial expression variation among LNT-productive strain recommendations. In agreement with the experimental library screen, these recommendations identify the *wbgO* bottleneck as a high priority for optimization, despite ART being unaware of LNT II accumulation. Furthermore, when allowed to recommend any spacer length up to 21 nucleotides, whether tested experimentally or not, ART frequently recommended *wbgO* levels above the highest experimentally-tested level. Collectively, these data underscore the idea that WbgO (β-1,3-galactosyltransferase) activity should be increased beyond maximal CRISPR activation of *wbgO* in this context.

To increase β-1,3-galactosyltransferase activity, we replaced WbgO with the GalT enzyme from *Chromobacterium violaceum* (*Cv*GalT), an enzyme with faster turnover.^72^ We placed *Cv*GalT under J6 control in the LNT pathway plasmid and paired it with the previously highest-producing scRNA library strain (medium-*lacY*, high-*lgtA*, high-*Cv*GalT). Compared to the corresponding WbgO strain, the *Cv*GalT strain produced a 5- to 10-fold increase in supernatant LNT titer, while LNT II accumulation decreased 5- to 20-fold, with the precise effect depending on the feedstock concentration (Figure 6e). These paired effects reflect the higher ability of *Cv*GalT to bind and convert LNT II before it is exported to accumulate in the supernatant^73^. The highest supernatant titer achieved by this system increased to 2.52 mM LNT (1.78 g/L), demonstrating a yield on lactose (0.432 mol/mol) much higher than previously reported yield for test-tube production (0.143 mol/mol)^73^. Relieving the bottleneck identified by our biosynthetic profiling approach therefore resulted in much more efficient conversion of feedstock into product.

Biosynthetic profiling of the LNT pathway by combinatorial CRISPRa indicated both the effects of *lacY* overexpression and the relative sensitivity of production to *wbgO* expression, demonstrating the potential of this approach to rapidly optimize enzyme expression levels. Crucially, the library is readily portable to different pathways. Applying combinatorial CRISPRa to a different pathway only requires a new output plasmid with the pathway enzymes expressed by the existing synthetic promoters, followed by cotransformation with the existing library of scRNA program plasmids.

## DISCUSSION

Synthetic biology and metabolic engineering offer a route for sustainable bioproduction of chemicals from renewable feedstocks. Many of these products are metabolically complex, requiring precise control over multi-gene networks to effectively redirect metabolic flux. Combinatorial CRISPRa programs can provide precise control over multiple targets, but require predictable scRNA efficacy. Developing general bacterial gRNA design rules and avoiding the typical trial-and-error validation of gRNA functionality will be an important factor for advancing multi-gene regulation programs. By combining computational RNA folding and experimental analyses, we uncovered strong correlations (*r_s_*=0.7-0.8) between CRISPR-activated expression and a set of thermodynamic and kinetic scRNA folding parameters^74,75^. Among the parameters examined, kinetic parameters associated with post-transcriptional RNA folding have the largest impacts on CRISPRa.

We found that a single kinetic parameter, Folding Barrier, can accurately predict bacterial CRISPRa across a broad range of expression levels, with a failure rate near zero for forward design of scRNAs. We speculate that the predictive value of Folding Barrier may be higher than that of Folding Energy because binding to dCas9 may stabilize the active scRNA structure (Supplementary Figures 2 and 3). The kinetic barrier to access the active structure determines the likelihood of dCas9 trapping the RNA in that structure, and is potentially more important than the intrinsic thermodynamic stability of the free RNA structure. dCas9 binding should also provide some resistance to RNA degradation^76^. The high predictability of scRNA design supplied by Folding Barrier should significantly facilitate the forward engineering of complex bacterial CRISPRa/i systems. Multi-guide applications that have remained inefficient or impractical with current gRNA failure rates, such as combinatorial expression screening^77^ or model- and data-driven strain engineering and optimization^18^, can therefore be accelerated. Recent metabolic engineering successes in related systems emphasize the value of predictive gRNA design^22,78^.

The Folding Barrier metric outperformed current state-of-the-art gRNA design tools in its ability to predict CRISPRa activity^21,30^. There are many possible explanations for the inability of existing models to apply to bacterial CRISPRa systems. First, many of these models account for genome structure, which will vary greatly between eukaryotes and prokaryotes^79,80^. Second, in regression models trained on large gene editing datasets, it is difficult to decouple gRNA efficiency from feedback on gene expression as part of the overall gene regulatory network, and therefore the predictions of these models may not be readily transferable between organisms. Third, these models were trained on unmodified gRNAs and do not capture potential folding effects of extended RNA elements included in scRNAs for bacterial CRISPRa. These models could likely be improved by incorporating biophysical parameters in their predictions. Finally, considerations of nucleic acid interactions in models tend to focus on the thermodynamics of spacer-DNA interactions, and neglect other important aspects of gRNA folding^29^. Developing models that combine solely sequence-based kinetic folding parameters with heuristics from large-scale functional screening should further improve our ability to design modified guide RNAs for bacterial CRISPRa.

Optimal multi-gene pathway expression could be influenced by many factors, possibly including total burden, enzyme imbalance, or toxic enzyme or metabolite effects. The difficulty in predicting these systems-level interactions means that finding global production optima often requires exploring large design spaces^81^. Toward this end, we successfully developed a scRNA library that can implement all combinations of four truncation-defined expression levels across three chosen genes, totaling 64 possible expression programs. For each of the pathways we examined, we found the optimal production to occur at non-maximal expression levels in at least one channel of expression (*rfp*, *sr*, and *lacY* in Figures 4, 5, and 6, respectively). Production from these pathways therefore maps ruggedly to the underlying design space of enzyme expression, and systematically profiling these effects revealed high-producing strains and also pathway bottlenecks potentially sensitive to optimization. Pursuing bottleneck optimization in the LNT pathway with an improved enzyme variant pushed titers into g/L magnitude. Broadly speaking, biosynthetic profiling using *trans*-acting scRNAs can greatly reduce the time needed to tune multi-gene programs, compared to traditional *cis*-acting tools like promoter, RBS, or ribozyme libraries^82,83^. We expect that the combinatorial scRNA library described here will provide a straightforward approach to identifying production maxima and optimizing burdensome pathways or toxic intermediate accumulation. In the future, this approach could be extended to non-model hosts with metabolic and physiological capabilities suitable for next-generation bioproduction applications^84–86^.

Combinatorial CRISPRa programs could also be extended to increase expression variation resolution or use alternative tuning methods^19,22,87^. Along with principles of high-dynamic-range CRISPRa promoter design^43^, the scRNA design rules from this work can in principle generate a virtually unlimited supply of high dynamic range CRISPR-activatable promoters. Expanding beyond the three synthetic promoters used here would allow activation of larger pathways, endogenously-targeted CRISPRa/i^16,88^ for flux optimization, or dynamic gene regulation through biosensors^89,90^. There are practical limits on the sizes of functional scRNA/gRNA arrays, due mostly to binding competition for a shared dCas9 pool^41,42^. New principles of gRNA design, including those reported in this work, and some autoregulatory circuit designs^91^ could be used to increase this limit and build large multi-guide programs. For large combinatorial libraries of genetic circuits, higher-throughput screening methods like biosensing technologies would be needed to screen through the added diversity^18,92,93^. For design spaces too large for current screening methods, data-driven and model-guided approaches like ART can be used to explore the full design space, informed by experimental efforts focused only on the most likely subsets of design parameters (Supplementary Figure 15). An optimal subset size depends on the complexity of the pathway to be optimized, but the experimental CRISPRa profiling approach can ease the construction of these subsets.

Iterative cycles of model-guided optimization and data-driven model refinement present a promising path forward for rapid generation and optimization of biosynthetic pathways. The value of this approach is especially demonstrated when used together with combinatorial CRISPRa/i programs to access model predictions and build iteratively improved strains. Optimized metabolic engineering programs can help realize a circular bioeconomy that decreases our reliance on fossil feedstocks for production of industrial chemicals and materials. To help meet this challenge, synthetic biologists can use the tools presented in this work to rapidly optimize strains for bioproduction of valuable chemicals from renewable feedstocks.

## METHODS

### Bacterial Strains and Plasmid Construction

Bacterial strains used in this study are described in Supplementary Table 1. JM109 was a gift from Joachim Messing (Addgene plasmid #49761)^94^. Plasmids were cloned using standard molecular biology protocols and are described in Supplementary Table 2. Guide RNA target sequences are provided in Supplementary Table 3. Orthogonal target sequences replacing J306 were 20 bp sequences selected at random from the human genome. Plasmids expressing the CRISPRa components (dCas9, the activation domain MCP-SoxS, and one or more scRNAs) were constructed using a p15A vector. *S. pyogenes* dCas9 (*Sp*-dCas9) was expressed using the endogenous *Sp.pCas9* promoter. The MCP-SoxS activation domain containing mutant SoxS (R93A and/or S101A; see Supplementary Table 2)^12^ was expressed using the BBa_J23107 promoter (http://parts.igem.org). The scRNAs were expressed using either the BBa_J23119 promoter or the BBa_J23105 (Supplementary Figure 9), unless otherwise noted. scRNAs used the b2 design, in which the endogenous tracr terminator hairpin upstream of MS2 is removed^11^. Plasmids expressing target genes for CRISPRa were constructed using a low-copy pSC101** vector. mRFP1, sfGFP, mTagBFP2, or metabolic pathway genes were expressed from the weak BBa_J23117 minimal promoter preceded by synthetic DNA sequences containing the CRISPRa target sites. Pathway gene RBSs were selected from a previously reported list^95^ and predicted to have high strength^96^ in the new context.

### Computational analysis of scRNA activity

Energetic parameters were generated using the RNAfold, RNAeval, RNAduplex, and Findpath programs from the ViennaRNA Package version 2.3.5^27^. Sequences of full scRNAs were input to a custom script that returned the following parameters. Folding Barrier was calculated by using Findpath to predict the barrier height for the direct refolding pathway from the MFE structure to the active structure (see Supplementary Figure 2). The active structure is defined as the structure in which the Cas9-binding handle is correctly folded and the spacer is unstructured. Binding Energy was calculated by evaluating the RNA-RNA free energy of the spacer sequence binding to its reverse-complement sequence using RNAduplex. The Folding Energy, or free energy difference between the MFE structure and the active structure, was evaluated using RNAfold with constraint folding. Folding Energy was then added to the Binding Energy in order to estimate the net energetics of binding to a single-stranded target sequence. This sum yields the Net Binding Energy, or the free energy difference between the MFE and the bound state. All scRNA sequences were verified to have a prediction of correct folding of the MS2 aptamer at the 3’ end, to avoid confounding cases of target occupancy without bound MCP-SoxS.

For the purpose of comparison to this work’s scRNA efficacy predictions, the Doench’16, Azimuth *in vitro*, and Moreno-Mateos tools for CRISPR guide design and evaluation were implemented using the CRISPOR webserver (http://crispor.tefor.net/)^97^. The 20 bp variable target sites for scRNA-directed CRISPRa flanked by 50 bp of upstream and 50 bp of downstream sequence (120 bp total) were used as inputs (sequence is shown in the Supplementary Methods). Analysis was carried out with the default settings for “No Genome” and Protospacer Adjacent Motif (PAM) set to “20bp-NGG - SpCas9, SpCas9-HF1, eSpCas9 1.1”. Each 20 bp target was evaluated using the “predicted guide efficiency” outputs generated by the respective CRISPR guide design tools.

### Construction of combinatorial scRNA library

To encode high, medium, and low activation of the J3, J5, and J6 promoters, we selected the 20, 14, and 11 nucleotide variants of J306; the 20, 18, and 14 nucleotide variants of J506; and the 20, 18, and 17 nucleotide variants of J606, respectively. For all three promoters, a fourth, unactivated condition was included *via* an off-target scRNA with a spacer sequence not complementary to any of the synthetic promoters. In the CRISPRa component plasmid library, a three-member array of scRNA expression, each with its own BBa_J23105 promoter and terminator, was constructed for every possible combination of the J306, J506, and J606 truncation variants. Including the off-target versions, this resulted in a 64-member combinatorial library of CRISPRa component plasmids, accounting for all combinations of high, medium, low, and baseline expression of all three synthetic promoters.

### Plate Reader Experiments

Single colonies from LB-agar plates were inoculated in triplicate in 500 μL EZ-RDM (Teknova) supplemented with appropriate antibiotics and grown in 96-deep-well plates at 37°C and shaking on a microplate orbital shaker (Heidolph) overnight. For mRFP1 detection, 150 μL of the overnight culture were transferred into a flat, clear-bottomed black 96-well plate and the OD_600_ and fluorescence (excitation wavelength: 540 nm; emission wavelength: 600 nm) were measured in a Biotek Synergy HTX plate reader.

### Flow Cytometry

Single colonies from LB-agar plates were inoculated in triplicate in 500 μL EZ-RDM (Teknova) supplemented with appropriate antibiotics and grown in 96-deep-well plates at 37°C and shaking on a microplate orbital shaker (Heidolph). Overnight cultures were diluted in 1:100 in DPBS and analyzed on a MACSQuant VYB flow cytometer (Miltenyi Biotec) using a previously described strategy to gate for single cells^11^. A side scatter threshold trigger (SSC-H) was applied to enrich for single cells. A narrow gate along the diagonal line on the SSC-H vs SSC-A plot was selected to exclude the events where multiple cells were grouped together. Within the selected population, events that appeared on the edges of the FSC-A vs. SSC-A plot and the fluorescence histogram were excluded. For sfGFP detection, the excitation wavelength was 488 nm and emission wavelength was 525 nm (50 nm bandpass). For mTagBFP2 detection, the excitation wavelength was 405 nm and emission wavelength was 450 nm (50 nm bandpass). For mRFP1 detection, the excitation wavelength was 561 nm and emission wavelength was 615 nm (20 nm bandpass). Data were analyzed using FlowJo. Median values were normalized to the highest observed value within each channel and were baseline-subtracted using a strain lacking the genes encoding the fluorescent proteins.

### Biopterin production experiments

Single colonies from LB-agar plates were inoculated in triplicate in 500 μL EZ-RDM (Teknova) supplemented with appropriate antibiotics and grown overnight in 96-deep-well plates at 37°C with shaking. 100 μL of the overnight culture were transferred into a flat, clear-bottomed black 96-well plate and the OD_600_ and fluorescence (excitation wavelength: 340 nm; emission wavelength: 440 nm) were measured in a monochromator-equipped plate reader (Tecan Infinite M1000) to assess pteridine production^15,98–100^. Fluorescence corresponds to BH4 and its oxidized derivative BH2.

### Lacto-*N*-tetraose production experiments

Single colonies from LB-agar plates were inoculated in singlicate in 2 mL EZ-RDM (Teknova) with 10 g/L glucose, 2 g/L lactose and supplemented with appropriate antibiotics. For the JM109 strain, agar plates used 100 μg/mL chloramphenicol and 100 μg/mL carbenicillin to avoid slightly chloramphenicol-resistant background growth, but liquid cultures used the more typical concentrations of 25 μg/mL chloramphenicol and 100 μg/mL carbenicillin. Cultures were grown in 14 mL polypropylene culture tubes at 37°C with shaking for 48 h. 500 μL of supernatant from each culture were loaded onto 10 kDa microcentrifuge filters (Millipore) and spun for 20 min at 14000 rcf. 1 μL of filtered supernatants were assayed with a Shimadzu HPLC using UV-vis detection at 210 nm. Lacto-*N*-tetraose (LNT) was separated using a Rezex ROA-Organic Acid H+ column (Phenomenex) and a 20 mM H_2_SO_4_ isocratic mobile phase. A standard curve was prepared by spiking known amounts of LNT or LNT II into supernatants derived from cultures of JM109 *E. coli* transformed with empty vectors. Product LNT was observed at 10.6 minutes, and intermediate LNT II, a triose, was observed at 11.4 minutes. LNT and LNT II peak areas were normalized by the area of an endogenous peak observed at 9.1 minutes. Normalized peak areas were baseline-subtracted using a control strain lacking the pathway genes. Cell pellets also contained significant LNT, as previously reported^52^ and verified in pellets lysed by boiling, but the difficulty of consistently quantifying lysis efficiency and the rich variation in supernatant titers led us to consider mainly supernatant data for comparative analysis.

### ART predictions and recommendations

The Automated Recommendation Tool (ART)^71^ was trained on the 64 experimental LNT strains, with J3-*lacY*, J5-*lgtA*, and J6-*wbgO* CRISPRa variations as input variables and LNT production as the response variable. ART is an ensemble model that linearly combines a variety of machine learning models. Models are cross-validated individually on the data, and the weight for each model represents its performance (higher for better-performing models, lower for worse-performing ones). These weights are considered as random variables with probability distributions obtained through Monte Carlo sampling. This approach enables quantification of both the prediction mean and uncertainty for any given input data. Predictions are possible at any point in the possible design space, not limited to the discrete high, medium, low, and off-target activation levels comprising the experimental library. ART was trained, however, using the exact activation levels from the experimental library, expressed as spacer length in nucleotides (e.g. 20 for high, 14 for medium, and 11 for low in the J3 case). In all cases, off-target spacers were expressed as an input of 0. Cross-validation correlations were also computed using exact library activation levels.

For the strain recommendations, strains are defined by their recommended input levels, expressed in scRNA spacer length for that channel. ART was allowed to recommend any spacer length from 0 to 21 nucleotides (non-integers allowed), with the constraint that new designs had to be at least 1 nucleotide away (in at least one dimension) from other recommendations and from training data. The 32 recommended strains resulting in the highest predicted LNT concentration were obtained from ART. In this work, recommendations were fully exploitative (α = 0), meaning that they prioritized maximizing LNT as opposed to minimizing the uncertainty in LNT predictions.

### Statistics

Statistical significance was calculated using two-tailed unpaired Welch’s *t*-tests. Quantitative correlations are expressed as Pearson correlations. Rank-order correlations are expressed as Spearman correlations. Hill function (Figure 2d) was fitted as the following nonlinear function in GraphPad Prism, using least squares regression:

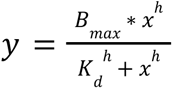

Dose-response function (Supplementary Figure 8) was fitted as the following nonlinear function in GraphPad Prism, using least squares regression:

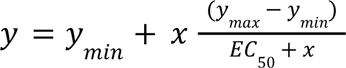

Simple linear and exponential fits (Supplementary Figures 1, 11, and 14) were performed using default settings in GraphPad Prism or Microsoft Excel.

## AUTHOR CONTRIBUTIONS

J.F., D.S.Y., I.F., H.G.M., J.G.Z, and J.M.C. designed experiments. J.F., D.S.Y., I.F., R.C., C.K., A.W., T.G.P., and P.C.K. performed experiments and analyzed data. J.F., D.S.Y., I.F., R.C., C.K., P.C.K., J.G.Z, and J.M.C. wrote and edited the manuscript with input from all of the authors.

## ACKNOWLEDGEMENTS

We thank Semira Beraki for technical assistance and Venkata P. Chavali for technical assistance and helpful discussions. We thank members of the Zalatan and Carothers groups for advice. This work was supported by U.S. National Science Foundation Awards MCB 1817623 (awarded to J.G.Z. and J.M.C.), MCB 2032794 (J.M.C. and J.G.Z.), CBET 1844152 (J.M.C.), U.S. Department of Energy Awards DE-EE0008927 and DE-SC0023091 (J.M.C., J.G.Z. and H.G.M.) and a grant from BASF, Inc. (J.M.C.).

