## Supplementary Information for "Guide RNA structure design enables combinatorial CRISPRa programs for biosynthetic profiling"

### SUPPLEMENTARY FIGURES

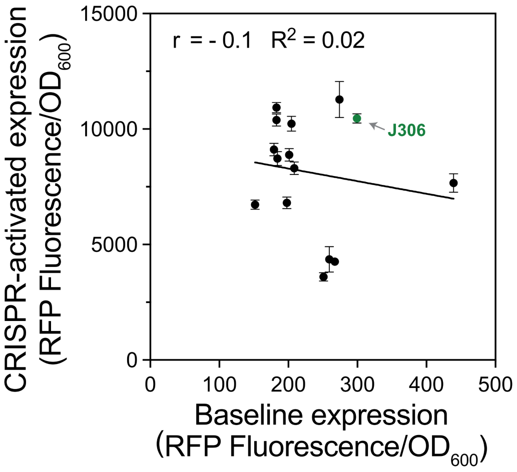


**Supplementary Figure 1. Small differences in baseline expression in synthetic promoters with different scRNA target sites do not explain CRISPR-activated expression level.** The CRISPR-activated gene expression of the synthetic promoters from Figure 2B is plotted against their baseline, unactivated expression. The J3 promoter was previously described[1]; each of the 14 other promoters contains a randomly selected scRNA target site and is targeted by a scRNA with complementary target sequence. Values on the *x*-axis represent the baseline expression of each promoter, obtained by measuring the Fluorescence/$OD_{600}$ of strains harboring each synthetic promoter and expressing an off-target scRNA. Values on the *y*-axis depict the Fluorescence/$OD_{600}$ of strains harboring each synthetic promoter and expressing the matching scRNAs, as in Figure 2B. The green dot indicated by the arrow represents a strain harboring the previously described J3 promoter and expressing its cognate J306 scRNA. r and $R^2$ represent, respectively, the Pearson correlation coefficient and the coefficient of determination for the linear fit between baseline expression and CRISPR-activated expression. Values represent the average ± standard deviation calculated from $n$ = 3 biologically independent samples. Source data are provided as a Source Data file.

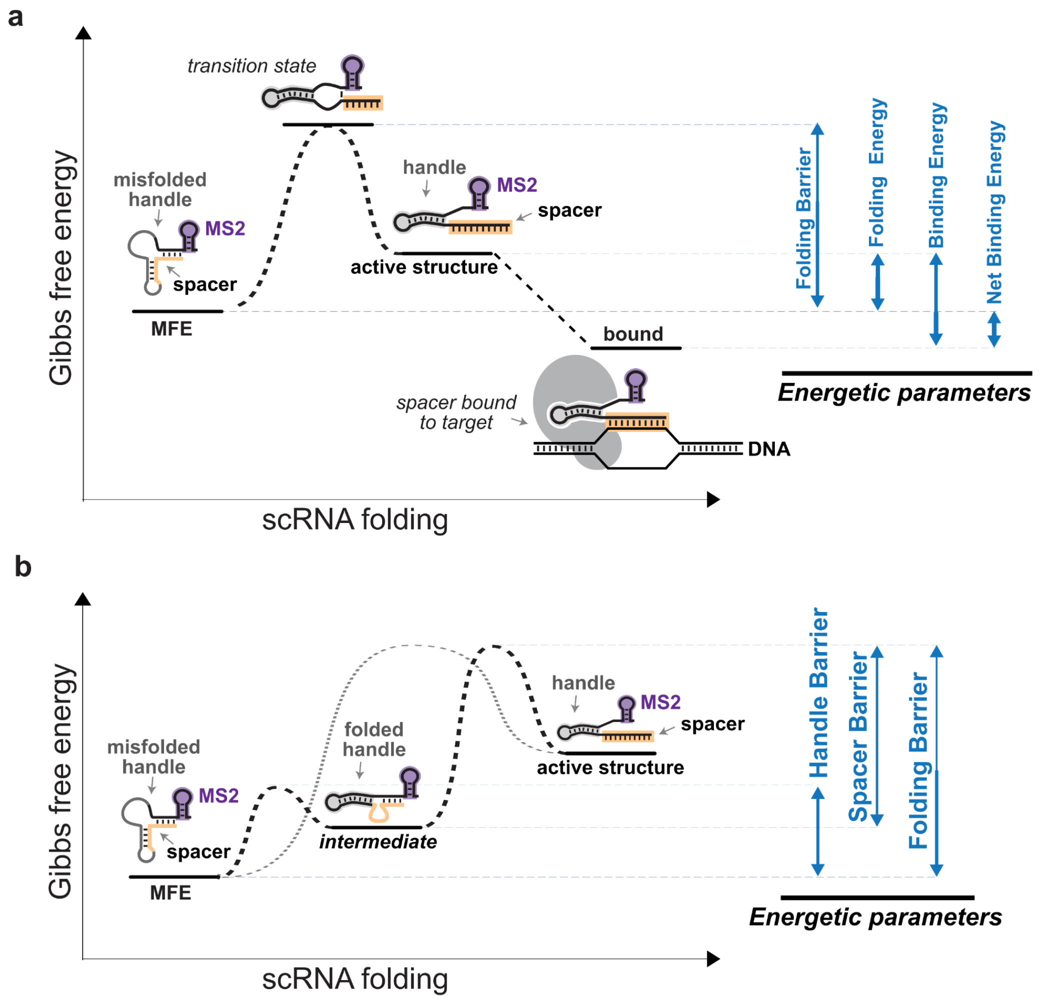


**Supplementary Figure 2. Kinetic and thermodynamic parameters provide a quantitative description of scRNA folding. a** Kinetic and thermodynamic parameters computationally analyze scRNA folding. The *y*-axis represents the Gibbs free energy of representatives states in scRNA folding. The *x*-axis represents the scRNA folding coordinate. The MFE state indicates the Minimum Free Energy structure adopted by an scRNA, which in this example contains a misfolded dCas9-binding handle and a sequestered spacer. The active structure indicates the correctly folded scRNA in which the spacer is unstructured and the dCas9-binding handle and MS2 hairpin are correctly folded. Folding Barrier represents the free energy barrier that must be overcome to fold the scRNA from the MFE state to the active structure, and is calculated as the free energy difference between the MFE state and the transition state to the active structure. The Folding Energy represents the free energy difference between the MFE state and the active structure. The bound state illustrates the binding of the scRNA to the target DNA and is approximated as the free energy of the scRNA in the active structure bound to a RNA 20-mer complementary to the spacer sequence. The Binding Energy represents the free energy difference between the active state and the bound state. The Net Binding Energy represents the free energy difference between the MFE state and the bound state. **b** Additional metrics were investigated to understand the role of kinetics in scRNA folding. An intermediate state between the MFE state and the active structure was defined in which the dCas9-binding handle is correctly folded. Handle Barrier represents the free energy barrier between the MFE state and the intermediate state with the dCas9-binding handle folded. Spacer Barrier represents the free energy barrier between the intermediate state and the active structure.

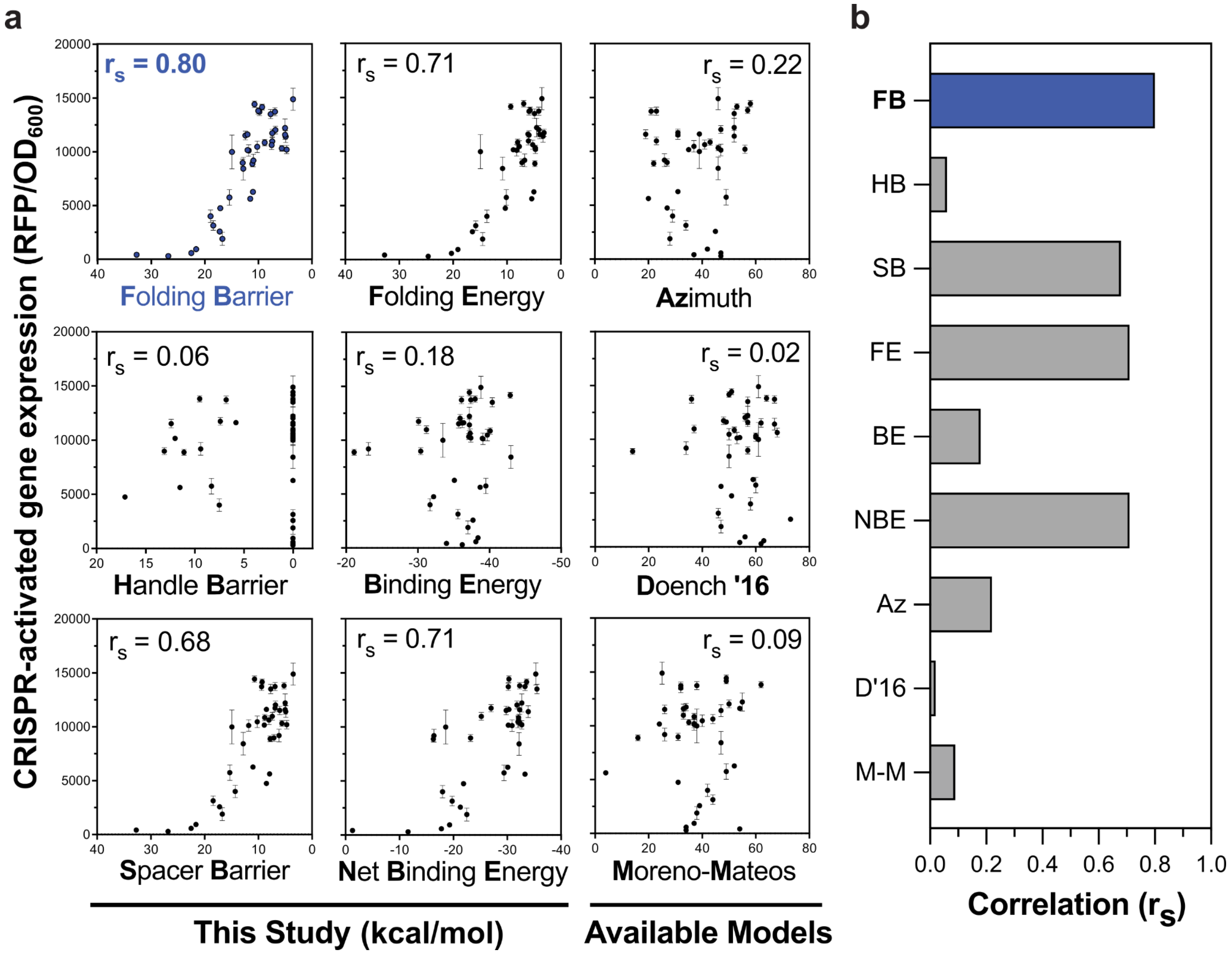


**Supplementary Figure 3. Folding barrier outperforms popular gRNA selection models for predicting scRNA activity in CRISPRa experiments.** **a** Comparison of kinetic and thermodynamic parameters from this study with three popular gRNA selection models for scRNA design. Individual plots describe the performance of the indicated parameter or model. A schematic of each parameter from this study is provided in Supplementary Figure 2. The gRNA selection models tested are Azimuth, Doench '16[2] and Moreno-Mateos[3], which were the three best performing models from the set of models found in the CRISPOR webserver[4]. Each plot contains 39 data points, corresponding to the scRNAs targeting each of the synthetic promoters described in Figure 2d. In each plot, the *y*-axis shows the CRISPR-activated RFP expression of strains harboring each synthetic promoter and expressing the matching scRNA. The *x*-axes represent different computed parameters for scRNA function. For the parameters described in this study, the *x*-axis is kcal/mol on a reverse scale (high values on the left, low values on the right). The reverse scale places scRNAs predicted to be more active to the right side of each plot. The depicted parameters from this study are Folding Barrier, Handle Barrier, Spacer Barrier, Folding Energy, Binding Energy, and Net Binding Energy. Among those, Folding Barrier, Folding Energy, and Net Binding Energy were most correlated with expression. For the Azimuth, Doench ‘16, and Moreno-Mateos models, values on the *x*-axis represent scores assigned by each model when the scRNA target site for each of the synthetic promoters described in Figure 2D is used as an input. Again, more active predictions should be to the right side of the plot. $r_s$ represents the Spearman’s rank order correlation coefficient between the predicted scRNA activity based on the *x*-axis values and experimental CRISPR-activated expression. **b** Correlation coefficients between predicted scRNA activity and experimental results. Bars represent the Spearman’s rank order correlation coefficient between predicted scRNA activity and the corresponding experimental CRISPR-activated expression for each parameter depicted in panel **a**. Labels are abbreviations of the metrics from this study and the gRNA selection models. FB: Folding Barrier, HB: Handle Barrier, SB: Spacer Barrier, FE: Folding Energy, BE: Binding Energy, NBE: Net Binding Energy, Az: Azimuth. D’16: Doench ‘16, M-M: Moreno-Mateos. Values in panel a represent the average ± standard deviation calculated from *n* = 3 biologically independent samples. Source data are provided as a Source Data file.

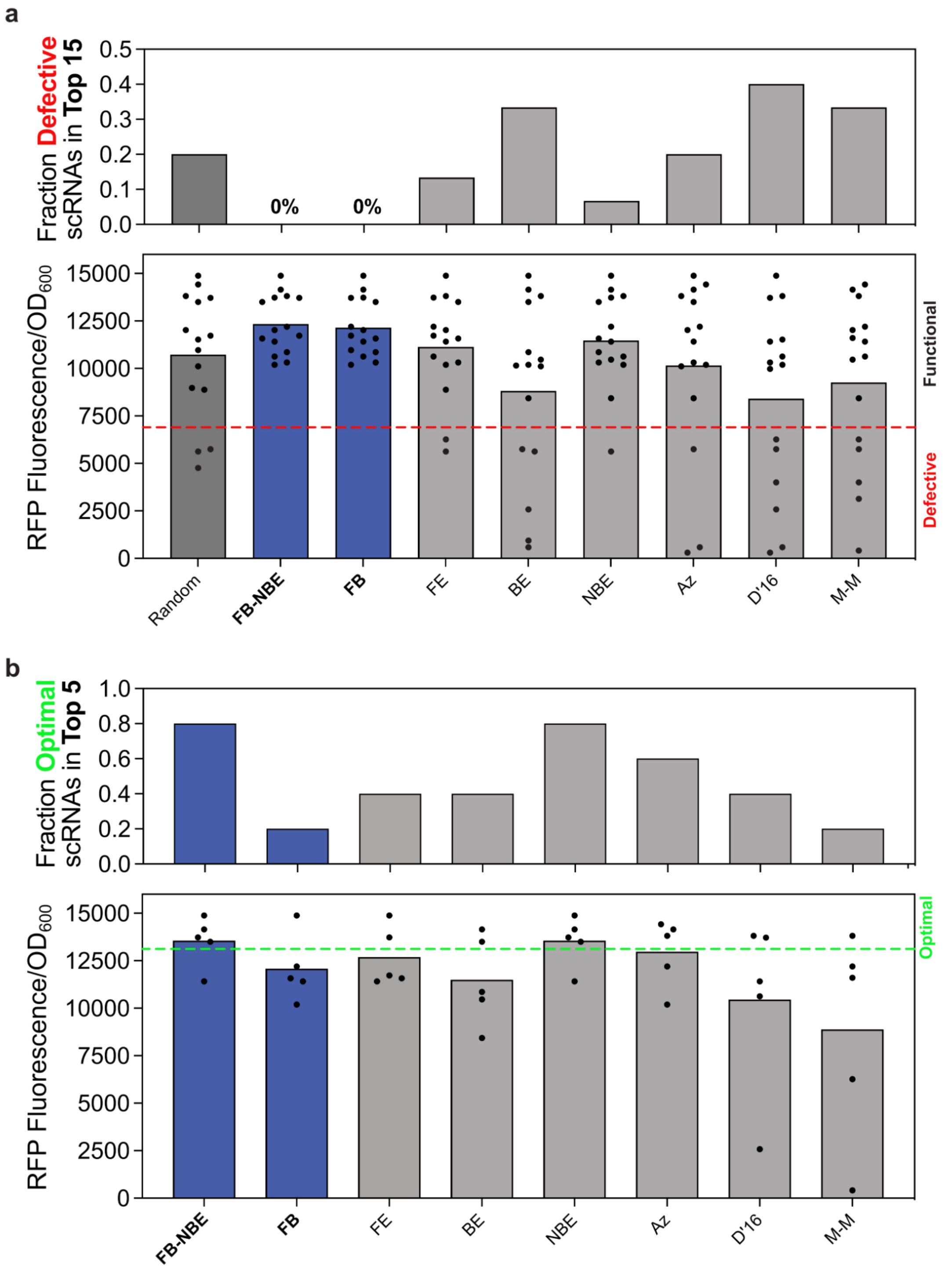


**Supplementary Figure 4. Selecting scRNAs based on Folding Barrier and Net Binding Energy parameters avoids defective scRNAs and enriches for optimal scRNAs. a** Selecting scRNAs with a low Folding Barrier avoids defective scRNAs. A schematic of each parameter from this study is provided in Supplementary Figure 2. Of the 39 scRNAs described in Figure 2D, the 15 scRNAs predicted by each parameter to perform the best are shown here. For the parameters described in this study, better performance is predicted by lower Gibbs free energy (Net Binding Energy, Folding Energy, Binding Energy) or smaller free energy barriers (Folding Barrier). FE-NBE was obtained by first screening the scRNAs for Folding Barriers ≤10 kcal/mol, and then ranking the remaining scRNAs based on their Net Binding Energy (Supplementary Methods). For the three available gRNA selection models, the highest-scoring 15 scRNAs are shown here[2,3] (using the scRNA target site as an input). The top graph shows the fraction of defective scRNAs found in the top 15 scRNAs selected by each parameter. Defective scRNAs are defined as those used in strains that yielded ≤50% of J306-level CRISPR-activated expression (Fluorescence/$OD_{600}$). The bottom graph shows the CRISPR-activated expression of strains expressing the top 15 scRNAs selected by each parameter. Dots represent individual strains, while bars represent the average expression from each set. The dashed red line shows the threshold for defective scRNAs. **b** Screening scRNAs with low Folding Barrier and minimizing Net Binding Energy yields optimal scRNAs. The top graph shows the fraction of optimal scRNAs found in the top 5 scRNAs selected by each parameter. scRNAs were selected in the same way described in panel **a**. Optimal scRNAs are defined as those used in strains that yielded ≥95% of J306-level CRISPR-activated expression (Fluorescence/$OD_{600}$). The bottom graph shows the CRISPR-activated expression of strains expressing the top 5 scRNAs selected by each parameter. FB-NBE and NBE happened to select the same scRNAs in this case, because they were all under the FB threshold. FB still has value for avoiding defective scRNAs in other cases, though, as seen in panel **a**. Inclusion of NBE ranking in FB-screened subsets does yield slightly more active scRNAs in this case (see Supplementary Methods for additional discussion). Dots represent individual strains, while bars represent the average expression from each set. The dashed green line shows the threshold for optimal scRNAs. FB: scRNA Folding Barrier, NBE: Net Binding Energy, HB: Handle Barrier, SB: Spacer Barrier, FE: Folding Energy, BE: Binding Energy, Az: Azimuth. D'16: Doench '16, M-M: Moreno-Mateos. Source data are provided as a Source Data file.

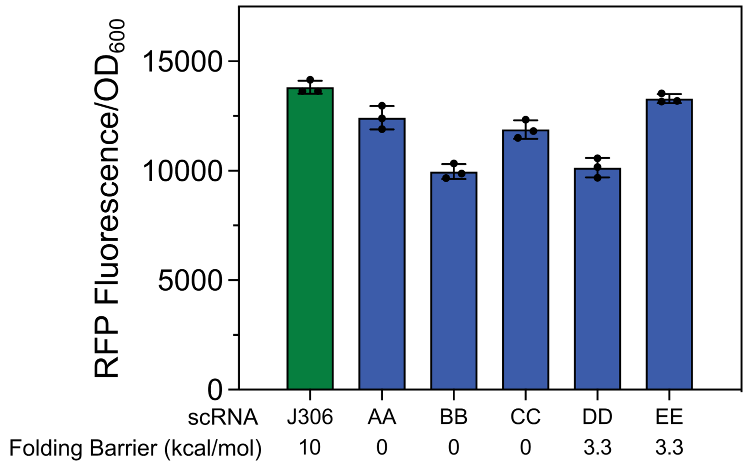


**Supplementary Figure 5. Forward engineering of scRNAs with minimal Folding Barriers.** As a test of Folding Barrier's predictive power for forward design of highly functional scRNAs, five additional scRNAs were specifically designed to have Folding Barriers below 5 kcal/mol, and as close to 0 kcal/mol as possible. To preserve the structure of the dCas9-binding handle, we had to provide additional stability to the handle by altering the sequence relative to wild-type. For AA, BB, and CC, the MFE was predicted to be the active structure, hence the zero barrier. CRISPRa using these scRNAs was very active for all five, as expected from Folding Barriers lower than the 10 kcal/mol threshold, but interestingly they were not the highest-performing scRNAs in the overall set included in this work. This relates to the sigmoidal fit proposed in Figure 2d, where Folding Barrier loses some of its predictive power below the threshold of FB ≤ 10 kcal/mol. Values represent the average calculated from $n$ = 3 biologically independent samples. Source data are provided as a Source Data file.

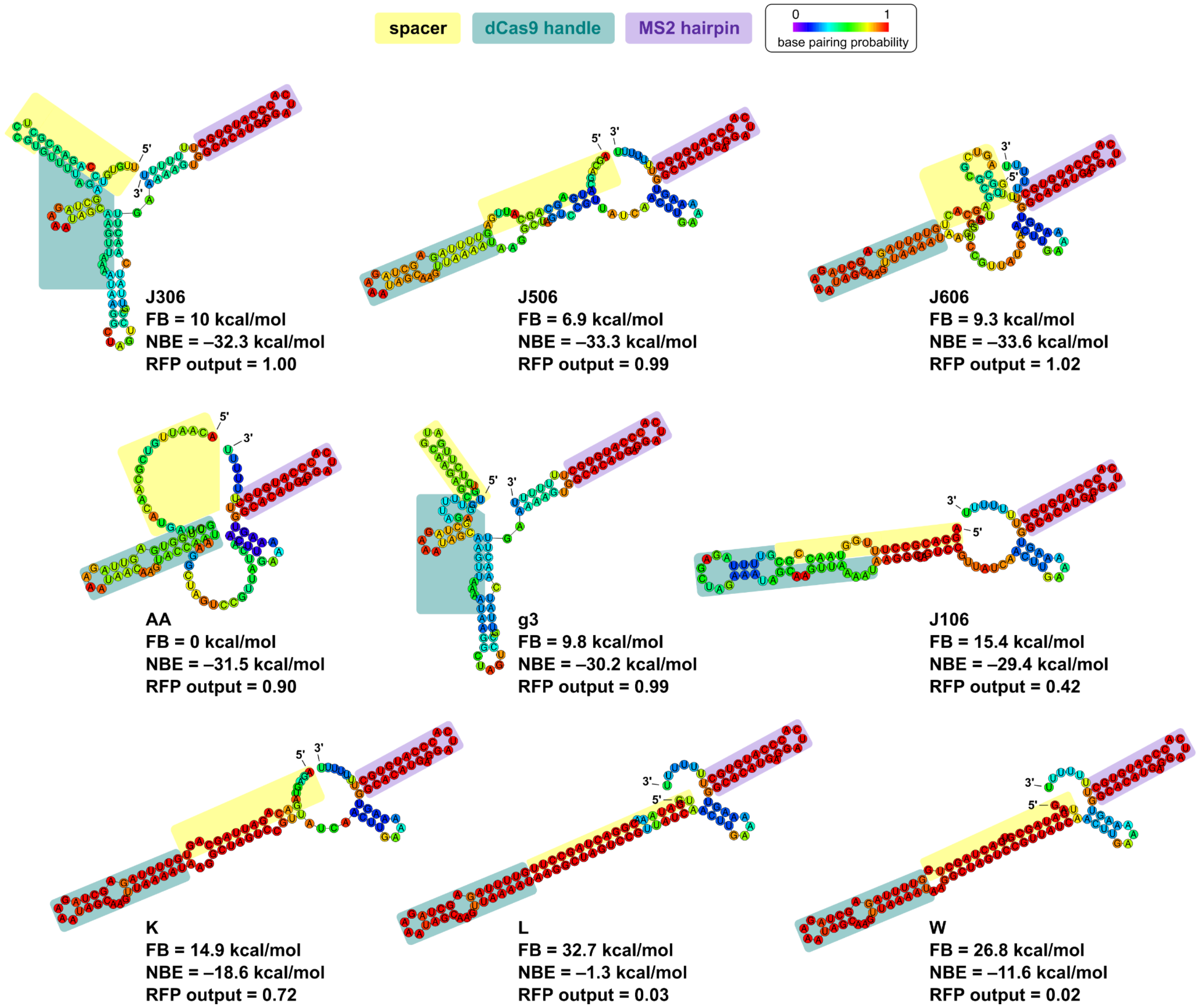


**Supplementary Figure 6. Select scRNA MFE structures.** Minimum Free Energy (MFE) structure predictions were generated using the RNAfold WebServer[5], and the resulting dot-bracket notations are also included in Supplementary Table 6. The scRNAs used in this study's combinatorial expression library—J306, J506, and J606—are shown in the top row, while the rest are roughly in ascending order of Folding Barrier and descending order of RFP output. Colors indicating base pairing probability near 0 indicate structures that probably require little energy to break and reform into an MFE-adjacent structure, as suggested by low Folding Barriers. High FBs, accompanied by low output, are often the result of strong hybridization of the spacer sequence to an internal sequence, exemplified by K, L, and W.

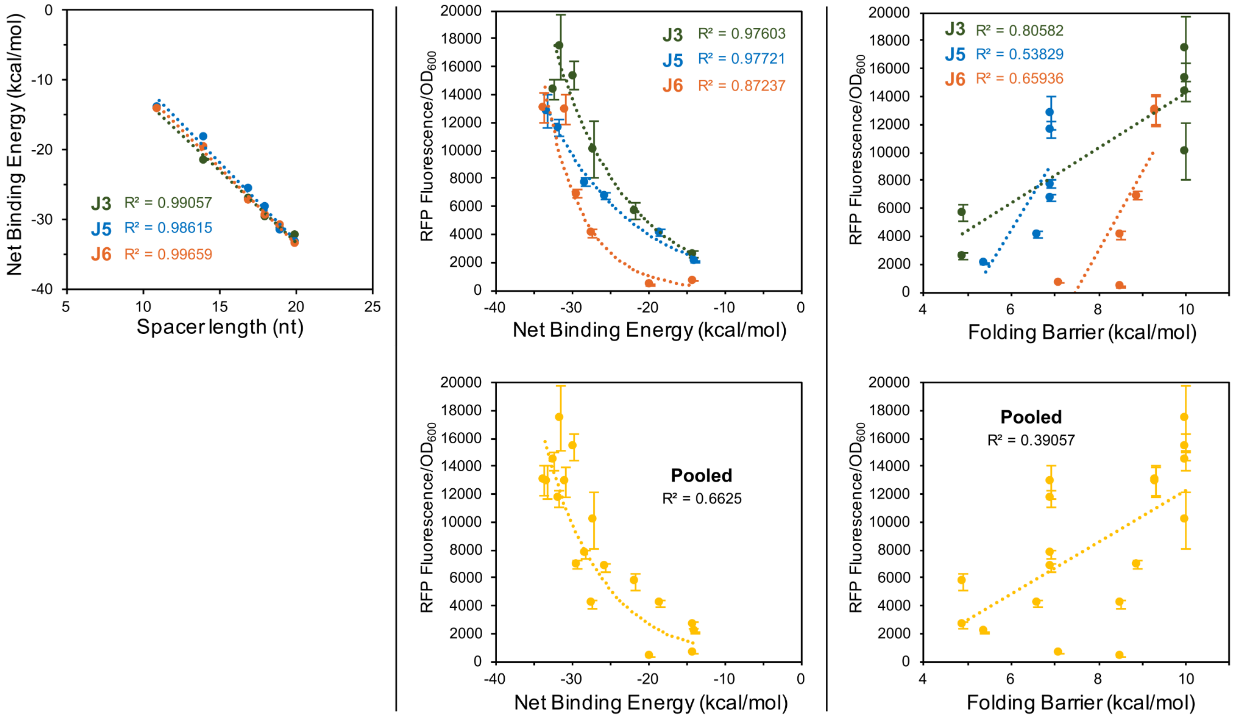


**Supplementary Figure 7. Parameter-output correlations in J3/J5/J6 truncation response.** Despite strong correlation between spacer length after truncation and Net Binding Energy (NBE) (left), correlation between NBE and RFP output from the J3, J5, and J6 promoters (center) is not strong enough for precise prediction of truncation effects on output. When individually fit to exponential functions (center top), the relatively sensitive truncation response of J6 results in a different function and a poorer correlation than that of J3 or J5. This means that when all the points are pooled into one correlation (center bottom), the overall correlation is poor. For the purpose of forward scRNA design, strong predictive power of a computational parameter would have come from a good correlation between that parameter and output, across all tested promoters. Therefore, our ability to predict spacer truncation effects on output remains poor, even though NBE correlates well with spacer length. Folding Barrier (FB) response to truncation (right) is much more stepwise than the NBE response, and does not correlate to output in a useful way. In the absence of computational prediction of truncation response, experimental mapping of that response (as in Figure 3c) remains necessary. *y*-axis values represent the average ± standard deviation calculated from $n$ = 3 biologically independent samples. Source data are provided as a Source Data file.

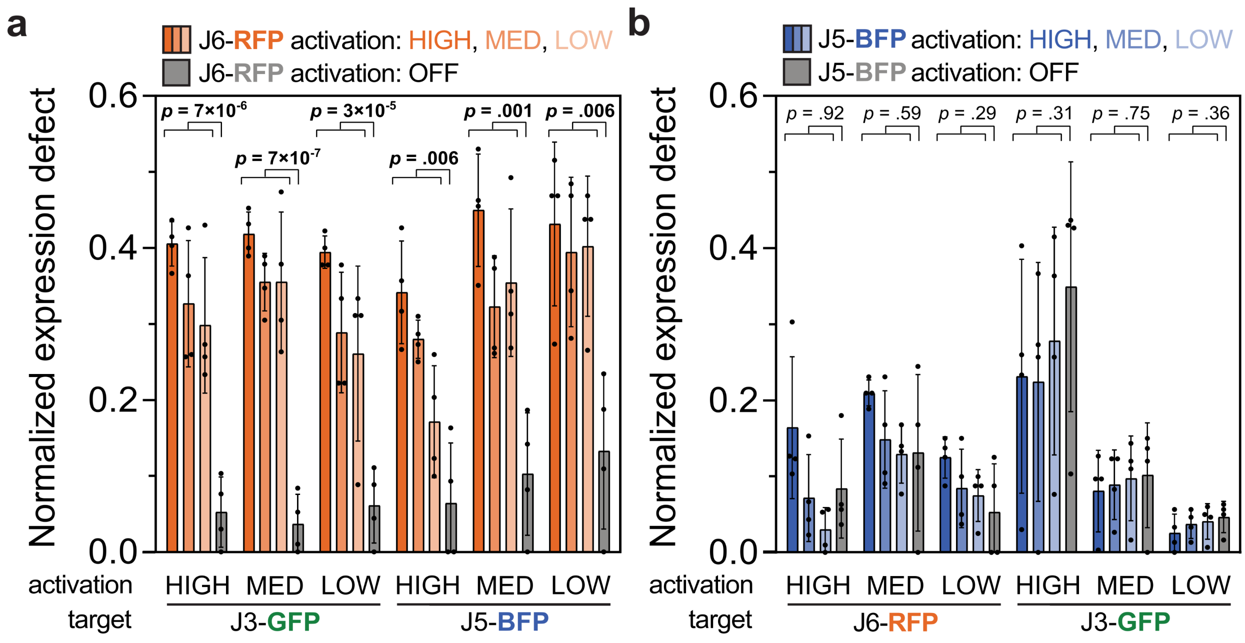


**Supplementary Figure 8. RFP expression burden affects simultaneous expression of GFP and BFP.** **a** Defect in GFP and BFP expression at noted RFP expression levels in strains expressing the combinatorial multi-scRNA library. Defect is calculated as the reduction in fluorescence within the activation state noted on the *x*-axis (e.g. high, medium, low expression), relative to the strain with the maximum fluorescence for that state. It is a quantification of the difference between expected fluorescence (dotted lines in Figure 4B) and measured fluorescence (bars in Figure 4B). Defects in GFP and BFP expression tend to be significantly higher when RFP is activated (red) than when RFP is not activated (grey), suggesting that RFP has more of an effect on overall expression burden than the other outputs. **b** Defect in RFP and GFP expression at noted BFP expression levels in strains expressing the combinatorial multi-scRNA library. Compared to the RFP effect in panel **a**, BFP activation has little effect on defect in RFP or GFP expression: J5-OFF strains (grey) have as much defect as J5-activated strains (blue). Each point is a ratio of two averages calculated from $n$ = 3 biologically independent samples. Bar values represent the average ± standard deviation calculated from 4 such ratios, because there are four activation states for each bar that are not specified by the *x*-axis and legend. Each *p*-value is derived from comparison between the 12 activated strains and the 4 unactivated strains contained in each *x*-axis label. Source data are provided as a Source Data file.

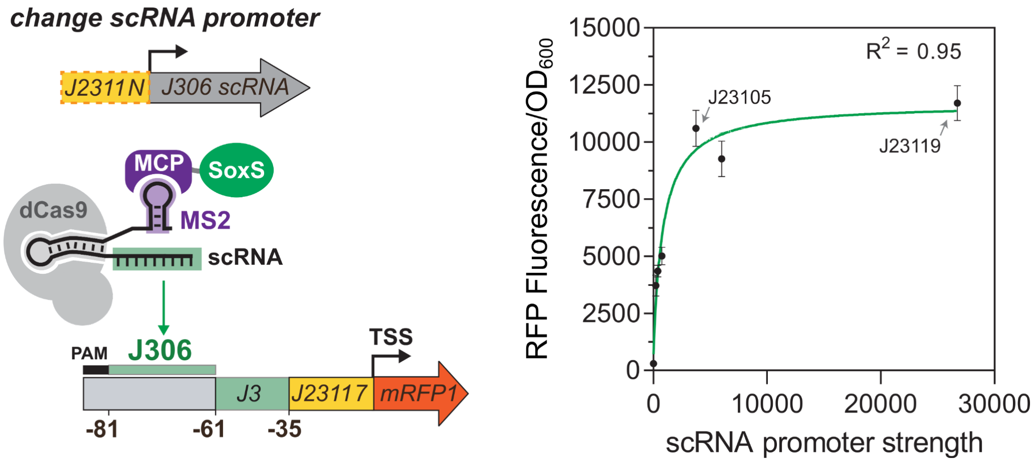


**Supplementary Figure 9. Activation dependence on scRNA expression level.** CRISPR-activated fluorescence resulting from a panel of six promoter strengths directing scRNA expression was compared scRNA expression level. All RFP expression was directed by the J3 synthetic promoter; fluorescence changes result from various amounts of J306 scRNA. Reducing scRNA expression by ~7-fold resulted in almost no defect in CRISPRa activity compared to the stronger, commonly-used J23119 promoter. Especially in multiple-scRNA systems, use of J23119 could therefore represent wasted expression capacity. Below the amount of scRNA directed by the J23105 promoter, output fluorescence from J3 drops dramatically, so J23105 was chosen as the minimum promoter strength directing expression of the triple-scRNA library. Promoter strength on the *x*-axis is quantified by the RFP fluorescence observed if that promoter was directly expressing mRFP1. These values are derived from Figure 3A in a previous report[1]. Green line represents a dose-response function fit to the data. Values on the *y*-axis represent the average ± standard deviation calculated from $n$ = 3 biologically independent samples. Source data are provided as a Source Data file.

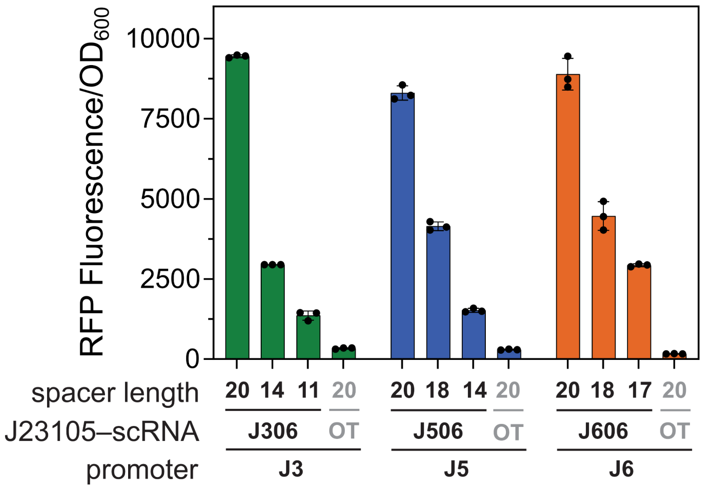


**Supplementary Figure 10. Effects of scRNA expression level on truncation-based tuning.** Adopting the J23105 promoter for expressing scRNAs in the triple-scRNA library format, here we verify that tuning CRISPR activation by truncating spacer lengths still results in the previously calibrated RFP output response (Figure 3c). Only truncation levels defined as high, medium, and low for the combinatorial scRNA library are included here. Bars represent RFP Fluorescence/$OD_{600}$ of strains harboring the J3, J5, or J6 synthetic promoters and expressing the J306, J506, and J606 scRNAs truncated to different lengths. Grey bars represent the baseline expression of the promoters and were obtained by measuring the Fluorescence/$OD_{600}$ of strains harboring each promoter and expressing an off-target scRNA. Values represent the average ± standard deviation calculated from $n$ = 3 biologically independent samples. Source data are provided as a Source Data file. The full sequence of the J3, J5, and J6 promoters is described in the Supplementary Methods.

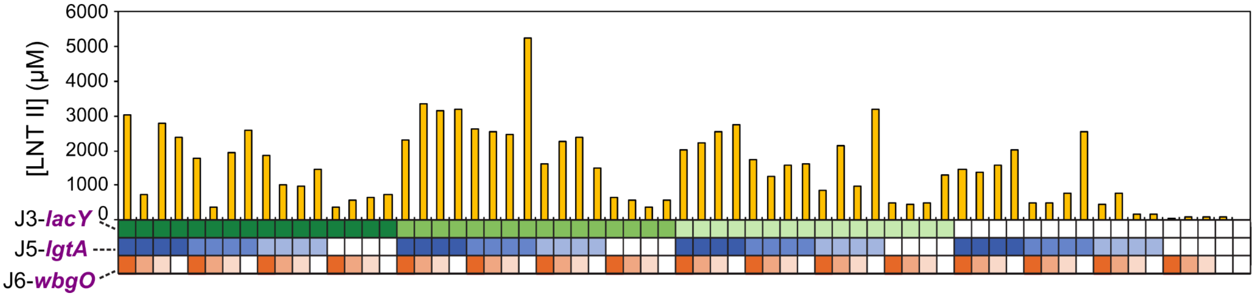


**Supplementary Figure 11. LNT II titers across the library.** HPLC analysis of supernatant from cultured library members indicates LNT II production levels from the scRNA combinations. The high LNT II titers relative to LNT titers, with relatively little dependence on *wbgO* expression, suggest galactosyltransferase activity as a limiting factor in the pathway. This bottleneck results in accumulation of LNT II and export to the supernatant, where it is inaccessible to WbgO. The *x*-axis heatmap is color coded to indicate the encoded promoter expression for each strain, as in Figure 6B. The 65th strain at right is a no-pathway control culture carrying an empty vector. Refer to Figure 6A for the pathway overview. Source data are provided as a Source Data file.

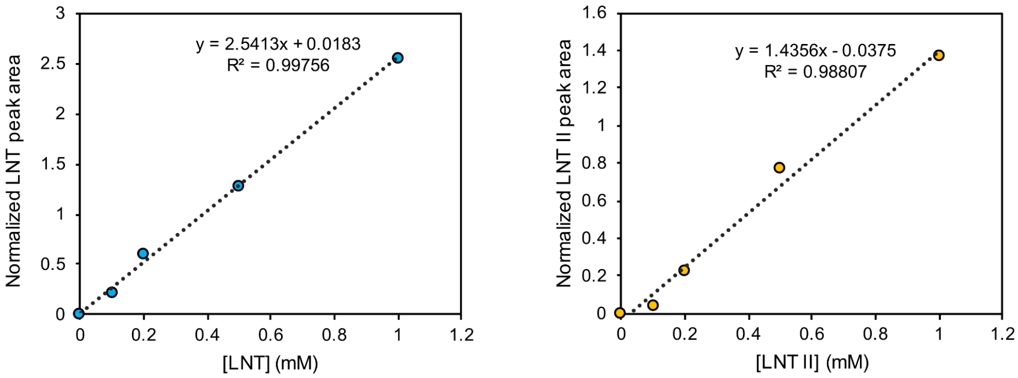


**Supplementary Figure 12. Standard curves for HPLC quantification of LNT and LNT II.** Peak areas resulting from LNT (10.6 min) or LNT II (11.4 min) were normalized by an endogenous peak (9.1 min) before conversion to molarity.

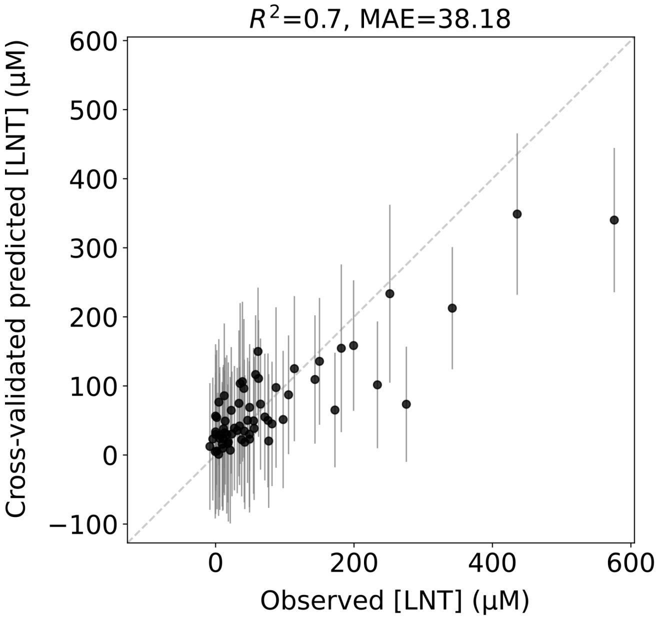


**Supplementary Figure 13. ART training and cross-validation for predicting LNT production.** Experimental observations of LNT production from the combinatorial scRNA library (*x*-axis) were used to train the Automated Recommendation Tool (ART) model. Here, we performed 10-fold cross-validation and plotted the predicted LNT production of each data point when it was in the test set (*y*-axis). When trained in this way, ART predictions correlate well with experimental values ($R^2$ = 0.7) for unseen training data. ART tended to underpredict the titer of the highest performing strains. Retraining on the full dataset was then performed before ART returned strain recommendations in the form of a set of input scRNA spacer lengths and a predicted LNT titer. Error bars indicate the 95% credible interval of the predictive posterior distribution. Source data are provided as a Source Data file.

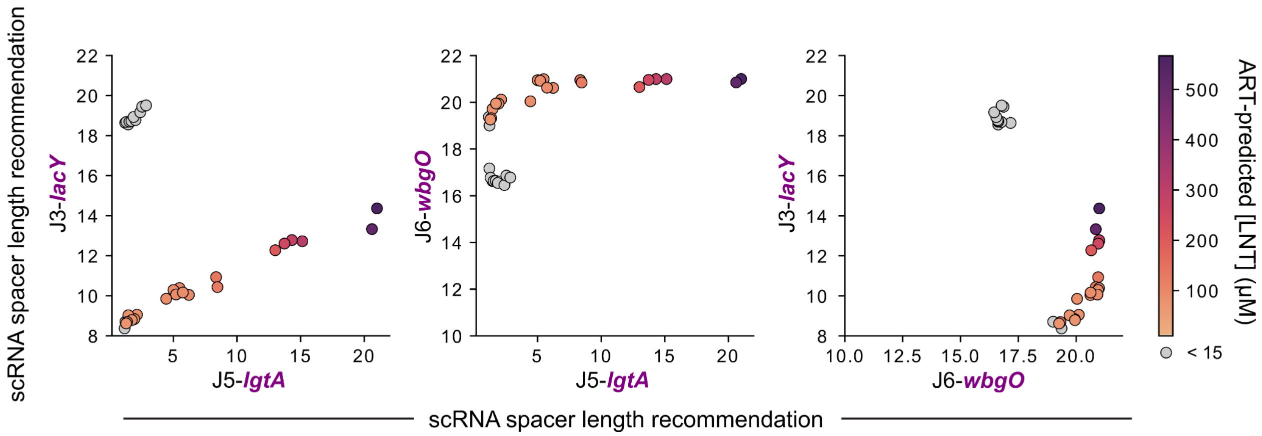


**Supplementary Figure 14. Scatter plots of ART recommendation combinations.** Each strain recommendation from the computational ART analysis in Figure 6d is plotted to demonstrate the relationship between each channel's spacer length recommendation. As in Figure 6d, the 20 strains with highest predicted LNT titer are highlighted in color on each sub-graph, while the 12 with lowest predicted LNT titer are shown in grey. Points are colored by their predicted LNT concentration: the same predictions as on the *y*-axis in Figure 6d. Source data are provided as a Source Data file.

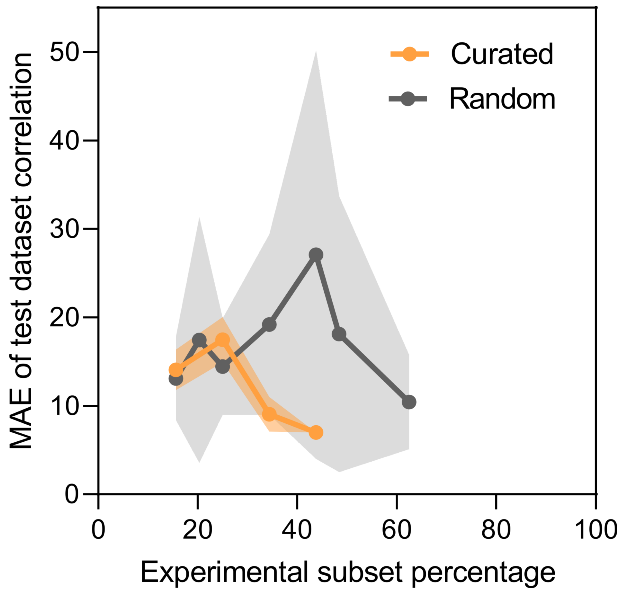


**Supplementary Figure 15. Degree of experimental sampling influences correlation between observed and predicted LNT titer.** Within the full experimental design space of 64 strains, ART predictions present the possibility of experimentally testing only a subset of the design space. The size of such subsets affects the accuracy of modeled production from the rest of the design space, presenting a tradeoff between experimental effort and prediction accuracy. Smaller subsets could result in more uncertainty in the model's predictions (higher

MAE), while larger subsets could provide more accurate predictions (lower MAE) at the cost of greater experimental effort. Using the LNT pathway, this trend was apparent in subsets consisting of a small number of specifically-selected strains (curated, orange) but not in randomly-selected subsets (grey). Strategic selection of the subset strain identities can also result in lower prediction uncertainty than most randomly-selected subsets. Beyond the constant inclusion of high-high-high, med-med-med, low-low-low, and OT-OT-OT, strain selection for the curated subsets drew randomly from a set of only the 24 strains that have non-duplicated inputs, e.g. medium-high-low but not medium-medium-low, instead of drawing from the whole set of 64 strains. MAE values are derived from observed vs. predicted correlation plots such as Supplementary Figure 13. This approach could potentially allow reasonably accurate model predictions with conveniently low experimental effort, although testing with other pathways should be undertaken. An optimal subset size depends on the complexity of the production landscape within the overall design space, and therefore on the complexity of the pathway itself. For the LNT pathway, for example, carefully choosing and experimentally testing only 22 (34%) of the 64 possible strains still results in reasonably low MAE, but subsets ≤25% result in higher MAE. We do not expect a particular subset size to be optimal for multiple different pathways, because production from different pathways will map differently onto the underlying design space. Different production landscapes will translate to different objective functions in the model, of variable complexity. More complex objective functions will require more training data to allow accurate predictions. Shading represents the standard deviation of MAE values from $n$ = 3 different subsets of each subset size. Source data and strain selections are provided as a Source Data file.

**SUPPLEMENTARY TABLES**

**Supplementary Table 1.** *E. coli* strains

| **Strain** | **Description** | **Genotype** |
|---|---|---|
| MG1655 | parent *E. coli* strain | F- λ- ilvG- rfb-50 rph-1 |
| JM109[6] | *lacZ*-deficient | recA1, endA1, gyrA96, thi, hsdR17, supE44, relA1, λ-, Δ(lac-proAB), [F' traD36, proAB, lacIqZΔM15] |

**Supplementary Table 2.** Select *E. coli* expression plasmids[a]

| Plasmid | Marker | Origin | Promoter | Gene | Terminator |
|---|---|---|---|---|---|
| pJF143.J3[b] | *AmpR* | *pSC101*** | J3 | *mRFP1* | BBa_B0015 |
| pJF143.J5 | *AmpR* | *pSC101*** | J5 | *mRFP1* | BBa_B0015 |
| pJF143.J6 | *AmpR* | *pSC101*** | J6 | *mRFP1* | BBa_B0015 |
| pJF185.x[c] | *CmR* | *p15A* | *1)* Sp.pCas9<br>*2)* J23107<br>*3)* J23119 | *1) dCas9*<br>*2) MCP-SoxS** [d]<br>*3)* scRNA[c] | *1)* BBa_B0015<br>*2)* BBa_B1002<br>*3)* ECK120033736[7] |
| pDSY23.x[b] | *AmpR* | *pSC101*** | J3 | *mRFP1* | BBa_B0015 |
| pDSY24.x[c] | *CmR* | *p15A* | *1)* Sp.pCas9<br>*2)* J23107<br>*3)* J23119 | *1) dCas9*<br>*2) MCP-SoxS** [d]<br>*3)* scRNA[c] | *1)* BBa_B0015<br>*2)* BBa_B1002<br>*3)* ECK120033736 |
| pJF144.J206 | *CmR* | *p15A* | *1)* Sp.pCas9<br>*2)* J23107<br>*3)* J23119 | *1) dCas9*<br>*2) MCP-SoxS**<br>*3)* J206 scRNA | *1)* BBa_B0015<br>*2)* BBa_B1002<br>*3)* TrrnB |
| pJF144.J306-n[e] | *CmR* | *p15A* | *1)* Sp.pCas9<br>*2)* J23107<br>*3)* J23119 | *1) dCas9*<br>*2) MCP-SoxS**<br>*3)* J306-n scRNA[e] | *1)* BBa_B0015<br>*2)* BBa_B1002<br>*3)* TrrnB |
| pJF144.J506-n[e] | *CmR* | *p15A* | *1)* Sp.pCas9<br>*2)* J23107<br>*3)* J23119 | *1) dCas9*<br>*2) MCP-SoxS**<br>*3)* J506-n scRNA[e] | *1)* BBa_B0015<br>*2)* BBa_B1002<br>*3)* TrrnB |
| pJF144.J606-n[e] | *CmR* | *p15A* | *1)* Sp.pCas9<br>*2)* J23107<br>*3)* J23119 | *1) dCas9*<br>*2) MCP-SoxS**<br>*3)* J606-n scRNA[e] | *1)* BBa_B0015<br>*2)* BBa_B1002<br>*3)* ECK120033736 |
| pJF144RS.J306-x[f] | *CmR* | *p15A* | *1)* Sp.pCas9<br>*2)* J23107<br>*3)* J23x[f] | *1) dCas9*<br>*2) MCP-SoxS*** [d]<br>*3)* J306 scRNA | *1)* BBa_B0015<br>*2)* BBa_B1002<br>*3)* ECK120033736 |
| pJF231.J306-n[g] | *CmR* | *p15A* | *1)* Sp.pCas9<br>*2)* J23107<br>*3)* J23105 | *1) dCas9*<br>*2) MCP-SoxS***<br>*3)* J306-n scRNA[g] | *1)* BBa_B0015<br>*2)* BBa_B1002<br>*3)* ECK120033736 |
| pJF231.J506-n[g] | *CmR* | *p15A* | *1)* Sp.pCas9<br>*2)* J23107<br>*3)* J23105 | *1) dCas9*<br>*2) MCP-SoxS***<br>*3)* J506-n scRNA[g] | *1)* BBa_B0015<br>*2)* BBa_B1002<br>*3)* ECK120033736 |

| | | | | | |
|---|---|---|---|---|---|
| pJF231.J606-n[g] | *CmR* | *p15A* | 1) Sp.pCas9<br>2) J23107<br>3) J23105 | 1) *dCas9*<br>2) *MCP-SoxS***<br>3) J606-n scRNA[g] | 1) BBa_B0015<br>2) BBa_B1002<br>3) ECK120033736 |
| pJF234.x | *CmR* | *p15A* | 1) Sp.pCas9<br>2) J23107<br>3) J23105<br>4) J23105<br>5) J23105 | 1) *dCas9*<br>2) *MCP-SoxS***<br>3) J306-n scRNA<br>4) J506-n scRNA<br>5) J606-n scRNA | 1) BBa_B0015<br>2) BBa_B1002<br>3) TrrnB<br>4) ECK120033736<br>5) ECK120033736 |
| pJF226C | *AmpR* | *pSC101*** | 1) J3<br>2) J5<br>3) J6 | 1) *sfgfp*<br>2) *mTag-BFP2*<br>3) *mRFP1* | 1) BBa_B0015<br>2) BBa_B0015<br>3) ECK120033736 |
| pCK200 | *AmpR* | *pSC101*** | 1) J3<br>2) J5<br>3) J6 | 1) *gtpch*<br>2) *ptps*<br>3) *sr* | 1) BBa_B0015<br>2) BBa_B0015<br>3) ECK120033736 |
| pARW15 | *AmpR* | *pSC101*** | 1) J3<br>2) J5<br>3) J6 | 1) *lacY*<br>2) *lgtA*<br>3) *wbgO* | 1) ECK120033737[7]<br>2) ECK120010818[7]<br>3) ECK120015440[7] |
| pIDF96C | *AmpR* | *pSC101*** | 1) J3<br>2) J5<br>3) J6 | 1) *lacY*<br>2) *lgtA*<br>3) *CvGalT* | 1) ECK120033737<br>2) ECK120010818<br>3) ECK120015440 |

[a] BBa sequences are from the Repository of Standard Biological Parts (http://parts.igem.org). dCas9 is the catalytically inactive form of *S. pyogenes* Cas9. Sp.pCas9 is the endogenous Cas9 promoter from *S. pyogenes*.

[b] Variants of pJF143.J3 in which the original J306 scRNA target site is replaced with the target site for the J106, the g1 to g13, or the A to W scRNAs (Supplementary Table 3) were used in Figure 2b and 2d. Variants of pDSY23, a derivative of pJF143.J3, in which the original J306 target site is replaced with the target site for the AA to EE scRNAs (Supplementary Table 3) were used in Supplementary Figure 5.

[c] Variants of the pJF185 plasmid in which the scRNA spacer sequence was g1 to g13 or A to W (Supplementary Table 3) were used in Figures 2b and 2d. Variants of the pDSY24 plasmid in which the scRNA spacer sequence was AA to EE (Supplementary Table 3) and with the variant dCas9-binding handle (Supplementary Table 4) were used in Supplementary Figure 5.

[d] To avoid widespread endogenous activation effects[1], SoxS includes either the single mutation R93A (denoted here by *) or double mutations R93A and S101A (denoted by **).
[e] Variants of the pJF144.J306, pJF144.J506 and pJF144.J606 plasmids where the J306, J506 and J606 spacers were n bases long, truncated from the 5' end, were used in Figure 3c (n = 20, 19, 18, 17, 14, 11, Supplementary Table 3).
[f] Variants of the pJF144RS.J306 plasmid where the promoter driving the scRNA was with BBa_J23105, BBa_J23106, BBa_J23112, BBa_J23114, BBa_J23117, or BBa_J23119 were used in Supplementary Figure 9.
[g] Variants of the pJF231.J306, pJF231.J506, and pJF231.J606 plasmids where the J306, J506 and J606 spacers were n bases long, truncated from the 5' end, were used in Figure 4b and Supplementary Figure 10 (for J306, n = 20, 14, 11; for J506, n = 20, 18, 14; for J606, n = 20, 18, 17).

**Supplementary Table 3.** scRNA spacer sequences

| scRNA target | DNA Sequence | Figures |
|---|---|---|
| J106[1,8] | AGGACGCCTTTGGTAACCGC | 2b, S1, S6 |
| J206[1,8] | TAGTAGCCGAACACGTCCTC | 2b, 3d |
| J306[1,8] | TTGTGTCCAGAACGCTCCGT | 2b, 2d, 3b, 3c, 4b, 5c, 6b-e, S1, S3-11 |
| g1 | GATCCTCATCCTGGCTTCGA | 2b, S1, S3, S4 |
| g2 | CGCAACTACTGGGGGTGTGA | 2b, S1, S3, S4 |
| g3 | TGTCTCTTGATGCAAGAGCG | 2b, S1, S3, S4, S6 |
| g4 | GTCCGCCACTTGATTTGTGT | 2b, S1, S3, S4 |
| g5 | TCGTGAATTCAGTGCCAACC | 2b, S1, S3, S4 |
| g6 | TCACACGCAGTTTCTCCCCT | 2b, S1, S3, S4 |
| g7 | GAACTCATGAAACTGATGTA | 2b, S1, S3, S4 |
| g8 | GCCTCCTCTTCTCTTTCTGC | 2b, S1, S3, S4 |
| g9 | CGAAGGGCATTAATGAGTTT | 2b, S1, S3, S4 |
| g10 | GGAGAAGGAGAAGGAAGAGT | 2b, S1, S3, S4 |
| g11 | CCAGCTAAACTCTAGTTCAA | 2b, S1, S3, S4 |
| g12 | AGAAGGTTATCTCAAGAAAC | 2b, S1, S3, S4 |
| g13 | TTTGAAATATTGACTTTTTA | 2b, S1, S3, S4 |
| A | AGCAACAGCGAGAACTCGTG | 2d, S3, S4 |
| B (J606) | GTCGCAGTCGCGCGAGCACT | 2d, 3b, 3c, 4b, 5c, 6b-e, S3, S4, S6-8, S10, S11 |
| C | AGCGCGATCGACTGATGCTC | 2d, S3, S4 |
| D | AGTAGCAGCACGCTAGACGC | 2d, S3, S4 |
| E (J506) | AGCAGCATGAGCAGCATTGA | 2d, 3b, 3c, 4b, 5c, 6b-e, S3, S4, S6-8, S10, S11 |

| | | |
|---|---|---|
| F | GTCGCTATGACGCTTCTCGC | 2d, S3, S4 |
| G | ACGAAGAGTGAGCGCAGATA | 2d, S3, S4 |
| H | ACAAGTATCGAAGCGAACTA | 4b, 5c, 6b, S8, S10 |
| I | GCGATAAACATATAAATTAT | 2d, S3, S4 |
| J | GAGACAGCTGAGAATGAGAA | 2d, S3, S4 |
| K | AGAGTAGACAGATTAGCAGT | 2d, S3, S4, S6 |
| L | GTGATAAACGGACTAGCCTT | 2d, S3, S4, S6 |
| M | AGCAGAAGTGTCAGCAGTGT | 2d, S3, S4 |
| N | GCAGTTACAGAAGTCGTCGC | 2d, S3, S4 |
| O | AGCAGCGTCGTCGAAAGTGT | 2d, S3, S4 |
| P | ATCGACAGAATCACTGCGCA | 2d, S3, S4 |
| Q | GCAGCGAAAGTCAGTGTCGT | 2d, S3, S4 |
| R | ACAGATGACGCTGCATAGCT | 2d, S3, S4 |
| S | GATGAGACAGACTAGTCGCG | 2d, S3, S4 |
| T | GATGACGCTGACAAGTGTTC | 2d, S3, S4 |
| U | ACGATCAGCGAGACTAGCTC | 2d, S3, S4 |
| V | GATGACAGCGTGACTAGCTG | 2d, S3, S4 |
| W | GATGATAGCGTGACTAGCTG | 4b, 5c, 6b, S6, S8, S10 |
| W108 | GAAGATCCGGCCTGCAGCCA | 2d, S3, S4 |
| AA | ACAATTGTCGCAACATGATC | S5, S6 |
| BB | GGCATCAAAAACACACGATC | S5 |
| CC | GAGACCATCGTAAATCCATA | S5 |
| DD | GGCGCCAAAAGTATCTCCGT | S5 |
| EE | ATCGCCGCTGTAACATCCAT | S5 |

| | | |
|---|---|---|
| J306-19 | TGTGTCCAGAACGCTCCGT | 3c |
| J306-18 | GTGTCCAGAACGCTCCGT | 3c |
| J306-17 | TGTCCAGAACGCTCCGT | 3c |
| J306-14 | CCAGAACGCTCCGT | 3c, 4b, 5c, 6b-e, S7, S10, S11 |
| J306-11 | GAACGCTCCGT | 3c, 4b, 5c, 6b, S7, S10, S11 |
| J506-19 | GCAGCATGAGCAGCATTGA | 3c |
| J506-18 | CAGCATGAGCAGCATTGA | 33c, 4b, 5c, 6b-e, S7, S10, S11 |
| J506-17 | AGCATGAGCAGCATTGA | 3c |
| J506-14 | ATGAGCAGCATTGA | 3c, 4b, 5c, 6b-e, S7, S10, S11 |
| J506-11 | ATGAGCAGCATTGA | 3c |
| J606-19 | TCGCAGTCGCGCGAGCACT | 3c |
| J606-18 | CGCAGTCGCGCGAGCACT | 3c, 4b, 5c, 6b-e, S7, S10, S11 |
| J606-17 | GCAGTCGCGCGAGCACT | 3c, 4b, 5c, 6b-e, S7, S10, S11 |
| J606-14 | GTCGCGCGAGCACT | 3c |
| J606-11 | GCGCGAGCACT | 3c |
| J119 | TGGTTGCCAAGGGTGACCTA | 4b, 5c, 6b, S8, S10 |
| J120 | AGGTCACCCTTGGCAACCAA | S10 |
| J118 | TGGTCACCATAGGTCACCCT | S10 |

**Supplementary Table 4.** scRNAs with orthogonal dCas9-binding handle sequences

| scRNA | RNA Sequence of Handle | Figure |
|---|---|---|
| Wild-type | GUUUUAGAGCUAGAAAUAGCAAGUUAAAAU | All except Suppl. 5 |
| AA – EE | GUUGGUGAGUUAGAAAUAACAAGUACCAAU | Suppl. 5 |

**Supplementary Table 5.** Select scRNA MFE structures (Supplementary Figure 6)

| scRNA | RNA sequence and predicted MFE structure[a] |
|---|---|
| J306 | UUGUGUCCAGAACGCUCCGUGUUUUAGAGCUAGAAAUAGCAAGUUAAAAUAAGGCUAGUCCGUUAUCAACUUGAAAAAGUGGCACAUGAGGAUCACCCAUGUGCUUUUUUU<br>.....((.(((((((...))))))).))((((....))))(((((...((((((.....)).)))).)))))..(((((.((((((((.((....)))))))))).))))) |
| J506 | AGCAGCAUGAGCAGCAUUGAGUUUUAGAGCUAGAAAUAGCAAGUUAAAAUAAGGCUAGUCCGUUAUCAACUUGAAAAAGUGGCACAUGAGGAUCACCCAUGUGCUUUUUUU<br>......(((.(((((.....((((((.((((....))))...))))))...))).)).))).....((((....))))((((((((.((....))))))))))...... |

| | |
|---|---|
| J606 | GUCGCAGUCGCGCGAGCACUGUUUUAGAGCUAGAAAUAGCAAGUUAAAAUAAGGCUAGUCCGUUAUCAACUUGAAAAAGUGGCACAUGAGGAUCACCCAUGUGCUUUUUUU<br>..(((......)))(((..((((((((.((((....))))...))))))))..)))...........((((....))))((((((((.((....))))))))))...... |
| AA | ACAAUUGUCGCAACAUGAUCGUUGGUGAGUUAGAAAUAACAAGUACCAAUAAGGCUAGUCCGUUAUCAACUUGAAAAAGUGGCACAUGAGGAUCACCCAUGUGCUUUUUUU<br>...................((((((.((((....))))...)))))))................((((....))))((((((((.((....))))))))))...... |
| g3 | UGUCUCUUGAUGCAAGAGCGGUUUUAGAGCUAGAAAUAGCAAGUUAAAAUAAGGCUAGUCCGUUAUCAACUUGAAAAAGUGGCACAUGAGGAUCACCCAUGUGCUUUUUUU<br>((.((((((...))))))))........((((....))))(((((...((((((.....)).)))).)))))..(((((.((((((((.((....)))))))))).))))) |
| J106 | AGGACGCCUUUGGUAACCGCGUUUUAGAGCUAGAAAUAGCAAGUUAAAAUAAGGCUAGUCCGUUAUCAACUUGAAAAAGUGGCACAUGAGGAUCACCCAUGUGCUUUUUUU<br>.((((((((...((((.((..(((........)))..))..))))....)))))..)))).......((((....))))((((((((.((....))))))))))...... |
| K | AGAGUAGACAGAUUAGCAGUGUUUUAGAGCUAGAAAUAGCAAGUUAAAAUAAGGCUAGUCCGUUAUCAACUUGAAAAAGUGGCACAUGAGGAUCACCCAUGUGCUUUUUUU<br>......(((.(((((((..((((((((.((((....))))...))))))))..))))))).)))....((((....))))((((((((.((....))))))))))...... |
| L | GUGAUAAACGGACUAGCCUUGUUUUAGAGCUAGAAAUAGCAAGUUAAAAUAAGGCUAGUCCGUUAUCAACUUGAAAAAGUGGCACAUGAGGAUCACCCAUGUGCUUUUUUU<br>.(((((.((((((((((((((((((.((((....))))...)))))))))))))))))))))))))((((....))))((((((((.((....))))))))))...... |
| W | GAUGAUAGCGUGACUAGCUGGUUUUAGAGCUAGAAAUAGCAAGUUAAAAUAAGGCUAGUCCGUUAUCAACUUGAAAAAGUGGCACAUGAGGAUCACCCAUGUGCUUUUUUU<br>..((((((((.((((((((.(((((((.((((....))))...)))))))..)))))))))))))))((((....))))((((((((.((....))))))))))...... |

[a] MFE structures predicted by the RNAfold WebServer and output in dot-bracket notation.

**Supplementary Table 6.** Select kinetic and thermodynamic parameters of scRNA truncations

| scRNA | Folding Barrier | Net Binding Energy |
|---|---|---|
| J306-20 | 10 | -32.3 |
| J306-19 | 10 | -31.4 |
| J306-18 | 10 | -29.8 |
| J306-17 | 10 | -27.1 |
| J306-14 | 4.9 | -21.7 |
| J306-11 | 4.9 | -14.2 |
| J506-20 | 6.9 | -33.3 |
| J506-19 | 6.9 | -31.7 |
| J506-18 | 6.9 | -28.3 |
| J506-17 | 6.9 | -25.7 |
| J506-14 | 6.6 | -18.4 |
| J506-11 | 5.4 | -14 |
| J606-20 | 9.3 | -33.6 |
| J606-19 | 9.3 | -30.9 |
| J606-18 | 8.9 | -29.4 |
| J606-17 | 8.5 | -27.4 |
| J606-14 | 8.5 | -19.8 |
| J606-11 | 7.1 | -14.2 |

### SUPPLEMENTARY METHODS

#### Computational analysis of scRNA activity (Supplementary Figures 3 and 4)

The following DNA sequence containing the CRISPRa target site was used as input to analyze scRNA activity using existing prediction models included with the CRISPOR analysis tool.

Upstream flanking sequence, variable target site, PAM site, downstream flanking sequence

CCCTAGGACTGAGCTAGCTGTCAATCTATAATCGCAACTTCAAGACGACGNNNNNNNNNNNNNNNNNNN NAGGAGAAGTGAGGAGACGAGCGAACGCGTCGTACGAGCTTTATGCATCTT

#### scRNA folding parameters for predicting effective CRISPR activation

Folding Barrier (FB) can be further divided into two separate barrier heights: the Handle Barrier and the Spacer Barrier (Supplementary Figure 2b). The Handle Barrier is defined as the barrier height for the conversion from the minimum free energy (MFE) structure to the most stable structure where the dCas9-binding handle is correctly folded. The Spacer Barrier represents the barrier height for the conversion from the most stable structure where the handle is correctly folded to the active structure. Within our dataset the Spacer Barrier explains substantially more of the variation in observed CRISPRa levels, suggesting that the ability of the dCas9-scRNA complex to bind to the DNA is a more important determinant of CRISPRa in our system than the dCas9 binding to an scRNA.

Overall, the kinetic parameter Folding Barrier has the most explanatory power of CRISPRa activity. To test the performance of Folding Barrier for predicting CRISPR-activated expression, we expanded our set of unique synthetic promoters for CRISPRa. We constructed 24 new CRISPRa promoters with target sites with Folding Barriers between about 5 kcal/mol and 35 kcal/mol. We chose these values in order to interrogate whether parameter values at the extremes of that range would enable us to identify highly active (or inactive) scRNAs that would be less likely to be identified by chance. Similar to the previous set where the scRNA target sites were randomly selected, we observed dramatic differences in CRISPRa activity with each promoter, varying by more than 40-fold (Figure 2d). The Folding Barrier parameter explained the majority of the variation in CRISPR-activated expression we observed, with a Spearman rank correlation coefficient ($r_S$) of 0.8 (Supplementary Figure 3). Interestingly, there appears to be a threshold—roughly 10 kcal/mol—below which reducing the Folding Barrier does not further increase CRISPRa levels.

To investigate if CRISPRa activity could be improved by further minimizing the Folding Barrier, we designed five additional scRNAs with Folding Barrier values lower than 3.5 kcal/mol. In order to design these highly-unstructured scRNAs, we had to mutate the dCas9-binding handle sequence to increase its folding stability: our computational predictions suggested that the stability of the wild-type handle was insufficient. For three of the five scRNAs, the MFE structure was the same as the active structure, implying that no further structural rearrangement was needed to bind to dCas9. CRISPR-activated gene expression using these scRNAs was generally high enough to indicate adequate activation, but was similar to the J306 (FB = 10 kcal/mol) level or slightly less (Supplementary Figure 5), suggesting a possible CRISPRa expression plateau in our system. This result further supports the idea of Folding Barrier's predictive power being most useful as a threshold rather than as a quantitative prediction. Below this threshold, other parameters are likely to have greater quantitative predictive power. These AA-EE scRNAs are not pictured in Figure 2d, but they further support the sigmoidal fit proposed there.

In addition to the kinetic parameters, we computed a set of metrics that analyze scRNA folding by relying solely on thermodynamic parameters. Net Binding Energy (NBE) is obtained by first calculating the ΔΔG between the MFE structure of a scRNA and its active conformation (Folding Energy), then adding the ΔG of the spacer binding the DNA target (Binding Energy), modeled here as an RNA-RNA duplex according to the Vienna tool RNAduplex (Supplementary Figure 2a). Net Binding Energy, then, is a composite metric combining the energy penalty associated with gRNA refolding with the energy of the spacer binding to its target.

The abilities of Folding Energy and Net Binding Energy to predict CRISPRa function were quite similar in our set of scRNAs (Supplementary Figure 3), suggesting that including Binding Energy adds little or no predictive value. Indeed, Binding Energy's correlation with CRISPRa activity is quite low. Still, Folding Energy has a high enough correlation to provide useful predictions, especially as a supplement to Folding Barrier below its 10 kcal/mol threshold.

To aid in identifying optimal scRNAs while avoiding defective ones, we additionally considered a combined screen using a kinetic parameter, Folding Barrier, and an energetic one, Net Binding Energy. The strategy was to first screen candidate scRNAs for those possessing sub-threshold FB values ≤ 10 kcal/mol, effectively screening against defective scRNAs that are kinetically trapped and unable to reach their active state. Remaining candidates would then be ranked based on their NBE value, enriching for optimal guides that possess ideal thermodynamic and kinetic properties (as in Supplementary Figure 4). We ultimately adopted the simpler use of only the FB screening for this work, which still represents an improvement over all of the common gRNA screening metrics we analyzed (Supplementary Figure 3), but the combined screening metric FB-NBE remains an effective strategy to simultaneously avoid defective guides and enrich for optimal guides.

Surprisingly, the handle fraction, which is the fraction of the population of scRNAs expected to have the Cas9-binding handle correctly folded, demonstrates no correlation with CRISPRa activity in our system (Supplementary Figure 3).

### Synthetic CRISPRa promoters

The synthetic CRISPRa promoters described in this study (J3, J5, and J6) are 170 bp in length and contain a PAM site for scRNA targeting at -81 bp to the TSS on the non-template strand, where maximum CRISPRa activity was observed in our previous study[8]. Each synthetic CRISPRa promoter contains a unique 20 bp scRNA target sequence and a unique 26 bp sequence between the target site and the minimal promoter. These sequences were previously characterized to give high CRISPR-activated expression (scRNA target/spacer sequences B and E and 26 bp sequences #24 and #25[1]). Sequences upstream of the -81 bp target site were randomized and verified to lack additional PAM sites. The BBa_J23117 minimal promoter was used for all three synthetic CRISPRa promoters, as it exhibited the highest dynamic range from the tested set of minimal promoters[1].

J3 promoter[1] (Figures 2-6 and Supplementary Figures 1, 3-5, and 7-11)

Upstream region, PAM site, J306 target site, unique 26 bp sequence (J3), BBa_J23117 promoter

AGCATTTGCGATCATTCACGCAGCGCTTATTCAGTTGCTCACTGCGATGTCATAATCATCGCTACGAGCTGTGAAAGATGCATAAAGCTCGTACGACGCGTTCGCTCGTCTCCTCACTTCTCCTACGGAGCGTTCTGGACACAACGTCGTCTTGAAGTTGCGATTATAGATTGACAGCTAGCTCAGTCCTAGGGATTGTGCTAGC

The experiments in Figure 2 and Supplementary Figures 1, 3, 4, and 5 were performed using a series of J3 promoter variants where the J306 target site is replaced with unique target sites (g1 through EE, Supplementary Table 3).

J5 promoter (Figures 3-6 and Supplementary Figures 7, 8, 10, and 11)

Upstream region, PAM site, J506 target site, unique 26 bp sequence (#25)[1], BBa_J23117 promoter

TATACATCGCATCACTACACTATTGATTATCATTGTGTACGTAACGAGCTTGCACAACGTGAAGTTCTTCGAGCACTTCAGCTCGCAACGTAAATGACAGTTGCTGTTAAGTGACGTGAATCCTTCAATGCTGCTCATGCTGCTGTCGTAAATAAGTAAGTCACTCCCACTTGACAGCTAGCTCAGTCCTAGGGATTGTGCTAGC

J6 promoter (Figures 3-6 and Supplementary Figures 7, 8, 10, and 11)

Upstream region, PAM site, J606 target site, unique 26 bp sequence (#24)[1], BBa_J23117 promoter

CTGCACGAGTTCGCTGTCGAGACAAGTCTCTTAGCGACGTATTACGAAGATCACATAGTCAGATGAAGCTATAGAGCACGACGCTAACGATTACGTCACGCTTGACACAACAGTTTCGCTACCTAGTGCTCGCGCGACTGCGACGTTGTCCTTCTAGTCGCCCATGACTCTTGACAGCTAGCTCAGTCCTAGGGATTGTGCTAGC

### Combinatorial CRISPRa library

**Supplementary Table 6.** scRNA library truncation levels

| Strain name | J306 spacer length (nt) | J3 activation level | J506 spacer length (nt) | J5 activation level | J606 spacer length (nt) | J6 activation level |
|---|---|---|---|---|---|---|
| 1 | 20 | high | 20 | high | 20 | high |
| 2 | 20 | high | 20 | high | 18 | medium |
| 3 | 20 | high | 20 | high | 17 | low |

| | | | | | | |
|---|---|---|---|---|---|---|
| 4 | 20 | high | 20 | high | OT | basal |
| 5 | 20 | high | 18 | medium | 20 | high |
| 6 | 20 | high | 18 | medium | 18 | medium |
| 7 | 20 | high | 18 | medium | 17 | low |
| 8 | 20 | high | 18 | medium | OT | basal |
| 9 | 20 | high | 14 | low | 20 | high |
| 10 | 20 | high | 14 | low | 18 | medium |
| 11 | 20 | high | 14 | low | 17 | low |
| 12 | 20 | high | 14 | low | OT | basal |
| 13 | 20 | high | OT | basal | 20 | high |
| 14 | 20 | high | OT | basal | 18 | medium |
| 15 | 20 | high | OT | basal | 17 | low |
| 16 | 20 | high | OT | basal | OT | basal |
| 17 | 14 | medium | 20 | high | 20 | high |
| 18 | 14 | medium | 20 | high | 18 | medium |
| 19 | 14 | medium | 20 | high | 17 | low |
| 20 | 14 | medium | 20 | high | OT | basal |
| 21 | 14 | medium | 18 | medium | 20 | high |
| 22 | 14 | medium | 18 | medium | 18 | medium |
| 23 | 14 | medium | 18 | medium | 17 | low |
| 24 | 14 | medium | 18 | medium | OT | basal |
| 25 | 14 | medium | 14 | low | 20 | high |
| 26 | 14 | medium | 14 | low | 18 | medium |
| 27 | 14 | medium | 14 | low | 17 | low |
| 28 | 14 | medium | 14 | low | OT | basal |
| 29 | 14 | medium | OT | basal | 20 | high |
| 30 | 14 | medium | OT | basal | 18 | medium |
| 31 | 14 | medium | OT | basal | 17 | low |
| 32 | 14 | medium | OT | basal | OT | basal |
| 33 | 11 | low | 20 | high | 20 | high |
| 34 | 11 | low | 20 | high | 18 | medium |
| 35 | 11 | low | 20 | high | 17 | low |
| 36 | 11 | low | 20 | high | OT | basal |
| 37 | 11 | low | 18 | medium | 20 | high |
| 38 | 11 | low | 18 | medium | 18 | medium |

| | | | | | | |
|---|---|---|---|---|---|---|
| 39 | 11 | low | 18 | medium | 17 | low |
| 40 | 11 | low | 18 | medium | OT | basal |
| 41 | 11 | low | 14 | low | 20 | high |
| 42 | 11 | low | 14 | low | 18 | medium |
| 43 | 11 | low | 14 | low | 17 | low |
| 44 | 11 | low | 14 | low | OT | basal |
| 45 | 11 | low | OT | basal | 20 | high |
| 46 | 11 | low | OT | basal | 18 | medium |
| 47 | 11 | low | OT | basal | 17 | low |
| 48 | 11 | low | OT | basal | OT | basal |
| 49 | OT | basal | 20 | high | 20 | high |
| 50 | OT | basal | 20 | high | 18 | medium |
| 51 | OT | basal | 20 | high | 17 | low |
| 52 | OT | basal | 20 | high | OT | basal |
| 53 | OT | basal | 18 | medium | 20 | high |
| 54 | OT | basal | 18 | medium | 18 | medium |
| 55 | OT | basal | 18 | medium | 17 | low |
| 56 | OT | basal | 18 | medium | OT | basal |
| 57 | OT | basal | 14 | low | 20 | high |
| 58 | OT | basal | 14 | low | 18 | medium |
| 59 | OT | basal | 14 | low | 17 | low |
| 60 | OT | basal | 14 | low | OT | basal |
| 61 | OT | basal | OT | basal | 20 | high |
| 62 | OT | basal | OT | basal | 18 | medium |
| 63 | OT | basal | OT | basal | 17 | low |
| 64 | OT | basal | OT | basal | OT | basal |

scRNA array sequence example: Strain #1

BBa_J23105 (SpeI) promoter, J306 spacer sequence, dCas9 handle, MS2 hairpin, rrnB terminator, BBa_J23105 (SpeI) promoter, J506 spacer sequence, dCas9 handle, MS2 hairpin, ECK120033736 terminator, BBa_J23105 (SpeI) promoter, J606 spacer sequence, dCas9 handle, MS2 hairpin, ECK120033736 terminator

TAAGTTGTTACTAGATTTACGGCTAGCTCAGTCCTAGGTACTATACTAGTTTGTGTCCAGAACGCTCCG
TGTTTTAGAGCTAGAAATAGCAAGTTAAAATAAGGCTAGTCCGTTATCAACTTGAAAAAGTGGCACATG

AGGATCACCCATGTGCTTTTTTTGAAGCTTGGGCCCGAACAAAAACTCATCTCAGAAGAGGATCTGAAT
AGCGCCGTCGACCATCATCATCATCATCATTGAGTTTAAACGGTCTCCAGCTTGGCTGTTTTGGCGGAT
GAGAGAAGATTTTCAGCCTGATACAGATTAAATCAGAACGCAGAAGCGGTCTGATAAAACAGAATTTGC
CTGGCGGCAGTAGCGCGGTGGTCCCACCTGACCCCATGCCGAACTCAGAAGTGAAACGCCGTAGCGCCG
ATGGTAGTGTGGGGTCTCCCCATGCGAGAGTAGGGAACTGCCAGGCATCAAATAAAACGAAAGGCTCAG
TCGAAAGACTGGGCCTTTCGTTTTATCTGTTGTTTGTCGGTGAACTGGATCCTTACAGATCATTTACGG
CTAGCTCAGTCCTAGGTACTATACTAGTAGCAGCATGAGCAGCATTGAGTTTTAGAGCTAGAAATAGCA
AGTTAAAATAAGGCTAGTCCGTTATCAACTTGAAAAAGTGGCACATGAGGATCACCCATGTGCTTTTTT
TAACGCATGAGAAAGCCCCCGGAAGATCACCTTCCGGGGGCTTTTTTATTGCGCAGATCATTTACGGCT
AGCTCAGTCCTAGGTACTATACTAGTGTCGCAGTCGCGCGAGCACTGTTTTAGAGCTAGAAATAGCAAG
TTAAAATAAGGCTAGTCCGTTATCAACTTGAAAAAGTGGCACATGAGGATCACCCATGTGCTTTTTTTA
ACGCATGAGAAAGCCCCCGGAAGATCACCTTCCGGGGGCTTTTTTATTGCGCAGATCTCCTTTGAGT

**Plasmid sequences for combinatorial CRISPRa experiments**

pJF226C (triple-fluorescent reporter: sfGFP, mTag-BFP2, mRFP1)

GAGCTCGTAAACTTGGTCTGACAGTTACCAATGCTTAATCAGTGAGGCACCTATCTCAGCGATCTGTCT
ATTTCGTTCATCCATAGTTGCCTGACTCCCCGTCGTGTAGATAACTACGATACGGGAGGGCTTACCATC
TGGCCCCAGTGCTGCAATGATACCGCGAGACCCACGCTCACCGGCTCCAGATTTATCAGCAATAAACCA
GCCAGCCGGAAGGGCCGAGCGCAGAAGTGGTCCTGCAACTTTATCCGCCTCCATCCAGTCTATTAATTG
TTGCCGGGAAGCTAGAGTAAGTAGTTCGCCAGTTAATAGTTTGCGCAACGTTGTTGCCATTGCTACAGG
CATCGTGGTGTCACGCTCGTCGTTTGGTATGGCTTCATTCAGCTCCGGTTCCCAACGATCAAGGCGAGT
TACATGATCCCCCATGTTGTGCAAAAAAGCGGTTAGCTCCTTCGGTCCTCCGATCGTTGTCAGAAGTAA
GTTGGCCGCAGTGTTATCACTCATGGTTATGGCAGCACTGCATAATTCTCTTACTGTCATGCCATCCGT
AAGATGCTTTTCTGTGACTGGTGAGTACTCAACCAAGTCATTCTGAGAATAGTGTATGCGGCGACCGAG
TTGCTCTTGCCCGGCGTCAATACGGGATAATACCGCGCCACATAGCAGAACTTTAAAAGTGCTCATCAT
TGGAAAACGTTCTTCGGGGCGAAAACTCTCAAGGATCTTACCGCTGTTGAGATCCAGTTCGATGTAACC
CACTCGTGCACCCAACTGATCTTCAGCATCTTTTACTTTCACCAGCGTTTCTGGGTGAGCAAAAACAGG
AAGGCAAAATGCCGCAAAAAAGGGAATAAGGGCGACACGGAAATGTTGAATACTCATACTCTTCCTTTT
TCAATATTATTGAAGCATTTATCAGGGTTATTGTCTCATGAGCGGATACATATTTGAATGTATTTAGAA
AAATAAACAAATAGGGGTTCCGCGCACATTTCCCCGAAAAGTGCCACCTGGGGCCCCTGCACGAGTTCG
CTGTCGAGACAAGTCTCTTAGCGACGTATTACGAAGATCACATAGTCAGATGAAGCTATAGAGCACGAC
GCTAACGATTACGTCACGCTTGACACAACAGTTTCGCTACCTAGTGCTCGCGCGACTGCGACGTTGTCC
TTCTAGTCGCCCATGACTCTTGACAGCTAGCTCAGTCCTAGGGATTGTGCTAGCGAATTCATTAAAGAG
GAGAAAGGTACCATGGCGAGTAGCGAAGACGTTATCAAAGAGTTCATGCGTTTCAAAGTTCGTATGGAA
GGTTCCGTTAACGGTCACGAGTTCGAAATCGAAGGTGAAGGTGAAGGTCGTCCGTACGAAGGTACCCAG
ACCGCTAAACTGAAAGTTACCAAAGGTGGTCCGCTGCCGTTCGCTTGGGACATCCTGTCCCCGCAGTTC
CAGTACGGTTCCAAAGCTTACGTTAAACACCCGGCTGACATCCCGGACTACCTGAAACTGTCCTTCCCG
GAAGGTTTCAAATGGGAACGTGTTATGAACTTCGAAGACGGTGGTGTTGTTACCGTTACCCAGGACTCC
TCCCTGCAAGACGGTGAGTTCATCTACAAAGTTAAACTGCGTGGTACCAACTTCCCGTCCGACGGTCCG
GTTATGCAGAAAAAAACCATGGGTTGGGAAGCTTCCACCGAACGTATGTACCCGGAAGACGGTGCTCTG
AAAGGTGAAATCAAAATGCGTCTGAAACTGAAAGACGGTGGTCACTACGACGCTGAAGTTAAAACCACC
TACATGGCTAAAAAACCGGTTCAGCTGCCGGGTGCTTACAAAACCGACATCAAACTGGACATCACCTCC

CACAACGAAGACTACACCATCGTTGAACAGTACGAACGTGCTGAAGGTCGTCACTCCACCGGTGCTTAA
GGATCCAAACTCGAGTAAGGATCTGTGCTTTTTTTAACGCATGAGAAAGCCCCCGGAAGATCACCTTCC
GGGGGCTTTTTTATTGCGCGGCCCTATAAACGCAGAAAGGCCCACCCGAAGGTGAGCCAGTGTGACTCT
AGTAGAGAGCGTTCACCGACAAACAACAGATAAAACGAAAGGCCCAGTCTTTCGACTGAGCCTTTCGTT
TTATTTGATGCCTGGATGCCTGGCAGTTCCCTACTCTCGCATGGGGAGACCCCACACTACCATCGGCGC
TACGGCGTTTCACTTCTGAGTTCGGCATGGGGTCAGGTGGGACCACCGCGCTACTGCCGCCAGGCAAAT
TCTGTTTTATCAGACCGCTTCTGCGTTCTGATTTAATCTGTATCAGGCTGAAAATCTTCTCTCATCCGC
CAAAACAGCCAAGCTTCGAATTCTTAATTAAGCTTGTGCCCCAGTTTGCTAGGGAGGTCGCAGTATCTG
GCCACTGCCACCTCGTGCTGCTCGACGTAGGTCTCGTTGTTGGCCTCCTTGATTCTTTCCAGTCTGTAG
TCCACATAGTAGACGCCAGGCATCTTGAGGTTCTTAGCGGGTTTCTTGGATCTATATGTGGTCTTGGCG
TTTGCGATCAGATGGCTCCCGCCCACGAGCTTCAGGGCCATGTCGTTTCTGCCTTCCAGGCCGCCGTCA
GCGGGGTACAGCGTCTCGGTGAAGGCCTCCCAGCCGAGTGTTTTCTTCTGCATCACAGGGCCGTTGGAT
GTGAAGTTCACCCCTCTGATCTTGACGTTGTAGATGAGGCAGCCGTCCTGGAGGCTGGTGTCCTGGGTA
GCGGTCAGCACGCCCCCGTCTTCGTATGTGGTGACTCTCTCCCATGTGAAGCCCTCAGGGAAGGACTGC
TTGAAGAAGTCGGGGATGCCCTGGGTGTGGTTGATGAAGGTCTTGCTGCCGTAGAGGAAGCTAGTAGCC
AGGATGTCGAAGGCGAAGGGGAGAGGGCCGCCCTCGACCACCTTGATTCTCATGGTCTGGGTGCCCTCG
TAGGGCTTGCCTTCGCCCTCGGATGTGCACTTGAAGTGATGGTTGTCCACGGTGCCCTCCATGTACAGC
TTCATGTGCATGTTCTCCTTAATCAGCTCGCTCATAGATCTGCCATGGTGATGGTGATGGTGGCTCCCC
ATGGTTAATTCCTCCTGTTAGCCCAAAAAACGGGTGCTAGCACAATCCCTAGGACTGAGCTAGCTGTCA
AGTGGGAGTGACTTACTTATTTACGACAGCAGCATGAGCAGCATTGAAGGATTCACGTCACTTAACAGC
AACTGTCATTTACGTTGCGAGCTGAAGTGCTCGAAGAACTTCACGTTGTGCAAGCTCGTTACGTACACA
ATGATAATCAATAGTGTAGTGATGCGATGTATAAGCATTTGCGATCATTCACGCAGCGCTTATTCAGTT
GCTCACTGCGATGTCATAATCATCGCTACGAGCTGTGAAAGATGCATAAAGCTCGTACGACGCGTTCGC
TCGTCTCCTCACTTCTCCTACGGAGCGTTCTGGACACAACGTCGTCTTGAAGTTGCGATTATAGATTGA
CAGCTAGCTCAGTCCTAGGGATTGTGCTAGCGAATTCATTAAAGAGGAGAAAGGTACCATGATGAGCAA
AGGAGAAGAACTTTTCACTGGAGTTGTCCCAATTCTTGTTGAATTAGATGGTGATGTTAATGGGCACAA
ATTTTCTGTCCGTGGAGAGGGTGAAGGTGATGCTACAAACGGAAAACTCACCCTTAAATTTATTTGCAC
TACTGGAAAACTACCTGTTCCGTGGCCAACACTTGTCACTACTCTGACCTATGGTGTTCAATGCTTTTC
CCGTTATCCGGATCACATGAAACGGCATGACTTTTTCAAGAGTGCCATGCCCGAAGGTTATGTACAGGA
ACGCACTATATCTTTCAAAGATGACGGGACCTACAAGACGCGTGCTGAAGTCAAGTTTGAAGGTGATAC
CCTTGTTAATCGTATCGAGTTAAAGGGTATTGATTTTAAAGAAGATGGAAACATTCTTGGACACAAACT
CGAGTACAACTTTAACTCACACAATGTATACATCACGGCAGACAAACAAAAGAATGGAATCAAAGCTAA
CTTCAAAATTCGCCACAACGTTGAAGATGGTTCCGTTCAACTAGCAGACCATTATCAACAAAATACTCC
AATTGGCGATGGCCCTGTCCTTTTACCAGACAACCATTACCTGTCGACACAATCTGTCCTTTCGAAAGA
TCCCAACGAAAAGCGTGACCACATGGTCCTTCTTGAGTTTGTAACTGCTGCTGGGATTACACATGGCAT
GGATGAGCTCTACAAATAAGGATCCAAACTCGAGTAAGGATCTCCAGGCATCAAATAAAACGAAAGGCT
CAGTCGAAAGACTGGGCCTTTCGTTTTATCTGTTGTTTGTCGGTGAACGCTCTCTACTAGAGTCACACT
GGCTCACCTTCGGGTGGGCCTTTCTGCGTTTATACCTAGGGTACGGGTTTTGCTGCCCGCAAACGGGCT
GTTCTGGTGTTGCTAGTTTGTTATCAGAATCGCAGATCCGGCTTCAGCCGGTTTGCCGGCTGAAAGCGC
TATTTCTTCCAGAATTGCCATGATTTTTTCCCCACGGGAGGCGTCACTGGCTCCCGTGTTGTCGGCAGC
TTTGATTCGATAAGCAGCATCGCCTGTTTCAGGCTGTCTATGTGTGACTGTTGAGCTGTAACAAGTTGT
CTCAGGTGTTCAATTTCATGTTCTAGTTGCTTTGTTTTACTGGTTTCACCTGTTCTATTAGGTGTTACA
TGCTGTTCATCTGTTACATTGTCGATCTGTTCATGGTGAACAGCTTTGAATGCACCAAAAACTCGTAAA
AGCTCTGATGTATCTATCTTTTTTACACCGTTTTCATCTGTGCATATGGACAGTTTTCCCTTTGATATG
TAACGGTGAACAGTTGTTCTACTTTTGTTTGTTAGTCTTGATGCTTCACTGATAGATACAAGAGCCATA
AGAACCTCAGATCCTTCCGTATTTAGCCAGTATGTTCTCTAGTGTGGTTCGTTGTTTTTGCGTGAGCCA
TGAGAACGAACCATTGAGATCATACTTACTTTGCATGTCACTCAAAAATTTTGCCTCAAAACTGGTGAG
CTGAATTTTTGCAGTTAAAGCATCGTGTAGTGTTTTTCTTAGTCCGTTATGTAGGTAGGAATCTGATGT
AATGGTTGTTGGTATTTTGTCACCATTCATTTTTATCTGGTTGTTCTCAAGTTCGGTTACGAGATCCAT
TTGTCTATCTAGTTCAACTTGGAAAATCAACGTATCAGTCGGGCGGCCTCGCTTATCAACCACCAATTT
CATATTGCTGTAAGTGTTTAAATCTTTACTTATTGGTTTCAAAACCCATTGGTTAAGCCTTTTAAACTC
ATGGTAGTTATTTTCAAGCATTAACATGAACTTAAATTCATCAAGGCTAATCTCTATATTTGCCTTGTG

AGTTTTCTTTTGTGTTAGTTCTTTTAATAACCACTCATAAATCCTCATAGAGTATTTGTTTTCAAAAGA
CTTAACATGTTCCAGATTATATTTTATGAATTTTTTTAACTGGAAAAGATAAGGCAATATCTCTTCACT
AAAAACTAATTCTAATTTTTCGCTTGAGAACTTGGCATAGTTTGTCCACTGGAAAATCTCAAAGCCTTT
AACCAAAGGATTCCTGATTTCCACAGTTCTCGTCATCAGCTCTCTGGTTGCTTTAGCTAATACACCATA
AGCATTTTCCCTACTGATGTTCATCATCTGAGCGTATTGGTTATAAGTGAACGATACCGTCCGTTCTTT
CCTTGTAGGGTTTTCAATCGTGGGGTTGAGTAGTGCCACACAGCATAAAATTAGCTTGGTTTCATGCTC
CGTTAAGTCATAGCGACTAATCGCTAGTTCATTTGCTTTGAAAACAACTAATTCAGACATACATCTCAA
TTGGTCTAGGTGATTTTAATCACTATACCAATTGAGATGGGCTAGTCAATGATAATTACTAGTCCTTTT
CCCGGGTGATCTGGGTATCTGTAAATTCTGCTAGACCTTTGCTGGAAAACTTGTAAATTCTGCTAGACC
CTCTGTAAATTCCGCTAGACCTTTGTGTGTTTTTTTTGTTTATATTCAAGTGGTTATAATTTATAGAAT
AAAGAAAGAATAAAAAAAGATAAAAAGAATAGATCCCAGCCCTGTGTATAACTCACTACTTTAGTCAGT
TCCGCAGTATTACAAAAGGATGTCGCAAACGCTGTTTGCTCCTCTACAAAACAGACCTTAAAACCCTAA
AGGCTTAAGTAGCACCCTCGCAAGCTCGGGCAAATCGCTGAATATTCCTTTTGTCTCCGACCATCAGGC
ACCTGAGTCGCTGTCTTTTTCGTGACATTCAGTTCGCTGCGCTCACGGCTCTGGCAGTGAATGGGGGTA
AATGGCACTACAGGCGCCTTTTATGGATTCATGCAAGAGATCTGAAACTACCCATAATACAAGAAAAGC
CCGTCACGGGCTTCTCAGGGCGTTTTATGGCGGGTCTGCTATGTGGTGCTATCTGACTTTTTGCTGTTC
AGCAGTTCCTGCCCTCTGATTTTCCAGTCTGACCACTTCGGATTATCCCGTGACAGGTCATTCAGACTG
GCTAATGCACCCAGTAAGGCAGCGGTATCATCAACAGGCTTACCCGTCTTACTGTCCCTAGTGCTTGGA
TTCTCACCAATAAAAAACGCCCGGCGGCAACCGAGCGTTCTGAACAAATCCAGATGGAGTTCTGAGGTC
ATTACTGGATCTATCAACAGGAGTCCAAGC

pCK200 (tetrahydrobiopterin pathway: *gtpch*, *ptps*, *sr*)

GAGCTCGTAAACTTGGTCTGACAGTTACCAATGCTTAATCAGTGAGGCACCTATCTCAGCGATCTGTCT
ATTTCGTTCATCCATAGTTGCCTGACTCCCCGTCGTGTAGATAACTACGATACGGGAGGGCTTACCATC
TGGCCCCAGTGCTGCAATGATACCGCGAGACCCACGCTCACCGGCTCCAGATTTATCAGCAATAAACCA
GCCAGCCGGAAGGGCCGAGCGCAGAAGTGGTCCTGCAACTTTATCCGCCTCCATCCAGTCTATTAATTG
TTGCCGGGAAGCTAGAGTAAGTAGTTCGCCAGTTAATAGTTTGCGCAACGTTGTTGCCATTGCTACAGG
CATCGTGGTGTCACGCTCGTCGTTTGGTATGGCTTCATTCAGCTCCGGTTCCCAACGATCAAGGCGAGT
TACATGATCCCCCATGTTGTGCAAAAAAGCGGTTAGCTCCTTCGGTCCTCCGATCGTTGTCAGAAGTAA
GTTGGCCGCAGTGTTATCACTCATGGTTATGGCAGCACTGCATAATTCTCTTACTGTCATGCCATCCGT
AAGATGCTTTTCTGTGACTGGTGAGTACTCAACCAAGTCATTCTGAGAATAGTGTATGCGGCGACCGAG
TTGCTCTTGCCCGGCGTCAATACGGGATAATACCGCGCCACATAGCAGAACTTTAAAAGTGCTCATCAT
TGGAAAACGTTCTTCGGGGCGAAAACTCTCAAGGATCTTACCGCTGTTGAGATCCAGTTCGATGTAACC
CACTCGTGCACCCAACTGATCTTCAGCATCTTTTACTTTCACCAGCGTTTCTGGGTGAGCAAAAACAGG
AAGGCAAAATGCCGCAAAAAAGGGAATAAGGGCGACACGGAAATGTTGAATACTCATACTCTTCCTTTT
TCAATATTATTGAAGCATTTATCAGGGTTATTGTCTCATGAGCGGATACATATTTGAATGTATTTAGAA
AAATAAACAAATAGGGGTTCCGCGCACATTTCCCCGAAAAGTGCCACCTGTGGCAATTCCGGGCCCTGC
ACGAGTTCGCTGTCGAGACAAGTCTCTTAGCGACGTATTACGAAGATCACATAGTCAGATGAAGCTATA
GAGCACGACGCTAACGATTACGTCACGCTTGACACAACAGTTTCGCTACCTAGTGCTCGCGCGACTGCG
ACGTTGTCCTTCTAGTCGCCCATGACTCTTGACAGCTAGCTCAGTCCTAGGGATTGTGCTAGCGAATTC
ATTAAAGAGGAGAAAGGTACCATGCATCACCATCACCACCATAGCAGTAAAGAACATCATTTGGTTATT
ATTAACGGTGTTAATAGAGGTTTCGGGCACTCCGTCGCGTTGGATTACATAAGACACTCAGGTGCTCAC
GCGGTGTCCTTTGTCTTGGTTGGTAGAACCCAGCATTCCTTGGAGCAAGTCTTAACGGAGCTGCACGAG
GCTGCATCCCACGCTGGTGTCGTCTTCAAGGGTGTCGTTGTGTCCGAGGTCGACCTGGCTCACTTGAAC
TCCCTCGACTCCAACCTCGCGAGGATACAGTCCGCCGCCGCTGACCTAAGAGACGAGGCGGCGCAAAGC
ACCAGAACTATCACTAAGTCGGTCCTCTTCAACAACGCGGGTAGCTTGGGTGACTTGTCCAAGACTGTT
AAGGAGTTCACCTGGCAAGAGGCTCGTTCCTACCTCGATTTCAACGTCGTGTCCCTCGTTGGTTTGTGC
TCCATGTTCTTGAAGGATACCCTCGAAGCATTCCCAAAGGAACAATACCCAGATCATAGAACTGTGGTC
GTGTCCATCTCTTCCCTATTAGCTGTTCAGGCTTTCCCAAACTGGGGTTTGTACGCTGCTGGTAAGGCA
GCTAGAGATAGACTATTAGGTGTTATTGCTCTCGAAGAAGCAGCTAATAACGTAAAGACCTTGAACTAC
GCTCCAGGTCCATTGGATAACGAAATGCAGGCTGACGTCCGCAGAACTTTGGGTGATAAGGAACAACTG

AAGATCTACGACGACATGCATAAGTCTGGTTCCTTGGTGAAGATGGAGGACTCCTCTAGAAAGTTGATT
CATTTGTTAAAGGCTGACACCTTCACCTCCGGTGGCCACATTGATTTCTACGACGAATAAGTGCTTTTT
TTAACGCATGAGAAAGCCCCCGGAAGATCACCTTCCGGGGGCTTTTTTATTGCGCGGCCCTATAAACGC
AGAAAGGCCCACCCGAAGGTGAGCCAGTGTGACTCTAGTAGAGAGCGTTCACCGACAAACAACAGATAA
AACGAAAGGCCCAGTCTTTCGACTGAGCCTTTCGTTTTATTTGATGCCTGGTTATTCACCTCTGTAAAC
GACAACGTTCTTGTCGGTTTCGTGCAACTTGATCTCGTACAAATCGTACGCGTCGGAAGGTGGCAAGTG
GCTCTTGATATTCTCCCACAAGAAGACAGCGAGGTTCTCAGTAGTGGAGGGTCTGGACTCGAAGTATGG
GACGTCTATATCCAAGTTTCTATGGTCACAAGGGTCCATGACAGCGACTTGCAAAGTCTTCTTAAGATC
GGTGATGTTGATGACCATGCCGGATTGTGGGTTGATCTGACCCTTGATGGTCACCTCGACCTTGTAGTT
GTGACCGTGACCGGAAGTGTGGTTGCACTTACCGAAGAGCTTGACGTTCTCAGCAGGCGAGAGGTGGAC
GGAGTTCAATCTGTGCGCAGCGGAGAAGTGTTCGATCCTGGTAACGTAAGCAGTTCTAACTGGAGTTGA
GGACGTATGGTGATGGTGGTGATGCATGGTACCTTTCTCCTCTTTAATGAATTCGCTAGCACAATCCCT
AGGACTGAGCTAGCTGTCAAGTGGGAGTGACTTACTTATTTACGACAGCAGCATGAGCAGCATTGAAGG
ATTCACGTCACTTAACAGCAACTGTCATTTACGTTGCGAGCTGAAGTGCTCGAAGAACTTCACGTTGTG
CAAGCTCGTTACGTACACAATGATAATCAATAGTGTAGTGATGCGATGTATACAGCATTTGCGATCATT
CACGCAGCGCTTATTCAGTTGCTCACTGCGATGTCATAATCATCGCTACGAGCTGTGAAAGATGCATAA
AGCTCGTACGACGCGTTCGCTCGTCTCCTCACTTCTCCTACGGAGCGTTCTGGACACAACGTCGTCTTG
AAGTTGCGATTATAGATTGACAGCTAGCTCAGTCCTAGGGATTGTGCTAGCGAATTCATTAAAGAGGAG
AAAGGTACCATGCATCACCATCACCATCACCCATCACTCAGTAAAGAAGCGGCCCTGGTTCATGAAGCG
TTAGTTGCGCGAGGACTGGAAACACCGCTGCGCCCGCCCGTGCATGAAATGGATAACGAAACGCGCAAA
AGCCTTATTGCTGGTCATATGACCGAAATCATGCAGCTGCTGAATCTCGACCTGGCTGATGACAGTTTG
ATGGAAACGCCGCATCGCATCGCTAAAATGTATGTCGATGAAATTTTCTCCGGTCTGGATTACGCCAAT
TTCCCGAAAATCACCCTCATTGAAAACAAAATGAAGGTCGATGAAATGGTCACCGTGCGCGATATCACT
CTGACCAGCACCTGTGAACACCATTTTGTTACCATCGATGGCAAAGCGACGGTGGCCTATATCCCGAAA
GATTCGGTGATCGGTCTGTCAAAAATTAACCGCATTGTGCAGTTCTTTGCCCAGCGTCCGCAGGTGCAG
GAACGTCTGACGCAGCAAATTCTTATTGCGCTACAAACGCTGCTGGGCACCAATAACGTGGCTGTCTCG
ATCGACGCGGTGCATTACTGCGTGAAGGCGCGTGGCATCCGCGATGCAACCAGTGCCACGACAACGACC
TCTCTTGGTGGATTGTTCAAATCCAGTCAGAATACGCGCCACGAGTTTCTGCGCGCTGTGCGTCATCAC
AACTAACCAGGCATCAAATAAAACGAAAGGCTCAGTCGAAAGACTGGGCCTTTCGTTTTATCTGTTGTT
TGTCGGTGAACGCTCTCTACTAGAGTCACACTGGCTCACCTTCGGGTGGGCCTTTCTGCGTTTATACCT
AGGGTACGGGTTTTGCTGCCCGCAAACGGGCTGTTCTGGTGTTGCTAGTTTGTTATCAGAATCGCAGAT
CCGGCTTCAGCCGGTTTGCCGGCTGAAAGCGCTATTTCTTCCAGAATTGCCATGATTTTTTCCCCACGG
GAGGCGTCACTGGCTCCCGTGTTGTCGGCAGCTTTGATTCGATAAGCAGCATCGCCTGTTTCAGGCTGT
CTATGTGTGACTGTTGAGCTGTAACAAGTTGTCTCAGGTGTTCAATTTCATGTTCTAGTTGCTTTGTTT
TACTGGTTTCACCTGTTCTATTAGGTGTTACATGCTGTTCATCTGTTACATTGTCGATCTGTTCATGGT
GAACAGCTTTGAATGCACCAAAAACTCGTAAAAGCTCTGATGTATCTATCTTTTTTACACCGTTTTCAT
CTGTGCATATGGACAGTTTTCCCTTTGATATGTAACGGTGAACAGTTGTTCTACTTTTGTTTGTTAGTC
TTGATGCTTCACTGATAGATACAAGAGCCATAAGAACCTCAGATCCTTCCGTATTTAGCCAGTATGTTC
TCTAGTGTGGTTCGTTGTTTTTGCGTGAGCCATGAGAACGAACCATTGAGATCATACTTACTTTGCATG
TCACTCAAAAATTTTGCCTCAAAACTGGTGAGCTGAATTTTTGCAGTTAAAGCATCGTGTAGTGTTTTT
CTTAGTCCGTTATGTAGGTAGGAATCTGATGTAATGGTTGTTGGTATTTTGTCACCATTCATTTTTATC
TGGTTGTTCTCAAGTTCGGTTACGAGATCCATTTGTCTATCTAGTTCAACTTGGAAAATCAACGTATCA
GTCGGGCGGCCTCGCTTATCAACCACCAATTTCATATTGCTGTAAGTGTTTAAATCTTTACTTATTGGT
TTCAAAACCCATTGGTTAAGCCTTTTAAACTCATGGTAGTTATTTTCAAGCATTAACATGAACTTAAAT
TCATCAAGGCTAATCTCTATATTTGCCTTGTGAGTTTTCTTTTGTGTTAGTTCTTTTAATAACCACTCA
TAAATCCTCATAGAGTATTTGTTTTCAAAAGACTTAACATGTTCCAGATTATATTTTATGAATTTTTTT
AACTGGAAAAGATAAGGCAATATCTCTTCACTAAAAACTAATTCTAATTTTTCGCTTGAGAACTTGGCA
TAGTTTGTCCACTGGAAAATCTCAAAGCCTTTAACCAAAGGATTCCTGATTTCCACAGTTCTCGTCATC
AGCTCTCTGGTTGCTTTAGCTAATACACCATAAGCATTTTCCCTACTGATGTTCATCATCTGAGCGTAT
TGGTTATAAGTGAACGATACCGTCCGTTCTTTCCTTGTAGGGTTTTCAATCGTGGGGTTGAGTAGTGCC
ACACAGCATAAAATTAGCTTGGTTTCATGCTCCGTTAAGTCATAGCGACTAATCGCTAGTTCATTTGCT
TTGAAAACAACTAATTCAGACATACATCTCAATTGGTCTAGGTGATTTTAATCACTATACCAATTGAGA

TGGGCTAGTCAATGATAATTACTAGTCCTTTTCCCGGGTGATCTGGGTATCTGTAAATTCTGCTAGACC
TTTGCTGGAAAACTTGTAAATTCTGCTAGACCCTCTGTAAATTCCGCTAGACCTTTGTGTGTTTTTTTT
GTTTATATTCAAGTGGTTATAATTTATAGAATAAAGAAAGAATAAAAAAAGATAAAAAGAATAGATCCC
AGCCCTGTGTATAACTCACTACTTTAGTCAGTTCCGCAGTATTACAAAAGGATGTCGCAAACGCTGTTT
GCTCCTCTACAAAACAGACCTTAAAACCCTAAAGGCTTAAGTAGCACCCTCGCAAGCTCGGGCAAATCG
CTGAATATTCCTTTTGTCTCCGACCATCAGGCACCTGAGTCGCTGTCTTTTTCGTGACATTCAGTTCGC
TGCGCTCACGGCTCTGGCAGTGAATGGGGGTAAATGGCACTACAGGCGCCTTTTATGGATTCATGCAAG
AGATCTGAAACTACCCATAATACAAGAAAAGCCCGTCACGGGCTTCTCAGGGCGTTTTATGGCGGGTCT
GCTATGTGGTGCTATCTGACTTTTTGCTGTTCAGCAGTTCCTGCCCTCTGATTTTCCAGTCTGACCACT
TCGGATTATCCCGTGACAGGTCATTCAGACTGGCTAATGCACCCAGTAAGGCAGCGGTATCATCAACAG
GCTTACCCGTCTTACTGTCCCTAGTGCTTGGATTCTCACCAATAAAAAACGCCCGGCGGCAACCGAGCG
TTCTGAACAAATCCAGATGGAGTTCTGAGGTCATTACTGGATCTATCAACAGGAGTCCAAGC

pARW15 (LNT pathway: *lacY*, *lgtA*, *wbgO*)

GAGCTCGTAAACTTGGTCTGACAGTTACCAATGCTTAATCAGTGAGGCACCTATCTCAGCGATCTGTCT
ATTTCGTTCATCCATAGTTGCCTGACTCCCCGTCGTGTAGATAACTACGATACGGGAGGGCTTACCATC
TGGCCCCAGTGCTGCAATGATACCGCGAGACCCACGCTCACCGGCTCCAGATTTATCAGCAATAAACCA
GCCAGCCGGAAGGGCCGAGCGCAGAAGTGGTCCTGCAACTTTATCCGCCTCCATCCAGTCTATTAATTG
TTGCCGGGAAGCTAGAGTAAGTAGTTCGCCAGTTAATAGTTTGCGCAACGTTGTTGCCATTGCTACAGG
CATCGTGGTGTCACGCTCGTCGTTTGGTATGGCTTCATTCAGCTCCGGTTCCCAACGATCAAGGCGAGT
TACATGATCCCCCATGTTGTGCAAAAAAGCGGTTAGCTCCTTCGGTCCTCCGATCGTTGTCAGAAGTAA
GTTGGCCGCAGTGTTATCACTCATGGTTATGGCAGCACTGCATAATTCTCTTACTGTCATGCCATCCGT
AAGATGCTTTTCTGTGACTGGTGAGTACTCAACCAAGTCATTCTGAGAATAGTGTATGCGGCGACCGAG
TTGCTCTTGCCCGGCGTCAATACGGGATAATACCGCGCCACATAGCAGAACTTTAAAAGTGCTCATCAT
TGGAAAACGTTCTTCGGGGCGAAAACTCTCAAGGATCTTACCGCTGTTGAGATCCAGTTCGATGTAACC
CACTCGTGCACCCAACTGATCTTCAGCATCTTTTACTTTCACCAGCGTTTCTGGGTGAGCAAAAACAGG
AAGGCAAAATGCCGCAAAAAAGGGAATAAGGGCGACACGGAAATGTTGAATACTCATACTCTTCCTTTT
TCAATATTATTGAAGCATTTATCAGGGTTATTGTCTCATGAGCGGATACATATTTGAATGTATTTAGAA
AAATAAACAAATAGGGGTTCCGCGCACATTTCCCCGAGGATCCAAAGTGCCACCTGTGGCAATTCCGAC
GTCGGGAAACACAGAAAAAAGCCCGCACCTGACAGTGCGGGCTTTTTTTTTCGACCAAAGGTTAAGCGA
CTTCATTCACCTGACGACGCAGCAGGGAAAGCGGGCCGGGGCCGCTAAGCGTGAACACGGAAATTAAGG
TGAAGCCCAGCGCCACCAGACCCAGCACCAGATAAGCGCCCTGGAAACCGATGCTTTCATACATATTGC
CCGCCAGTACAGACATAAAAATCATCGCCAGTTGCTTAAAGAAGCAGAAACAGACCAGATAAATCGTCG
CTGAAAAACGCACTTCAAACTGGCTGGTAATATATTTAAAGCAGCCCACCAGCAGGAACGGTACTTCAA
ACATATGCAGCGTTTTCAGAATAACCACTTCCAGCGCTGAGGTGGCGAACGATGAGCCAATAATACGTA
CAGACATAATAGTGCCAGCCAGCAGCAGGGCGTTTTTCCCACCGATGCGATTAATGATCAGTGGCGCAA
AGAACATAATCGAGGCGTTAAGTAATTCGCCCATTGTCGTTACGTAGCCAAATACCCGCGTACCCTGTT
CACCGGTAGCAAAGAACGAAGTAAAGAAATTAGCAAACTGTTGGTCAAAAACATCGTAGGTGCAGGAAA
CGCCAATAACATACAGTGACAAAAACCACAGTTTTGGCTGTCTGAACAGTTCCAGTGCCAGCTTAAGGC
TAAATGCCGAATGGTTGGCACCTACCGCATTGGCAACCGTGGCAGAAGAGGGCGCATCCGTTTTGGCGA
AAAAGAGTAAAACGGCGAGGATGAGTGCACAGCCAGAGCCCAGCCAGAAAACAAACTGATTATTGATGG
TGAACATGATGCCGACAATCGAGGCACACAGCGCCCAGCCAACACAGCCAAACATCCGCGCGCGACCAA
ATTCGAAATTACTGCGACGGCTGACTTTCTCAATAAATGCCTCTACTGCTGGCGCACCGGCGTTAAAAC
AAAAGCCTAGATAAATACCACCAACAATCGATCCTACTAAAATGTTGTATTGTAACAGTGGCCCGAAGA
TAAAAATAAAGAACGGCGCAAACATCACTAACATGCCGGTAATAATCCACAGCAGGTATTTGCGCAGCC
CGAGTTTGTCAGAAAGCAGACCAAACAGCGGTTGGAATAATAGCGAGAACAGAGAAATAGCGGCAAAAA
TAATACCCGTATCACTTTTGCTGATATGGTTGATGTCATGTAGCCAAATCGGGAAAAACGGGAAGTAGG
CTCCCATGATAAAAAAGTAAAAGAAAAAGAATAAACCGAACATCCAAAAGTTTGTGTTTTTTAAATAGT
ACATATGTATATCTCCTTCTTAAAAGATCTTTGCTAGCACAATCCCTAGGACTGAGCTAGCTGTCAATC
TATAATCGCAACTTCAAGACGACGTTGTGTCCAGAACGCTCCGTAGGAGAAGTGAGGAGACGAGCGAAC
GCGTCGTACGAGCTTTATGCATCTTTCACAGCTCGTAGCGATGATTATGACATCGCAGTGAGCAACTGA

ATAAGCGCTGCGTGAATGATCGCAAATGCTGACGTCGGAATTGCCACAGGTGGCACTTTTCGGGGAAAT
GTGCGCGTCTAGAATCGTAGCTAGCTGATCGTGCTAGTCGTATCGATCTAGCTCTCGAGTGGCAATTCC
GACGTCTGGCAATTCCGACGTCTATACATCGCATCACTACACTATTGATTATCATTGTGTACGTAACGA
GCTTGCACAACGTGAAGTTCTTCGAGCACTTCAGCTCGCAACGTAAATGACAGTTGCTGTTAAGTGACG
TGAATCCTTCAATGCTGCTCATGCTGCTGTCGTAAATAAGTAAGTCACTCCCACTTGACAGCTAGCTCA
GTCCTAGGGATTGTGCTAGCAACTGGTAATTTGAGGAGGTAATTTATGCCGTCGGAAGCCTTTCGTCGC
CATCGTGCGTACCGTGAAAACAAACTTCAGCCTTTGGTGTCTGTTTTGATCTGCGCGTATAATGTAGAG
AAGTATTTTGCACAGAGTCTGGCTGCGGTTGTGAACCAAACCTGGCGTAACTTAGACATCTTAATCGTC
GATGACGGAAGTACAGATGGCACGTTGGCGATCGCACAACGTTTCCAAGAGCAAGATGGTCGCATCCGT
ATCTTAGCCCAGCCACGCAATTCGGGACTTATCCCATCTTTGAACATTGGACTTGATGAACTGGCTAAA
AGTGGCATGGGAGAATACATTGCGCGCACTGATGCGGATGACATTGCAGCGCCCGACTGGATCGAGAAA
ATCGTGGGAGAGATGGAGAAAGATCGCTCTATCATCGCTATGGGAGCATGGTTAGAGGTTCTGTCAGAG
GAGAAGGACGGCAACCGTTTAGCTCGTCACCATGAGCATGGAAAAATCTGGAAAAAACCCACCCGCCAC
GAAGACATCGCCACGGTCTTCCCATTCGGTAACCCAATCCACAATAATACAATGATTATGCGTCGTAGC
GTGATTGATGGAGGTCTTCGCTACAACACTGAGCGCGATTGGGCCGAGGATTACCAGTTTTGGTATGAC
GTGAGCAAGTTAGGCCGTCTGGCTTACTATCCAGAAGCGCTGGTGAAGTACCGTCTTCATGCGAACCAG
GTGTCAAGTAAATACAGTGTACGTCAACATGAAATCGCACAGGGTATCCAAAAAACAGCGCGCAATGAT
TTTTTGCAGAGCATGGGTTTTAAAACACGCTTCGACAGCTTGGAGTATCGCCAAATCAAAGCCGTAGCC
TATGAATTACTGGAAAAGCATCTGCCGGAGGAAGATTTTGAGCGTGCCCGCCGCTTTTTATACCAATGT
TTTAAACGCACCGACACCCCACCAGCTGGCGCCTGGCTTGATTTTGCCGCCGACGGCCGTATGCGTCGC
TTATTTACTTTGCGCCAATATTTCGGTATCTTACGCCGTCTGTTGAAAAATCGCTGAGTCAGTTTCACC
TGTTTTACGTAAAAACCCGCTTCGGCGGGTTTTTACTTTTGGGAATTCATGTCTCTGCTCGACAGTGCA
CTAGCTGACTAGCTGACGCTCTGAGCTAGGCCTCCTGCACGAGTTCGCTGTCGAGACAAGTCTCTTAGC
GACGTATTACGAAGATCACATAGTCAGATGAAGCTATAGAGCACGACGCTAACGATTACGTCACGCTTG
ACACAACAGTTTCGCTACCTAGTGCTCGCGCGACTGCGACGTTGTCCTTCTAGTCGCCCATGACTCTTG
ACAGCTAGCTCAGTCCTAGGGATTGTGCTAGCAATCTCATAGATCAAATATAGGGGGGATCATATGATT
ATTGACGAGGCCGAGTCTGCAGAAAGCACACATCCGGTAGTATCTGTAATTCTGCCTGTAAATAAAAAA
AATCCGTTCTTGGATGAGGCGATTAACTCGATTTTGAGCCAAACCTTCAGCAGCTTTGAGATCATCATT
GTAGCGAATTGTTGCACGGATGACTTCTATAACGAGCTGAAACATAAAGTCAATGACAAAATCAAGCTG
ATCCGCACAAACATTGCTTACTTACCTTATTCTCTGAACAAAGCTATTGACTTGAGTAACGGTGAGTTC
ATCGCTCGTATGGATTCTGATGATATCTCCCACCCGGACCGCTTTACCAAGCAAGTAGATTTTTTAAAG
AATAATCCATACGTAGACGTTGTTGGGACAAACGCGATCTTTATTGATGACAAAGGACGCGAAATCAAC
AAAACCAAGTTACCCGAAGAAAATTTAGACATCGTGAAGAATCTTCCGTACAAATGCTGTATTGTCCAT
CCTTCTGTCATGTTTCGTAAGAAAGTAATTGCATCAATCGGCGGTTATATGTTTTCGAATTATTCTGAA
GATTACGAGCTTTGGAATCGTTTGTCCCTTGCTAAAATCAAATTCCAGAATCTTCCTGAATATCTGTTT
TATTACCGCCTTCATGAAGGCCAATCTACTGCAAAAAAGAACCTTTACATGGTGATGGTAAATGACCTT
GTCATCAAAATGAAGTGTTTCTTCCTTACGGGAAATATTAATTACTTATTTGGAGGGATTCGTACCATT
GCGAGTTTCATCTATTGCAAGTATATTAAATGAGGACTTCTACCTGGTCTCCGGCAATTAAAAAAGCGG
CTAACCACGCCGCTTTTTTTACGTCTGCACCTAGACGTACAGTCATCATGCGTACTGACTGACTGCCAT
GGCATCATGCTACTAGGGTACGGGTTTTGCTGCCCGCAAACGGGCTGTTCTGGTGTTGCTAGTTTGTTA
TCAGAATCGCAGATCCGGCTTCAGCCGGTTTGCCGGCTGAAAGCGCTATTTCTTCCAGAATTGCCATGA
TTTTTTCCCCACGGGAGGCGTCACTGGCTCCCGTGTTGTCGGCAGCTTTGATTCGATAAGCAGCATCGC
CTGTTTCAGGCTGTCTATGTGTGACTGTTGAGCTGTAACAAGTTGTCTCAGGTGTTCAATTTCATGTTC
TAGTTGCTTTGTTTTACTGGTTTCACCTGTTCTATTAGGTGTTACATGCTGTTCATCTGTTACATTGTC
GATCTGTTCATGGTGAACAGCTTTGAATGCACCAAAAACTCGTAAAAGCTCTGATGTATCTATCTTTTT
TACACCGTTTTCATCTGTGCATATGGACAGTTTTCCCTTTGATATGTAACGGTGAACAGTTGTTCTACT
TTTGTTTGTTAGTCTTGATGCTTCACTGATAGATACAAGAGCCATAAGAACCTCAGATCCTTCCGTATT
TAGCCAGTATGTTCTCTAGTGTGGTTCGTTGTTTTTGCGTGAGCCATGAGAACGAACCATTGAGATCAT
ACTTACTTTGCATGTCACTCAAAAATTTTGCCTCAAAACTGGTGAGCTGAATTTTTGCAGTTAAAGCAT
CGTGTAGTGTTTTTCTTAGTCCGTTATGTAGGTAGGAATCTGATGTAATGGTTGTTGGTATTTTGTCAC
CATTCATTTTTATCTGGTTGTTCTCAAGTTCGGTTACGAGATCCATTTGTCTATCTAGTTCAACTTGGA
AAATCAACGTATCAGTCGGGCGGCCTCGCTTATCAACCACCAATTTCATATTGCTGTAAGTGTTTAAAT

CTTTACTTATTGGTTTCAAAACCCATTGGTTAAGCCTTTTAAACTCATGGTAGTTATTTTCAAGCATTA
ACATGAACTTAAATTCATCAAGGCTAATCTCTATATTTGCCTTGTGAGTTTTCTTTTGTGTTAGTTCTT
TTAATAACCACTCATAAATCCTCATAGAGTATTTGTTTTCAAAAGACTTAACATGTTCCAGATTATATT
TTATGAATTTTTTTAACTGGAAAAGATAAGGCAATATCTCTTCACTAAAAACTAATTCTAATTTTTCGC
TTGAGAACTTGGCATAGTTTGTCCACTGGAAAATCTCAAAGCCTTTAACCAAAGGATTCCTGATTTCCA
CAGTTCTCGTCATCAGCTCTCTGGTTGCTTTAGCTAATACACCATAAGCATTTTCCCTACTGATGTTCA
TCATCTGAGCGTATTGGTTATAAGTGAACGATACCGTCCGTTCTTTCCTTGTAGGGTTTTCAATCGTGG
GGTTGAGTAGTGCCACACAGCATAAAATTAGCTTGGTTTCATGCTCCGTTAAGTCATAGCGACTAATCG
CTAGTTCATTTGCTTTGAAAACAACTAATTCAGACATACATCTCAATTGGTCTAGGTGATTTTAATCAC
TATACCAATTGAGATGGGCTAGTCAATGATAATTACTAGTCCTTTTCCCGGGTGATCTGGGTATCTGTA
AATTCTGCTAGACCTTTGCTGGAAAACTTGTAAATTCTGCTAGACCCTCTGTAAATTCCGCTAGACCTT
TGTGTGTTTTTTTTGTTTATATTCAAGTGGTTATAATTTATAGAATAAAGAAAGAATAAAAAAAGATAA
AAAGAATAGATCCCAGCCCTGTGTATAACTCACTACTTTAGTCAGTTCCGCAGTATTACAAAAGGATGT
CGCAAACGCTGTTTGCTCCTCTACAAAACAGACCTTAAAACCCTAAAGGCTTAAGTAGCACCCTCGCAA
GCTCGGGCAAATCGCTGAATATTCCTTTTGTCTCCGACCATCAGGCACCTGAGTCGCTGTCTTTTTCGT
GACATTCAGTTCGCTGCGCTCACGGCTCTGGCAGTGAATGGGGGTAAATGGCACTACAGGCGCCTTTTA
TGGATTCATGCAAGGAAACTACCCATAATACAAGAAAAGCCCGTCACGGGCTTCTCAGGGCGTTTTATG
GCGGGTCTGCTATGTGGTGCTATCTGACTTTTTGCTGTTCAGCAGTTCCTGCCCTCTGATTTTCCAGTC
TGACCACTTCGGATTATCCCGTGACAGGTCATTCAGACTGGCTAATGCACCCAGTAAGGCAGCGGTATC
ATCAACAGGCTTACCCGTCTTACTGTCCCTAGTGCTTGGATTCTCACCAATAAAAAACGCCCGGCGGCA
ACCGAGCGTTCTGAACAAATCCAGATGGAGTTCTGAGGTCATTACTGGATCTATCAACAGGAGTCCAAG
C

pIDF96C (LNT pathway: *lacY*, *lgtA*, *CvGalT*)

GAGCTCGTAAACTTGGTCTGACAGTTACCAATGCTTAATCAGTGAGGCACCTATCTCAGCGATCTGTCT
ATTTCGTTCATCCATAGTTGCCTGACTCCCCGTCGTGTAGATAACTACGATACGGGAGGGCTTACCATC
TGGCCCCAGTGCTGCAATGATACCGCGAGACCCACGCTCACCGGCTCCAGATTTATCAGCAATAAACCA
GCCAGCCGGAAGGGCCGAGCGCAGAAGTGGTCCTGCAACTTTATCCGCCTCCATCCAGTCTATTAATTG
TTGCCGGGAAGCTAGAGTAAGTAGTTCGCCAGTTAATAGTTTGCGCAACGTTGTTGCCATTGCTACAGG
CATCGTGGTGTCACGCTCGTCGTTTGGTATGGCTTCATTCAGCTCCGGTTCCCAACGATCAAGGCGAGT
TACATGATCCCCCATGTTGTGCAAAAAAGCGGTTAGCTCCTTCGGTCCTCCGATCGTTGTCAGAAGTAA
GTTGGCCGCAGTGTTATCACTCATGGTTATGGCAGCACTGCATAATTCTCTTACTGTCATGCCATCCGT
AAGATGCTTTTCTGTGACTGGTGAGTACTCAACCAAGTCATTCTGAGAATAGTGTATGCGGCGACCGAG
TTGCTCTTGCCCGGCGTCAATACGGGATAATACCGCGCCACATAGCAGAACTTTAAAAGTGCTCATCAT
TGGAAAACGTTCTTCGGGGCGAAAACTCTCAAGGATCTTACCGCTGTTGAGATCCAGTTCGATGTAACC
CACTCGTGCACCCAACTGATCTTCAGCATCTTTTACTTTCACCAGCGTTTCTGGGTGAGCAAAAACAGG
AAGGCAAAATGCCGCAAAAAAGGGAATAAGGGCGACACGGAAATGTTGAATACTCATACTCTTCCTTTT
TCAATATTATTGAAGCATTTATCAGGGTTATTGTCTCATGAGCGGATACATATTTGAATGTATTTAGAA
AAATAAACAAATAGGGGTTCCGCGCACATTTCCCCGAGGATCCAAAGTGCCACCTGTGGCAATTCCGAC
GTCGGGAAACACAGAAAAAAGCCCGCACCTGACAGTGCGGGCTTTTTTTTTCGACCAAAGGTTAAGCGA
CTTCATTCACCTGACGACGCAGCAGGGAAAGCGGGCCGGGGCCGCTAAGCGTGAACACGGAAATTAAGG
TGAAGCCCAGCGCCACCAGACCCAGCACCAGATAAGCGCCCTGGAAACCGATGCTTTCATACATATTGC
CCGCCAGTACAGACATAAAAATCATCGCCAGTTGCTTAAAGAAGCAGAAACAGACCAGATAAATCGTCG
CTGAAAAACGCACTTCAAACTGGCTGGTAATATATTTAAAGCAGCCCACCAGCAGGAACGGTACTTCAA
ACATATGCAGCGTTTTCAGAATAACCACTTCCAGCGCTGAGGTGGCGAACGATGAGCCAATAATACGTA
CAGACATAATAGTGCCAGCCAGCAGCAGGGCGTTTTTCCCACCGATGCGATTAATGATCAGTGGCGCAA
AGAACATAATCGAGGCGTTAAGTAATTCGCCCATTGTCGTTACGTAGCCAAATACCCGCGTACCCTGTT
CACCGGTAGCAAAGAACGAAGTAAAGAAATTAGCAAACTGTTGGTCAAAAACATCGTAGGTGCAGGAAA
CGCCAATAACATACAGTGACAAAAACCACAGTTTTGGCTGTCTGAACAGTTCCAGTGCCAGCTTAAGGC
TAAATGCCGAATGGTTGGCACCTACCGCATTGGCAACCGTGGCAGAAGAGGGCGCATCCGTTTTGGCGA
AAAAGAGTAAAACGGCGAGGATGAGTGCACAGCCAGAGCCCAGCCAGAAAACAAACTGATTATTGATGG

TGAACATGATGCCGACAATCGAGGCACACAGCGCCCAGCCAACACAGCCAAACATCCGCGCGCGACCAA
ATTCGAAATTACTGCGACGGCTGACTTTCTCAATAAATGCCTCTACTGCTGGCGCACCGGCGTTAAAAC
AAAAGCCTAGATAAATACCACCAACAATCGATCCTACTAAAATGTTGTATTGTAACAGTGGCCCGAAGA
TAAAAATAAAGAACGGCGCAAACATCACTAACATGCCGGTAATAATCCACAGCAGGTATTTGCGCAGCC
CGAGTTTGTCAGAAAGCAGACCAAACAGCGGTTGGAATAATAGCGAGAACAGAGAAATAGCGGCAAAAA
TAATACCCGTATCACTTTTGCTGATATGGTTGATGTCATGTAGCCAAATCGGGAAAAACGGGAAGTAGG
CTCCCATGATAAAAAAGTAAAAGAAAAAGAATAAACCGAACATCCAAAAGTTTGTGTTTTTTAAATAGT
ACATATGTATATCTCCTTCTTAAAAGATCTTTGCTAGCACAATCCCTAGGACTGAGCTAGCTGTCAATC
TATAATCGCAACTTCAAGACGACGTTGTGTCCAGAACGCTCCGTAGGAGAAGTGAGGAGACGAGCGAAC
GCGTCGTACGAGCTTTATGCATCTTTCACAGCTCGTAGCGATGATTATGACATCGCAGTGAGCAACTGA
ATAAGCGCTGCGTGAATGATCGCAAATGCTGACGTCGGAATTGCCACAGGTGGCACTTTTCGGGGAAAT
GTGCGCGTCTAGAATCGTAGCTAGCTGATCGTGCTAGTCGTATCGATCTAGCTCTCGAGTGGCAATTCC
GACGTCTGGCAATTCCGACGTCTATACATCGCATCACTACACTATTGATTATCATTGTGTACGTAACGA
GCTTGCACAACGTGAAGTTCTTCGAGCACTTCAGCTCGCAACGTAAATGACAGTTGCTGTTAAGTGACG
TGAATCCTTCAATGCTGCTCATGCTGCTGTCGTAAATAAGTAAGTCACTCCCACTTGACAGCTAGCTCA
GTCCTAGGGATTGTGCTAGCAACTGGTAATTTGAGGAGGTAATTTATGCCGTCGGAAGCCTTTCGTCGC
CATCGTGCGTACCGTGAAAACAAACTTCAGCCTTTGGTGTCTGTTTTGATCTGCGCGTATAATGTAGAG
AAGTATTTTGCACAGAGTCTGGCTGCGGTTGTGAACCAAACCTGGCGTAACTTAGACATCTTAATCGTC
GATGACGGAAGTACAGATGGCACGTTGGCGATCGCACAACGTTTCCAAGAGCAAGATGGTCGCATCCGT
ATCTTAGCCCAGCCACGCAATTCGGGACTTATCCCATCTTTGAACATTGGACTTGATGAACTGGCTAAA
AGTGGCATGGGAGAATACATTGCGCGCACTGATGCGGATGACATTGCAGCGCCCGACTGGATCGAGAAA
ATCGTGGGAGAGATGGAGAAAGATCGCTCTATCATCGCTATGGGAGCATGGTTAGAGGTTCTGTCAGAG
GAGAAGGACGGCAACCGTTTAGCTCGTCACCATGAGCATGGAAAAATCTGGAAAAAACCCACCCGCCAC
GAAGACATCGCCACGGTCTTCCCATTCGGTAACCCAATCCACAATAATACAATGATTATGCGTCGTAGC
GTGATTGATGGAGGTCTTCGCTACAACACTGAGCGCGATTGGGCCGAGGATTACCAGTTTTGGTATGAC
GTGAGCAAGTTAGGCCGTCTGGCTTACTATCCAGAAGCGCTGGTGAAGTACCGTCTTCATGCGAACCAG
GTGTCAAGTAAATACAGTGTACGTCAACATGAAATCGCACAGGGTATCCAAAAAACAGCGCGCAATGAT
TTTTTGCAGAGCATGGGTTTTAAAACACGCTTCGACAGCTTGGAGTATCGCCAAATCAAAGCCGTAGCC
TATGAATTACTGGAAAAGCATCTGCCGGAGGAAGATTTTGAGCGTGCCCGCCGCTTTTTATACCAATGT
TTTAAACGCACCGACACCCCACCAGCTGGCGCCTGGCTTGATTTTGCCGCCGACGGCCGTATGCGTCGC
TTATTTACTTTGCGCCAATATTTCGGTATCTTACGCCGTCTGTTGAAAAATCGCTGAGTCAGTTTCACC
TGTTTTACGTAAAAACCCGCTTCGGCGGGTTTTTACTTTTGGGAATTCATGTCTCTGCTCGACAGTGCA
CTAGCTGACTAGCTGACGCTCTGAGCTAGGCCTCCTGCACGAGTTCGCTGTCGAGACAAGTCTCTTAGC
GACGTATTACGAAGATCACATAGTCAGATGAAGCTATAGAGCACGACGCTAACGATTACGTCACGCTTG
ACACAACAGTTTCGCTACCTAGTGCTCGCGCGACTGCGACGTTGTCCTTCTAGTCGCCCATGACTCTTG
ACAGCTAGCTCAGTCCTAGGGATTGTGCTAGCAATCTCATAAATCAAATATAGGGAGGATCATATGGAC
ACCATCATGATTAAACGTCCGCTGGTTAGCGTTATTCTGCCGGTGAATAAAAACAATCCGCATCTGGAA
GAAGCAATCCAGAGCATTAAAAACCAGACCTATAAAGAGCTGGAACTGATCATTATTGCCAACAACTGC
GAGGATAACTTTTATAGCCTGCTGCTGAAATATCAGGACCAGAAAACCAAAATTATCCGCACCAGCATC
AAATATCTGCCGTTTAGCCTGAATCTGGGTGTTCATCTGAGCCAGGGTGAATATATTGCACGTATGGAT
TCAGATGATATCAGCGTTCTGGATCGCATTGAAAAACAGGTTAAACGCTTTCTGAATACACCGGAACTG
AGCATTCTGGGTAGCAATGTTGAATATATCAATGAAGCCAGCGAAAGCATTGGCTATAGCAACTATCCG
CTGGATCATAGCAGCATTGTTAATAGCTTTCCGTTTCGTTGTAATCTGGCACATCCGACCATTATGGTT
AAAAAAGAAGTGATTACCACGCTTGGTGGCTATATGTATGGTAGCCTGAGCGAAGATTATGATCTGTGG
ATTCGTGCAAGCCGTCATGGCAATTTCAAATTTAGCAATATTGATGAACCGCTGCTGAAGTACCGTATT
CATAAAGGTCAGGCAACCAATAAAAGCAACGCCTATAACATCTTTGCCTTTGATAGCAGCCTGAAAATC
CGTGAATTTCTGCTGAATGGTAATGTGCAGTATCTGCTGGGTGCAGCACGTGGTTTTTTTGCATTTCTG
TATGTGCGCTTCATCAAAAAAATGAGGACTTCTACCTGGTCTCCGGCAATTAAAAAAGCGGCTAACCACG
CCGCTTTTTTTACGTCTGCACCTAGACGTACAGTCATCATGCGTACTGACTGACTGCCATGGCATCATG
CTACTAGGGTACGGGTTTTGCTGCCCGCAAACGGGCTGTTCTGGTGTTGCTAGTTTGTTATCAGAATCG
CAGATCCGGCTTCAGCCGGTTTGCCGGCTGAAAGCGCTATTTCTTCCAGAATTGCCATGATTTTTTCCC
CACGGGAGGCGTCACTGGCTCCCGTGTTGTCGGCAGCTTTGATTCGATAAGCAGCATCGCCTGTTTCAG

GCTGTCTATGTGTGACTGTTGAGCTGTAACAAGTTGTCTCAGGTGTTCAATTTCATGTTCTAGTTGCTT
TGTTTTACTGGTTTCACCTGTTCTATTAGGTGTTACATGCTGTTCATCTGTTACATTGTCGATCTGTTC
ATGGTGAACAGCTTTGAATGCACCAAAAACTCGTAAAAGCTCTGATGTATCTATCTTTTTTACACCGTT
TTCATCTGTGCATATGGACAGTTTTCCCTTTGATATGTAACGGTGAACAGTTGTTCTACTTTTGTTTGT
TAGTCTTGATGCTTCACTGATAGATACAAGAGCCATAAGAACCTCAGATCCTTCCGTATTTAGCCAGTA
TGTTCTCTAGTGTGGTTCGTTGTTTTTGCGTGAGCCATGAGAACGAACCATTGAGATCATACTTACTTT
GCATGTCACTCAAAAATTTTGCCTCAAAACTGGTGAGCTGAATTTTTGCAGTTAAAGCATCGTGTAGTG
TTTTTCTTAGTCCGTTATGTAGGTAGGAATCTGATGTAATGGTTGTTGGTATTTTGTCACCATTCATTT
TTATCTGGTTGTTCTCAAGTTCGGTTACGAGATCCATTTGTCTATCTAGTTCAACTTGGAAAATCAACG
TATCAGTCGGGCGGCCTCGCTTATCAACCACCAATTTCATATTGCTGTAAGTGTTTAAATCTTTACTTA
TTGGTTTCAAAACCCATTGGTTAAGCCTTTTAAACTCATGGTAGTTATTTTCAAGCATTAACATGAACT
TAAATTCATCAAGGCTAATCTCTATATTTGCCTTGTGAGTTTTCTTTTGTGTTAGTTCTTTTAATAACC
ACTCATAAATCCTCATAGAGTATTTGTTTTCAAAAGACTTAACATGTTCCAGATTATATTTTATGAATT
TTTTTAACTGGAAAAGATAAGGCAATATCTCTTCACTAAAAACTAATTCTAATTTTTCGCTTGAGAACT
TGGCATAGTTTGTCCACTGGAAAATCTCAAAGCCTTTAACCAAAGGATTCCTGATTTCCACAGTTCTCG
TCATCAGCTCTCTGGTTGCTTTAGCTAATACACCATAAGCATTTTCCCTACTGATGTTCATCATCTGAG
CGTATTGGTTATAAGTGAACGATACCGTCCGTTCTTTCCTTGTAGGGTTTTCAATCGTGGGGTTGAGTA
GTGCCACACAGCATAAAATTAGCTTGGTTTCATGCTCCGTTAAGTCATAGCGACTAATCGCTAGTTCAT
TTGCTTTGAAAACAACTAATTCAGACATACATCTCAATTGGTCTAGGTGATTTTAATCACTATACCAAT
TGAGATGGGCTAGTCAATGATAATTACTAGTCCTTTTCCCGGGTGATCTGGGTATCTGTAAATTCTGCT
AGACCTTTGCTGGAAAACTTGTAAATTCTGCTAGACCCTCTGTAAATTCCGCTAGACCTTTGTGTGTTT
TTTTTGTTTATATTCAAGTGGTTATAATTTATAGAATAAAGAAAGAATAAAAAAAGATAAAAAGAATAG
ATCCCAGCCCTGTGTATAACTCACTACTTTAGTCAGTTCCGCAGTATTACAAAAGGATGTCGCAAACGC
TGTTTGCTCCTCTACAAAACAGACCTTAAAACCCTAAAGGCTTAAGTAGCACCCTCGCAAGCTCGGGCA
AATCGCTGAATATTCCTTTTGTCTCCGACCATCAGGCACCTGAGTCGCTGTCTTTTTCGTGACATTCAG
TTCGCTGCGCTCACGGCTCTGGCAGTGAATGGGGGTAAATGGCACTACAGGCGCCTTTTATGGATTCAT
GCAAGGAAACTACCCATAATACAAGAAAAGCCCGTCACGGGCTTCTCAGGGCGTTTTATGGCGGGTCTG
CTATGTGGTGCTATCTGACTTTTTGCTGTTCAGCAGTTCCTGCCCTCTGATTTTCCAGTCTGACCACTT
CGGATTATCCCGTGACAGGTCATTCAGACTGGCTAATGCACCCAGTAAGGCAGCGGTATCATCAACAGG
CTTACCCGTCTTACTGTCCCTAGTGCTTGGATTCTCACCAATAAAAAACGCCCGGCGGCAACCGAGCGT
TCTGAACAAATCCAGATGGAGTTCTGAGGTCATTACTGGATCTATCAACAGGAGTCCAAGC

### Terminators

TrrnB

GAAGCTTGGGCCCGAACAAAAACTCatctcagaagaggatctgaatagcgccgtcgaccatcatcatca
tcatcattgagtttaaacggtctccagcttggctgttttggcggatgagagaagattttcagcctgata
cagattaaatcagaacgcagaagcggtctgataaaacagaatttgcctggcggcagtagcgcggtggtc
ccacctgaccccatgccgaactcagaagtgaaacgccgtagcgccgatggtagtgtggggtctccccat
gcgagagtagggaactgccaggcatcaaataaaacgaaaggctcagtcgaaagactGGGCCTTTCGTTT
TATCTGTTGTTTGTCGGTGAACT

BBa_B0015

ccaggcatcaaataaaacgaaaggctcagtcgaaagactgggcctttcgttttatctgttgtttgtcgg
tgaacgctctctactagagtcacactggctcaccttcgggtgggcctttctgcgtttata

BBa_B1002

cgcaaaaaaccccgcttcggcggggttttttcgc

ECK120033736[7]

aacgcatgagAAAGCCCCCGGAAGATCACCTTCCGGGGGCTTTtttattgcgc

ECK120033737[Z]
ggaaacacagAAAAAAGCCCGCACCTGACAGTGCGGGCTTTTTTTTTcgaccaaagg

ECK120010818[Z]
GTCAGTTTCACCTGTTTTACGTAAAAACCCGCTTCGGCGGGTTTTTACTTTTGG

ECK120015440[Z]
tccggcaattAAAAAAGCGGCTAACCACGCCGCTTTTTTtacgtctgca
